# Domain-swapped LuxR-type quorum sensing receptors reveal divergent ligand-response mechanisms among homologs

**DOI:** 10.64898/2026.04.27.721074

**Authors:** Irene M. Stoutland, Susan A. Walker, Helen E. Blackwell

**Affiliations:** Department of Chemistry, University of Wisconsin–Madison, 1101 University Ave., Madison, WI, 53706 USA

**Keywords:** bacteria, cell-cell signaling, chemical biology, dissociative mechanism, quorum sensing, signal transduction, transcription factor

## Abstract

Many common bacteria use quorum sensing to regulate cell density-dependent phenotypes, including luminescence, biofilm formation, virulence, and symbiosis. The LuxI/R system is the best-characterized quorum sensing pathway in Gram-negative bacteria and consists of a LuxI-type synthase that produces an *N*-acyl L-homoserine lactone (AHL) autoinducer and a LuxR-type transcription factor that is regulated by AHL binding. Binding of native AHL signal promotes DNA binding and transcriptional regulation in some LuxR homologs (associative-type), while other homologs regulate transcription in the absence of ligand and are inactivated by native signal binding (dissociative-type). To better characterize what features determine ligand-response type, we generated structural mutants of two associative receptors (LasR of *Pseudomonas aeruginosa* and MrtR of *Mesorhizobium tianshanense*) and two dissociative receptors (EsaR of *Pantoea stewartii* and ExpR2 of *Pectobacterium versatile*). Swapping domains between these receptors revealed that the ligand-binding domain primarily determines associative vs. dissociative activity in response to native AHL agonists. Further, non-native AHL-derived antagonists maintained their activity profiles in receptors with interchanged DNA-binding domains. We also found that the extended linker between domains observed in the dissociative receptors does not determine mechanism of ligand response, and that inter-domain interactions may play an important role in activation for some receptors but not others. Notably, deletion of just one residue from the dissociative receptor EsaR produced a mutant with associative activity, the first time such mechanism switching has been reported for a LuxR-type receptor. These findings illuminate features essential for ligand response and highlight the mechanistic diversity of the LuxR family.

**IMPORTANCE:** LuxI/R quorum sensing regulates various cell density-dependent phenotypes in Gram-negative bacteria. Prior research has developed small molecule modulators of LuxR-type receptors, with potential applications in anti-virulence, anti-biofouling, and bioengineering. Competitive antagonists have been reported for receptors active in the presence of native ligand but not for receptors active in its absence. A lack of knowledge about the molecular mechanisms of receptor response to ligand limits both our fundamental understanding of the LuxI/R quorum sensing process and the rational design of chemical modulators with superior activity profiles. We used a structural mutagenesis strategy with four LuxR-type receptors that operate via two distinct mechanisms to begin to dissect the structural features that drive differences in ligand response between receptors. These insights could aid in efforts to characterize novel LuxR homologs, understand potential interspecies communication via quorum sensing, and develop improved chemical probes to alter LuxR-type receptor activity.

## INTRODUCTION

Quorum sensing (QS) is a cell-cell signaling pathway used by common bacteria to regulate population density-dependent phenotypes, including virulence factor production, motility, biofilm formation, symbioses, bioluminescence, and natural product production (1, 2). Many Gram-negative proteobacteria encode one or more LuxI/R QS systems, consisting of a LuxI-type synthase that produces an *N*-acyl L-homoserine lactone (AHL) signaling molecule and a cytoplasmic LuxR-type transcription factor regulated by AHL binding (3, 4). AHL concentration increases with cell density, and once a threshold AHL concentration is achieved in the local environment, AHL:LuxR-receptor binding alters the expression of QS-dependent genes (shown schematically in **Figure 1**).

**Figure 1.**
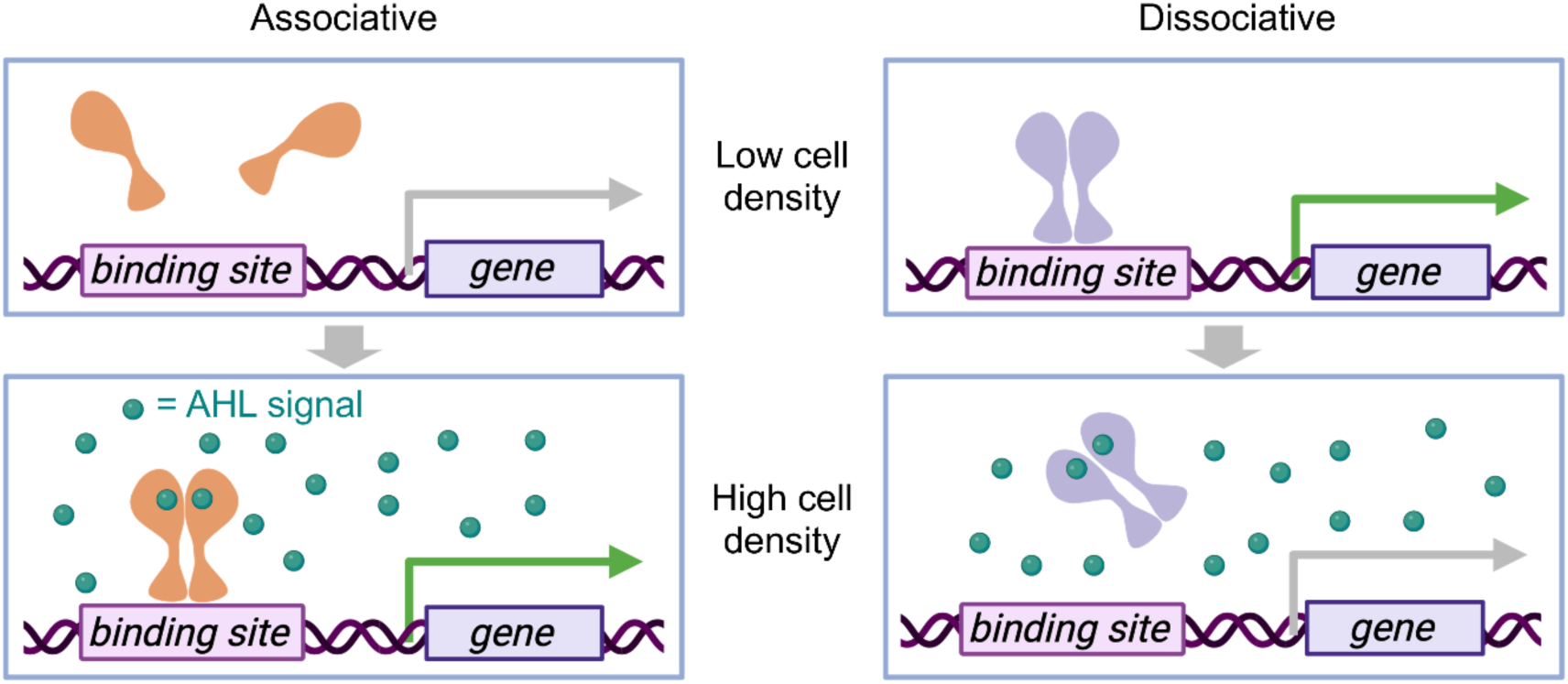
LuxR-type QS receptor mechanisms. Associative LuxR-type receptors (*left*) are activated by AHL binding. The AHL:receptor complex dimerizes and binds to DNA to regulate transcription. In contrast, dissociative LuxR-type receptors (*right*) regulate transcription in the absence of AHL and are inactivated upon AHL binding.

Of the thousands of putative LuxR-type receptors found in bacteria (5), only ∼10 have been biochemically and structurally characterized (6–12). These LuxR-type receptors are unified by their similarity in fold but differ greatly in amino acid sequence, stability, and mode of ligand response. LuxR homologs share low (<25%) sequence identity and typically consist of a ∼170 residue N-terminal ligand-binding domain (LBD) and a ∼60 residue C-terminal helix-turn-helix DNA-binding domain (DBD) connected by a ∼10-20 residue linker. In most well-studied LuxR homologs, ligand binding increases DNA binding and transcriptional regulation (i.e., an associative mechanism, **Figure 1**, left). Members of a less-studied group of LuxR-type receptors activate or repress transcription in the absence of ligand and are inactivated by ligand binding (i.e., a dissociative mechanism, **Figure 1**, right).

Strategies to modulate LuxI/R-type QS systems are of interest for applications in anti-virulence, anti-biofouling, synthetic biology, and bioengineering (13–15), and small molecules that activate or inhibit LuxR-type receptors represent an established strategy to influence bacterial phenotypes and probe mechanistic questions about these QS systems (13, 16–21). Despite broad interest in modulating and engineering LuxI/R-type systems, the mechanisms by which ligand binding affects LuxR-type receptor activity are poorly understood. Associative and dissociative receptors do not appear to favor distinct AHL signals. In fact, many AHLs are shared across receptors of both types; for example, LuxR of *Vibrio fischeri* (associative), ExpR2 of *Pectobacterium versatile* (previously *Pectobacterium carotovorum*) (dissociative), and EsaR of *Pantoea stewartii* (dissociative) all respond to the same native AHL (22, 23).

Competitive antagonists (i.e., compounds with activity that opposes that of the native ligand) have been reported for numerous associative receptors (16, 18, 20, 24) but not for dissociative receptors. For example, in a recent screen of ∼100 AHLs and AHL analogs, including many known agonists and antagonists of associative LuxR-type receptors, no compounds antagonized dissociative LuxR-type receptor EsaR (25, 26). Whether or not dissociative receptors are generally harder to antagonize than their associative counterparts remains to be seen. Of the known competitive antagonists for associative LuxR-type receptors, most have only moderate potencies (low μM IC_50_ values in cells), and extensive structural alterations have not revealed insights into routes to potency improvement (14, 27). A better understanding of the molecular mechanisms of ligand response among LuxR homologs, including differences between associative and dissociative receptors, could support the rational design of chemical probes with improved potencies and specificities, and may lead to the development of antagonists targeting dissociative LuxR-type receptors.

The low in vitro solubility of many LuxR-type receptors, especially in the absence of AHL, hinders biochemical and structural approaches to directly determine the molecular mechanisms of ligand response in associative vs. dissociative receptors (8, 10), but mutational analysis can yield informative clues (28, 29). Interestingly, dissociative receptors tend to have longer linkers between the LBD and DBD (∼9 vs. ∼16 residues) and a shorter N-terminus than associative receptors (by ∼4-15 residues) (**Figure S1**), but the functional importance of these regions and their relationship to receptor mechanism is poorly understood. Although several studies have examined LuxR domain truncations (29–31), investigations into the role of the linker in LuxR-type receptor activity are limited (28), which is relatively surprising given its role in connecting the two functional domains (LBD and DBD) of this protein class. Of the few structures of full-length LuxR-type receptors reported (6–12), all act via an associative mechanism. From these structures, it is obvious that the LBD and DBD can engage in distinct intra– and interprotein interactions; the length and conformation of the linker domain could play a role in positioning these domains to generate a functional dimer.

In this study, we decipher characteristics of the LBD, DBD, and linker region that govern ligand response and activation in associative vs. dissociative LuxR-type receptors. Our investigations focused on two associative receptors, LasR of *Pseudomonas aeruginosa* (8) and MrtR of *Mesorhizobium tianshanense* (32), and two dissociative receptors, EsaR of *P. stewartii* (33) and ExpR2 of *P. versatile* (25), as their mechanistic classes have been well validated (25, 26, 28, 29, 32). The AlphaFold 3 (34) or X-ray crystal structures of these receptors are shown in **Figure 2**, highlighting features of interest examined in this study. Our results establish that LuxR homologs differ markedly in their tolerance of domain swapping and linker perturbations, refining the mechanistic understanding of this receptor family.

**Figure 2.**
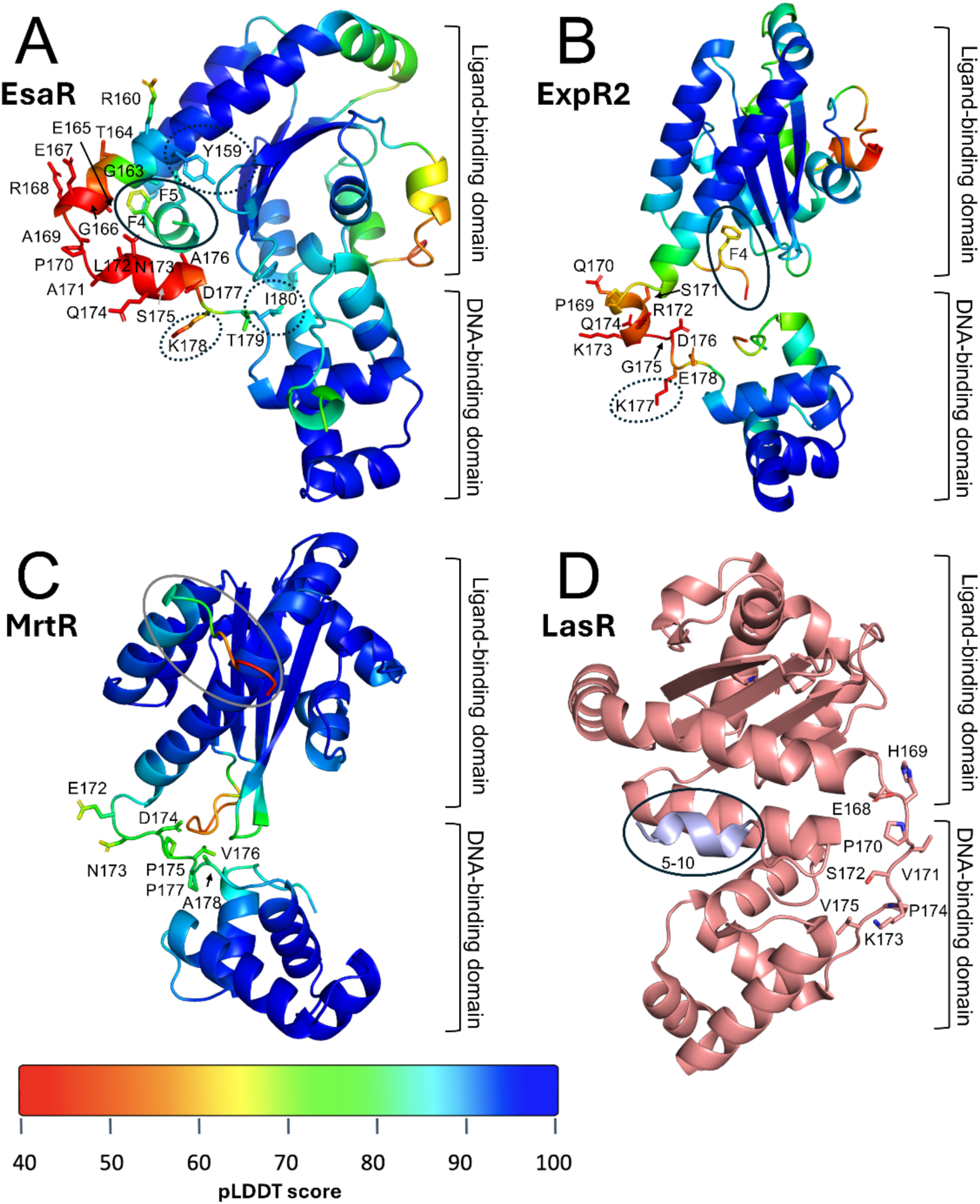
Structures of LuxR-type proteins in this study. AlphaFold 3-predicted structures of EsaR **(A)**, ExpR2 **(B)**, and MrtR **(C)** and crystal structure of LasR (PDB ID: 6V7X) (32) **(D)**, highlighting features of interest. AlphaFold structures are colored by prediction confidence as indicated by pLDDT scale shown (>90 = very high confidence, 70-90 = confident, 50-70 low confidence, <50 = very low confidence). The LBD and DBD are indicated at right of each panel. Residues around the linker region are labeled, and the far N-terminus is circled. The extended N-terminus observed in LasR is colored in violet (residues 5-10). Residues at which point mutations were found to have a large impact on activity are indicated by dashed circles.

## RESULTS

### Ligand-binding domain determines associative or dissociative activity profile

Domain swapping has been applied to various transcription factor families, often to develop synthetic biology tools (33). Chimeric LuxR-type receptors have been used to tune ligand and promoter preferences in engineered regulators (34, 35), activate biosynthetic gene clusters (36), and investigate differences between an associative and a ligand-unresponsive receptor (37). Here, we report the first domain swapping between associative and dissociative LuxR-type receptors. Chimeras were designed by fusing the LBD of an associative receptor (LasR or MrtR) to the DBD of a dissociative receptor (EsaR or ExpR) and vice versa (chimera compositions listed in **Table 1**). A total of 13 EsaR-LasR chimeras were created to test the effect of junction point on activity, and 1-4 chimeras were created for each of the other receptor pairs to explore the mechanistic effects of different receptor combinations. Sequence alignments (**Figure S1**) and structural models (shown in **Figure 2**) were used to guide junction point selections.

**Table 1.**
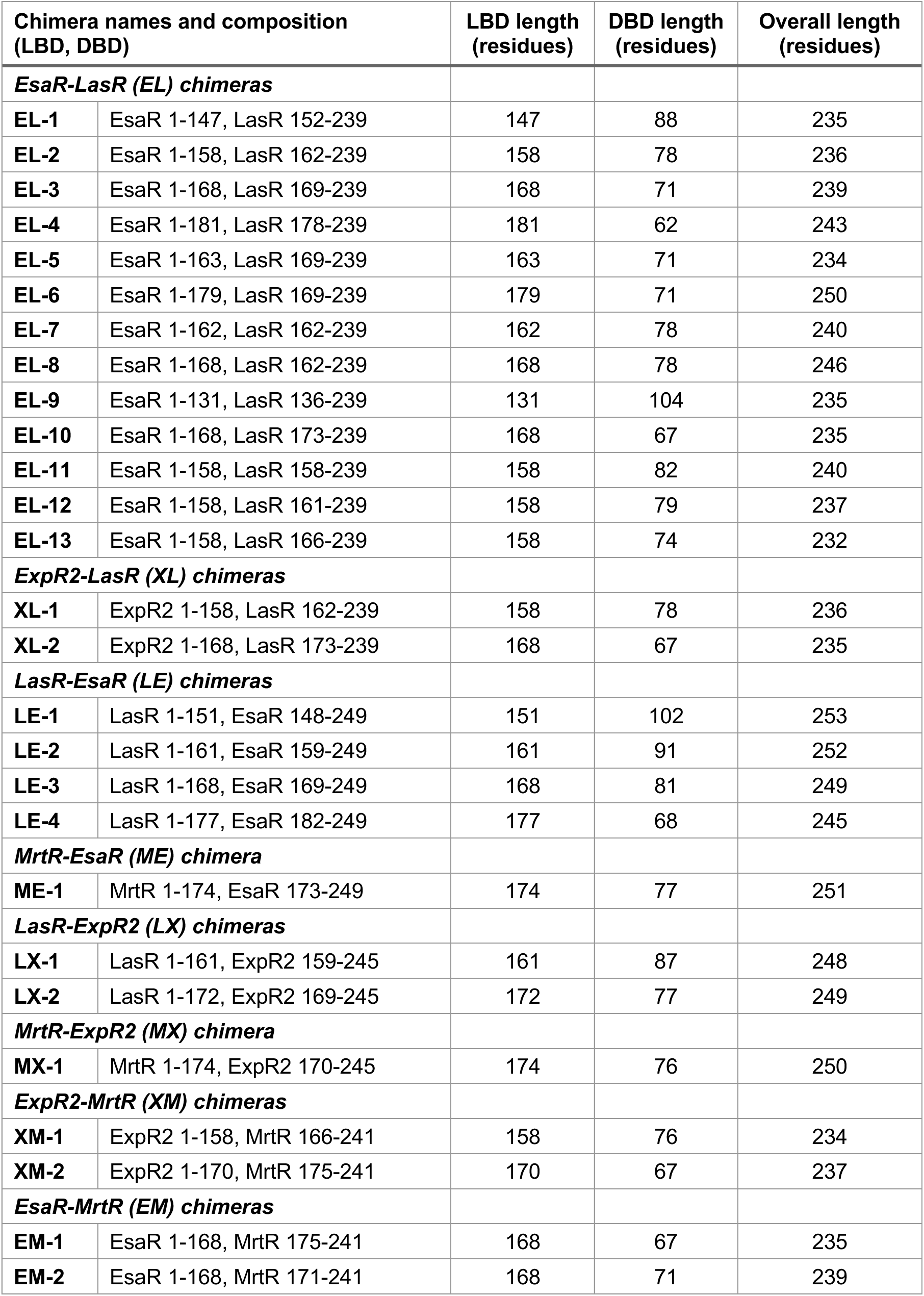
Compositions of LuxR chimeras evaluated in this study. Activity data in reporter assays are shown in **Figures 3** and **S4**. See **Table S1** for WT protein sequences.

| Chimera names and composition (LBD, DBD) |  | LBD length (residues) | DBD length (residues) | Overall length (residues) |
| --- | --- | --- | --- | --- |
| ***EsaR-LasR (EL) chimeras*** |  |  |  |  |
| **EL-1** | EsaR 1-147, LasR 152-239 | 147 | 88 | 235 |
| **EL-2** | EsaR 1-158, LasR 162-239 | 158 | 78 | 236 |
| **EL-3** | EsaR 1-168, LasR 169-239 | 168 | 71 | 239 |
| **EL-4** | EsaR 1-181, LasR 178-239 | 181 | 62 | 243 |
| **EL-5** | EsaR 1-163, LasR 169-239 | 163 | 71 | 234 |
| **EL-6** | EsaR 1-179, LasR 169-239 | 179 | 71 | 250 |
| **EL-7** | EsaR 1-162, LasR 162-239 | 162 | 78 | 240 |
| **EL-8** | EsaR 1-168, LasR 162-239 | 168 | 78 | 246 |
| **EL-9** | EsaR 1-131, LasR 136-239 | 131 | 104 | 235 |
| **EL-10** | EsaR 1-168, LasR 173-239 | 168 | 67 | 235 |
| **EL-11** | EsaR 1-158, LasR 158-239 | 158 | 82 | 240 |
| **EL-12** | EsaR 1-158, LasR 161-239 | 158 | 79 | 237 |
| **EL-13** | EsaR 1-158, LasR 166-239 | 158 | 74 | 232 |
| ***ExpR2-LasR (XL) chimeras*** |  |  |  |  |
| **XL-1** | ExpR2 1-158, LasR 162-239 | 158 | 78 | 236 |
| **XL-2** | ExpR2 1-168, LasR 173-239 | 168 | 67 | 235 |
| ***LasR-EsaR (LE) chimeras*** |  |  |  |  |
| **LE-1** | LasR 1-151, EsaR 148-249 | 151 | 102 | 253 |
| **LE-2** | LasR 1-161, EsaR 159-249 | 161 | 91 | 252 |
| **LE-3** | LasR 1-168, EsaR 169-249 | 168 | 81 | 249 |
| **LE-4** | LasR 1-177, EsaR 182-249 | 177 | 68 | 245 |
| ***MrtR-EsaR (ME) chimera*** |  |  |  |  |
| **ME-1** | MrtR 1-174, EsaR 173-249 | 174 | 77 | 251 |
| ***LasR-ExpR2 (LX) chimeras*** |  |  |  |  |
| **LX-1** | LasR 1-161, ExpR2 159-245 | 161 | 87 | 248 |
| **LX-2** | LasR 1-172, ExpR2 169-245 | 172 | 77 | 249 |
| ***MrtR-ExpR2 (MX) chimera*** |  |  |  |  |
| **MX-1** | MrtR 1-174, ExpR2 170-245 | 174 | 76 | 250 |
| ***ExpR2-MrtR (XM) chimeras*** |  |  |  |  |
| **XM-1** | ExpR2 1-158, MrtR 166-241 | 158 | 76 | 234 |
| **XM-2** | ExpR2 1-170, MrtR 175-241 | 170 | 67 | 237 |
| ***EsaR-MrtR (EM) chimeras*** |  |  |  |  |
| **EM-1** | EsaR 1-168, MrtR 175-241 | 168 | 67 | 235 |
| **EM-2** | EsaR 1-168, MrtR 171-241 | 168 | 71 | 239 |

The chimeras were tested for their ability to regulate transcription in *E. coli* reporter systems containing a protein production plasmid and a reporter plasmid with a LuxR-regulated promoter upstream of either GFP or β-galactosidase (see **Table S2** for strains and plasmids). The translational start site of MrtR is unclear (38), and for this study, we used the start site that was assigned to CinR of *Rhizobium etli* (96% identical to MrtR, **Figure S2**) based on codon usage, GC content, and similarity to known genes (39). As each of the four WT receptors examined has a different promoter specificity, chimeras were tested for their ability to regulate transcription at the promoter corresponding to the DBD, and their activity was compared to that of the WT receptor with the same DBD. EsaR has been shown to activate some promoters and repress others (28), and chimeras with an EsaR DBD were tested at both an EsaR-activated and an EsaR-repressed promoter.

With regard to AHL ligands, the native AHL for LasR (29) (*N*-(3-oxo)-dodecanoyl-L-homoserine lactone, 3OC12 hereafter) and the native AHL for EsaR (23) and ExpR2 (22) (*N-*(3-oxo)-hexanoyl-L-homoserine lactone, 3OC6 hereafter) are well established (structures shown in **Figure 3A**). For MrtR, the native AHL is less established, but the AHL synthase of *M. tianshanense*, MrtI, is 98% identical to CinI of *Rhizobium leguminosaurum* (**Figure S2**), which produces *N*-(3*R*-hydroxy)-7-*cis*-tetradecenoyl-L-homoserine lactone (7*Z*-3*R*-OHC14 hereafter) (40–42). In addition, mass spectrometry data suggest that the major AHL produced by *M. tianshanense* has a molecular weight of 325.2 g/mol, which could correspond to either 3OC14 or 3OHC14 with an unsaturated lipid tail (43). Together, these data suggest that the major AHL produced by *M. tianshanense* is likely 7*Z*-3R-OHC14. We found that MrtR was strongly activated by 7*Z*-3OHC14 (mix of 3*R* and 3*S* diastereomers) in an *E. coli* reporter and therefore used this AHL as the native ligand to test MrtR mutants and chimeras with an MrtR LBD. Selected chimeras were tested for their ability to respond to the native AHL sensed by the LBD receptor vs. the native AHL sensed by the DBD receptor, and all responded more strongly to the ligand corresponding to the LBD receptor (**Figure S3**), as previously observed for other chimeras of LuxR-type receptors (34). Thus, the native AHL corresponding to the LBD receptor was used to screen chimera activity at concentrations sufficient to induce maximum activity from the WT receptors.

**Figure 3.**
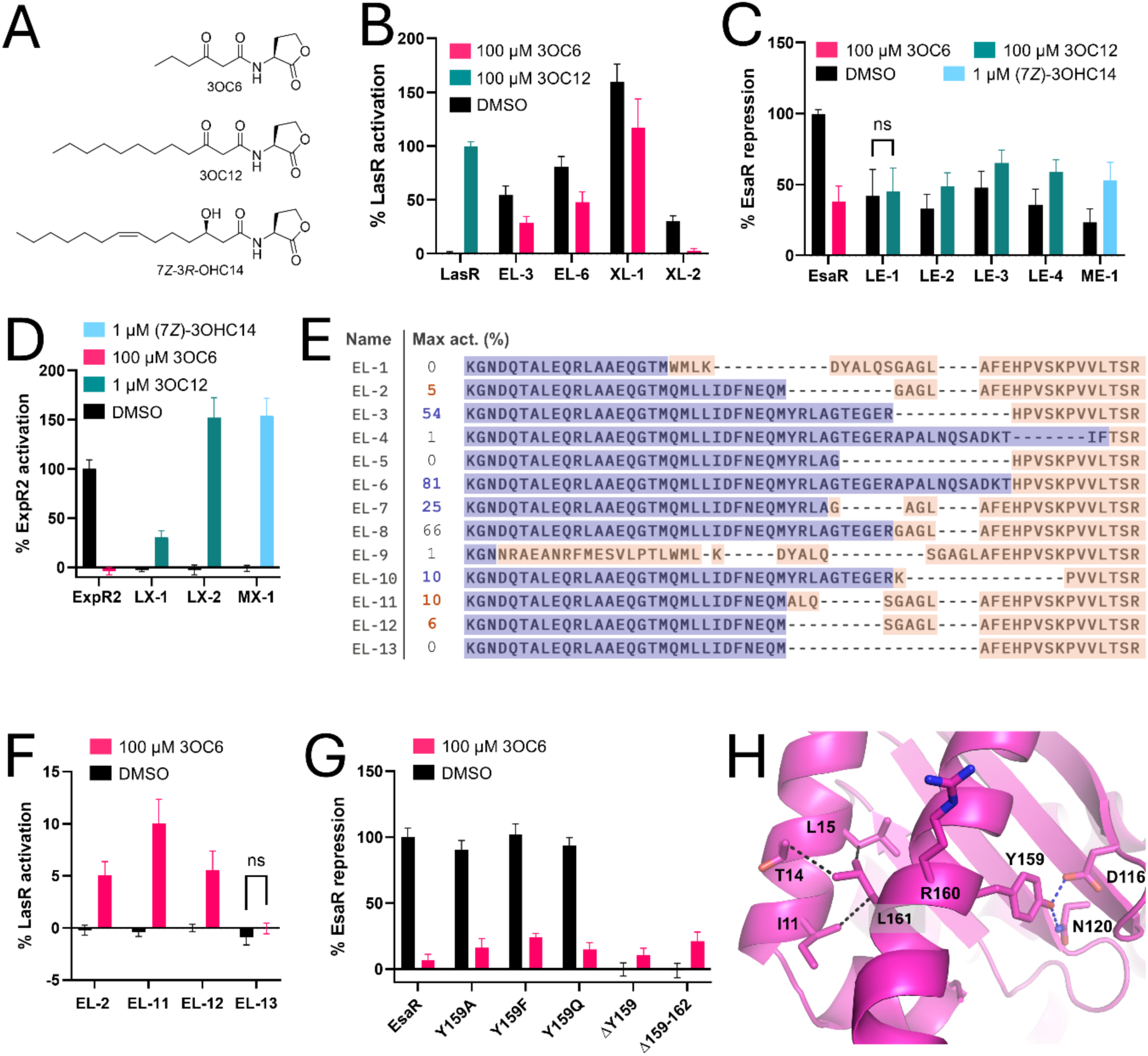
Transcriptional regulation by chimeras of associative and dissociative LuxR-type receptors measured in *E. coli* reporters. For chimera names and compositions, see Table 1. **(A)** Structures of native AHL signals for LasR, EsaR, ExpR2, and MrtR. **(B)** Transcriptional activation of the *lasI* promoter by representative chimeras with a LasR DBD in an *E. coli* JLD271 reporter, normalized to uninduced culture (0%) and LasR saturated with 3OC12 (100%). WT LasR activity is shown for reference. Chimeras with a LasR DBD that showed low activity or no ligand response are shown in **Figure S4C**. **(C)** Transcriptional repression of the *esaR* promoter by chimeras with an EsaR DBD in a JLD271 reporter, normalized to uninduced culture (0%) and maximum EsaR repression (100%). WT EsaR activity is shown for reference. Activity of the same chimeras in an EsaR-activated reporter is shown in **Figure S4B**. **(D)** Transcriptional activation of the *rsmA* promoter by chimeras with an ExpR2 DBD in a JLD271 reporter, normalized to uninduced culture (0%) and maximum ExpR2 activation (100%). WT ExpR2 activity is shown for reference. **(E)** Multiple sequence alignment of the junction points of EsaR-LasR chimeras. Maximum activity of each chimera relative to WT LasR is indicated and colored based on whether activity was higher with DMSO (dissociative, purple), with 3OC6 (associative, orange), or not significantly different with and without ligand (black). Sequence is colored based on whether it originated from EsaR (purple) or LasR (orange). **(F)** Transcriptional activation of the *lasI* promoter by chimeras with an EsaR LBD (residues 1-158) and a LasR DBD in a JLD271 reporter, normalized to uninduced culture (0%) and LasR saturated with 3OC12 (100%). **(G)** Transcriptional repression of the *esaR* promoter by EsaR mutants in an *E. coli* BW27749 reporter, normalized to uninduced culture (0%) and maximum EsaR repression (100%). WT EsaR activity is shown for reference. Activity of the same mutants in an EsaR-activated reporter is shown in **Figure S4D**. **(H)** Section of an AlphaFold 3-predicted structure of EsaR. Blue dashed lines indicate potential hydrogen bonds to Y159 (<3 Å), and black dashed lines indicate potential hydrophobic interactions involving L161 (<4.5 Å). **(All graphs)** Error bars represent standard deviation. Unless indicated as “ns,” activity was significantly different with and without ligand based on multiple t-tests with the Holm-Šídák correction for multiple comparisons and p ≤ 0.01. See **Table S2** for full sequence details. All graphs represent at least 3 biological replicates, each performed in technical triplicate.

Overall, we found that the mechanism of the LBD receptor determined whether a chimera showed higher activity in the presence of ligand (associative) or in the absence of ligand (dissociative) (**Figure 3B-D**). Active chimeras with the LBD of a dissociative receptor (EsaR or ExpR2) and the DBD of the associative receptor LasR were more active in the absence of ligand than in its presence (i.e., a dissociative profile for EL and XL chimeras, **Figure 3B**), with a few notable exceptions, discussed below. In contrast, active chimeras with the LBD of an associative receptor (LasR or MrtR) and the DBD of a dissociative receptor (EsaR or ExpR2) were more active in the presence of ligand than in its absence (i.e., an associative profile for LM, LX, ME, and MX chimeras, **Figure 3C-D**). All chimeras with the MrtR DBD were inactive (**Figure S4A**). It is possible that the MrtR DBD requires specific contacts with the LBD that are lost in the chimeras, as proposed for poorly functional chimeras from other protein families (33, 44). Chimeras with an EsaR DBD were unable to activate transcription from an EsaR-activated promoter (**Figure S4B**) but could weakly repress transcription at an EsaR-repressed promoter (**Figure 3C**), suggesting that these chimeras may be deficient for interactions with RNA polymerase (RNAP). MrtR-EsaR and LasR-EsaR constructs showed greater repression of transcription in the presence of ligand than in its absence (an associative profile, matching the LBD).

Based on western blots of a selection of chimera reporter strains, a lack of protein accumulation did not explain the low activity of LasR-EsaR chimeras, and the relative accumulation of chimeras in the presence and absence of ligand correlated with mechanism—LasR and associative chimeras showed increased accumulation with ligand, while EsaR and dissociative chimeras did not (**Figure S5A-E**; see SI for further discussion). In addition, the activity of chimeras with LasR, ExpR2, or EsaR DBDs was not due to constitutive activity of the DBDs alone (**Figure S6A** and **S6B**; see SI for further discussion).

### Junction point affects chimera activity and mechanism

We next constructed 13 EsaR-LasR chimeras to investigate the effect of the junction point on transcriptional activation (**Figure 3E**) and found that the choice of LBD and DBD segment had a large effect on activity, congruent with previous studies of other LuxR-type receptor chimeras (35, 36). Chimeras with the shortest N-terminal domains (EL-1, EL-9), shortest DBDs (EL-4, EL-10), shortest length overall (EL-13) showed little or no activity. Among chimeras with the same DBD, a longer LBD led to higher activity (EL6 > EL-3 > EL-5; EL-8 > EL-7 > EL-2), and among those with the same LBD, a longer DBD tended to lead to higher activity (EL-8 > EL-3 > EL-10; EL-11 > EL-12 = EL-2 > EL-13), suggesting that too short a linker between domains is detrimental to activity.

In a few specific cases, altering the junction point reversed chimera mechanism. While most of the EsaR-LasR chimeras showed dissociative activity, like WT EsaR, EL-2 was inactive in the absence of ligand yet weakly active (∼5%) with 100 µM 3OC6 (**Figure 3F**). In contrast to EL-3, which showed an EC_50_ of ∼240 nM for 3OC6 (comparable to its previously reported EC_50_ for EsaR) (25, 26), EL-2 had an EC_50_ >100 µM and reached a maximum of ∼30% LasR activity at 1 mM 3OC6 (**Figures S6C** and **S6D**). In addition, EL-2 was the only EsaR-LasR chimera that showed higher accumulation in the presence of ligand, as determined via western blot (**Figure S5E**), which may explain its associative activity. Changing the length of the DBD (−1 or +4 residues) produced chimeras that showed the same weakly associative activity as EL-2 (i.e., chimeras EL-11 and EL-12, **Figure 3F**), suggesting that the choice of LBD segment is responsible for the associative activity. Interestingly, the EL-2 LBD segment only differs from EL-7, which shows the expected dissociative activity, by four residues at the end of the EsaR LBD segment (EsaR 159-162), suggesting that these residues in the LBD play an important role in determining associative vs. dissociative activity. Single point mutations (Y159A, Y159F, or Y159Q) in EsaR substantially decreased activation but had little effect on repression, and deletion of Y159 or residues 159-162 rendered EsaR unable to activate transcription but, surprisingly, more able to repress transcription in the *presence* of ligand than in its absence (**Figures 3G** and **S4D**). These results support that deletion of Y159 converts EsaR from a dissociative to a weakly associative receptor. This finding is significant, as it unveils potential structural contacts important to EsaR’s dissociative mechanism.

Based on an AlphaFold 3 prediction of the EsaR structure, Y159 appears to form a hydrogen bond with D116 and/or N120, located on the β-strands forming the ligand binding pocket, and L161 engages in hydrophobic interactions near the N-terminus (**Figure 3H**). The fact that deletion of Y159 or residues 159-162, but not point mutations of Y159, reversed receptor mechanism suggests that perturbations of nearby interactions in the LBD, not just the loss of contacts to Y159 itself, are responsible for the dissociative to associative mechanistic reversal. Ligand binding has been shown to stabilize EsaR against in vitro proteolysis (23), and, in our hands, purified EsaR is more soluble with 3OC6 than without (data not shown). Accordingly, the *loss* of interactions that stabilize EsaR in the absence of ligand may ablate activity in the absence of ligand, and ligand-induced stabilization of ΔY159 and Δ159-162 thus may account for the higher activity of these mutants in the presence of ligand.

### Divergent responses to linker mutations indicate differences in molecular mechanisms

As introduced above, the linker region between the LBD and DBD varies across LuxR-type receptors. Dissociative receptors tend to have longer linkers than associative receptors by ∼7 residues (4) (**Figure S1**), yet it is unclear if this difference in length contributes to differences in receptor mechanism. To probe the role of this region in receptor function, we generated mutants of LasR, MrtR, EsaR, and ExpR2 with shortened or lengthened linkers.

First, we tested the effect of linker deletions on receptor activity using reporter assays in *E. coli,* as described above. Compared to EsaR, the linkers of ExpR2, LasR, and MrtR are approximately 1, 7, and 7 residues shorter, respectively (see sequence alignment in **Figure S1**). Receptors with short linkers showed a low tolerance for linker deletions: LasR and MrtR mutants with 3 residues deleted from the linker were <10% active (**Figure 4A**, **B**). In contrast, receptors with longer linkers were more tolerant to linker deletions: EsaR remained highly active with up to 8 residues deleted from the linker (**Figure 4C**), and ExpR2 maintained moderate activity with up to 6 residues deleted (**Figure 4D**). These data suggest that, at least within this set of receptors, the minimum functional linker length may be similar for associative and dissociative receptors, rather than dissociative receptors requiring an extended linker for function.

**Figure 4.**
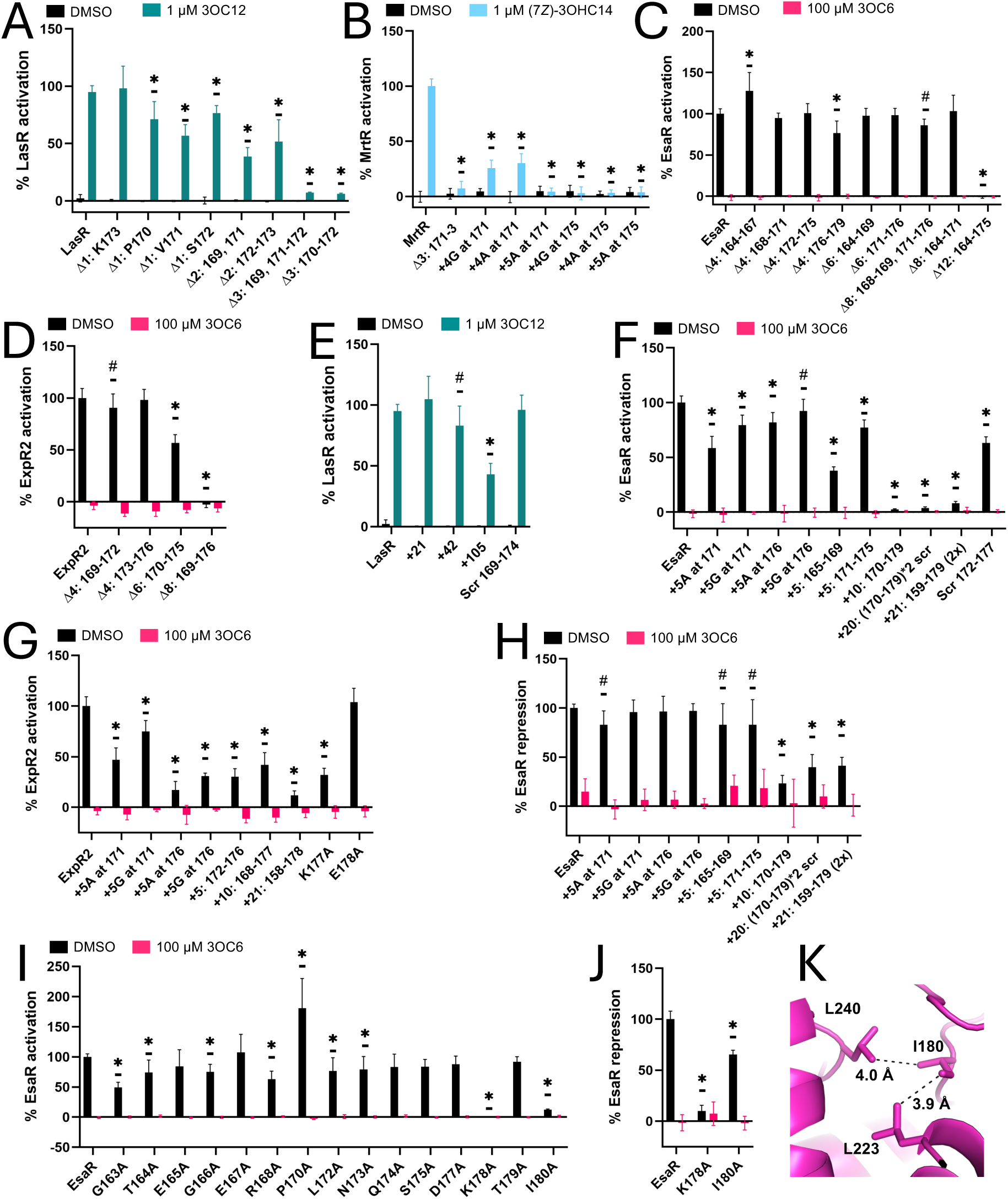
Activities of LasR, EsaR, ExpR2, and MrtR linker mutants in *E. coli* reporter assays. **(A)** Transcriptional activation of the *lasI* promoter by LasR mutants in *E. coli* BW27749, normalized to uninduced culture (0%) and LasR saturated with 3OC12 (100%). **(B)** Transcriptional activation of the *mrtI* promoter by MrtR mutants in BW27749, normalized to uninduced culture (0%) and MrtR saturated with 7Z-3OHC14 (100%). **(C)** Transcriptional activation of the *esaS* promoter by EsaR mutants in BW27749, normalized to uninduced culture (0%) and maximum EsaR activation (100%). **(D)** Transcriptional activation of the *rsmA* promoter by ExpR2 mutants in *E. coli* JLD271, normalized to uninduced culture (0%) and maximum ExpR2 activation (100%). **(E)** Transcriptional activation of the *lasI* promoter by LasR mutants in BW27749, normalized to uninduced culture (0%) and LasR saturated with 3OC12 (100%). The number of inserted residues (derived from the EsaR linker) is indicated. **(F)** Transcriptional activation of the *esaS* promoter by EsaR mutants in BW27749, normalized to uninduced culture (0%) and maximum EsaR activation (100%). **(G)** Transcriptional activation of the *rsmA* promoter by ExpR2 mutants in JLD271, normalized to uninduced culture (0%) and maximum ExpR2 activation (100%). **(H)** Transcriptional repression of the *esaR* promoter by EsaR mutants in BW27749, normalized to uninduced culture (0%) and maximum EsaR repression (100%). **(I)** Transcriptional activation of the *esaS* promoter by EsaR mutants in BW27749, normalized to uninduced culture (0%) and maximum EsaR activation (100%). **(J)** Transcriptional repression of the *esaR* promoter by EsaR mutants in BW27749, normalized to uninduced culture (0%) and maximum EsaR repression (100%). **(K)** Section of an AlphaFold 3-predicted structure of EsaR, showing potential interactions between I180 and other residues in the DBD. Dashed lines indicate potential hydrophobic interactions involving I180, with distances indicated. **(All graphs)** Labels indicate the number and identity of residues added or removed. Addition of 5 alanine or glycine residues is indicated by “+5A” and “+5G”, respectively. “Scr” means the indicated sequence was scrambled. In cases where insertions represent a duplication, indicated by “2x,” duplicated residues are indicated. A 2-way ANOVA was used to compare activity across mutants with the Dunnett correction for multiple comparisons. For MrtR and LasR mutants, activity in the absence of compound did not differ from WT. For EsaR and ExpR2 mutants, activity in the presence of 3OC6 did not differ from WT. Asterisks and hash marks indicate that maximum mutant activity differed from WT with p ≤ 0.01 or p ≤ 0.05, respectively. All graphs represent at least 3 biological replicates, each performed in triplicate. See **Table S2** for full sequence details.

Although the extended linker may not be crucial for EsaR’s transcriptional activation or dissociative mechanism, it appears to affect ligand sensitivity. In transcriptional reporter assays, EsaR linker deletion mutants showed 3– to 16-fold increases in EC_50_ for native ligand 3OC6 relative to EsaR, while ExpR2 linker deletions showed slight *decreases* in EC_50_ for native ligand 3OC6 relative to ExpR2 (**Figure S7**), implying that the extended linker may have different roles in ligand sensitivity for ExpR2 vs. EsaR.

Second, we tested the effect of linker extensions on receptor activity in *E. coli* reporter assays. LasR tolerated large linker insertions: for example, adding 21 residues from the EsaR linker to the LasR linker did not decrease LasR activation. Even a LasR mutant with 105 amino acids added to the linker (five repeats of the same 21-residue segment, scrambled) maintained ∼40% WT LasR activation (**Figure 4E**). In contrast, EsaR did not tolerate large linker insertions: duplicating 21 residues of the EsaR linker led to a >90% decrease in transcriptional activation, as did a smaller 10-residue duplication or the addition of a scrambled 20-residue sequence (**Figure 4F**). These results suggest that the linker may influence specific inter-domain contacts in EsaR but not in LasR. Consistent with this hypothesis, scrambling residues 169-174 of the LasR linker had no significant effect on activation, while scrambling EsaR residues 172-177 led to a 37% decrease in activation, (**Figure 4E**, **F**). Like EsaR, ExpR2 was intolerant to long linker insertions (**Figure 4G**).

Western blots of EsaR-activated reporter strains for select linker mutants showed substantial accumulation for 5, 10, 20, and 21-residue additions, suggesting that differences in activation were not caused by differences in protein accumulation (**Figure S8**). Linker insertions caused a smaller effect on EsaR transcriptional repression than on activation (**Figure 4H**). The difference in linker mutant activity between the EsaR-activated and EsaR-repressed reporters could be due to differences in expression levels between the two reporters or due to the inability of linker extension mutants to engage in interactions with RNAP that are required for transcriptional activation but not for repression (4).

Third, we looked at the effects of inserting short sequences of repeating residues into the linker for EsaR, ExpR2, and MrtR. Transcriptional activation by EsaR and ExpR2 was sensitive to the length, position, and identity of these linker insertions. EsaR was more tolerant of a 5-residue alanine insertion at residue 176 than at residue 171, while the opposite was true of ExpR2 (see **Figure 4F**, **G** and **Tables S3** and **S4** for pairwise comparisons). Both EsaR and ExpR2 were more tolerant of 5-glycine vs. 5-alanine insertions at residue 171, suggesting that the flexibility of the linker insertion plays a role in receptor activity. While EsaR and ExpR2 maintained moderate activity with various 5-residue insertions, insertion of 5 alanine or glycine residues rendered MrtR inactive. In MrtR, four-residue insertions of alanine or glycine were tolerated at residue 171 but not at residue 175, and we observed no difference between the addition of alanine vs. glycine residues (**Figure 4B**, **Table S5**).

We were curious if some of the sensitivity to linker insertions observed in EsaR, ExpR2, and MrtR was due to disruption of specific interactions with linker residues. We focused first on EsaR to explore how changes in linker sequence affected activity, building on the result that EsaR showed decreased activity with a scrambled linker. An alanine scan of the EsaR linker (**Figure 4I**) showed that most residues had a mild to moderate effect on activation (0–50% decrease), while others appear to play a crucial role in EsaR function.

Most notably, the EsaR K178A mutant lost the ability to activate or repress transcription, while I180A showed weak activation and moderate repression (**Figure 4I**, **J**). The AlphaFold 3 model of EsaR places I180 in a hydrophobic interaction with the DBD (**Figure 4K**), which may explain its impact on EsaR activity. K178 is solvent-exposed in the model but is predicted with low confidence, preventing strong inference of its molecular role. The importance of K178 to EsaR function may help explain why EsaR Δ176-179 was the only < 8-residue deletion that decreased EsaR activation and why a previous report found that EsaR Δ171-178 was inactive (28), whereas we show above that other EsaR 8-residue linker deletions that maintain K178 are active. ExpR2 K177A, analogous to EsaR K178A, showed a ∼68% decrease in activation, whereas mutation of the adjacent residue, E178A, caused no decrease in ExpR2 activation (**Figure 4G**). The importance of specific linker residues to EsaR and ExpR2 activity helps explain why these receptors did not tolerate long linker extensions.

### Far N-terminal domain perturbations influence activity

A multiple sequence alignment (**Figure S1**) illustrates that dissociative LuxR-type receptors typically have shorter N-termini than associative LuxR-type receptors, but the functional importance of this region is unclear. The AlphaFold 3 structures of EsaR and ExpR2 predict that these receptors lack the far N-terminal helix present in LasR and that hydrophobic residues at the far N-terminus (e.g., F4 and F5 in EsaR, and F4 in ExpR2) are buried between the central β-sheet and the last helix of the LBD (**Figure 5A**, **B**). In support of these computational predictions, similar interactions are observed in crystal structure of the LBD of the dissociative receptor YenR (residues I3 and F6) (10). To probe these potential interactions, we generated EsaR F4A, EsaR F5A and ExpR2 F4A mutants, all of which displayed little or no activity in *E. coli* reporter assays (**Figure 5C-E**). These results support that the distinctive N-terminus observed in these dissociative receptors is crucial for activity.

**Figure 5.**
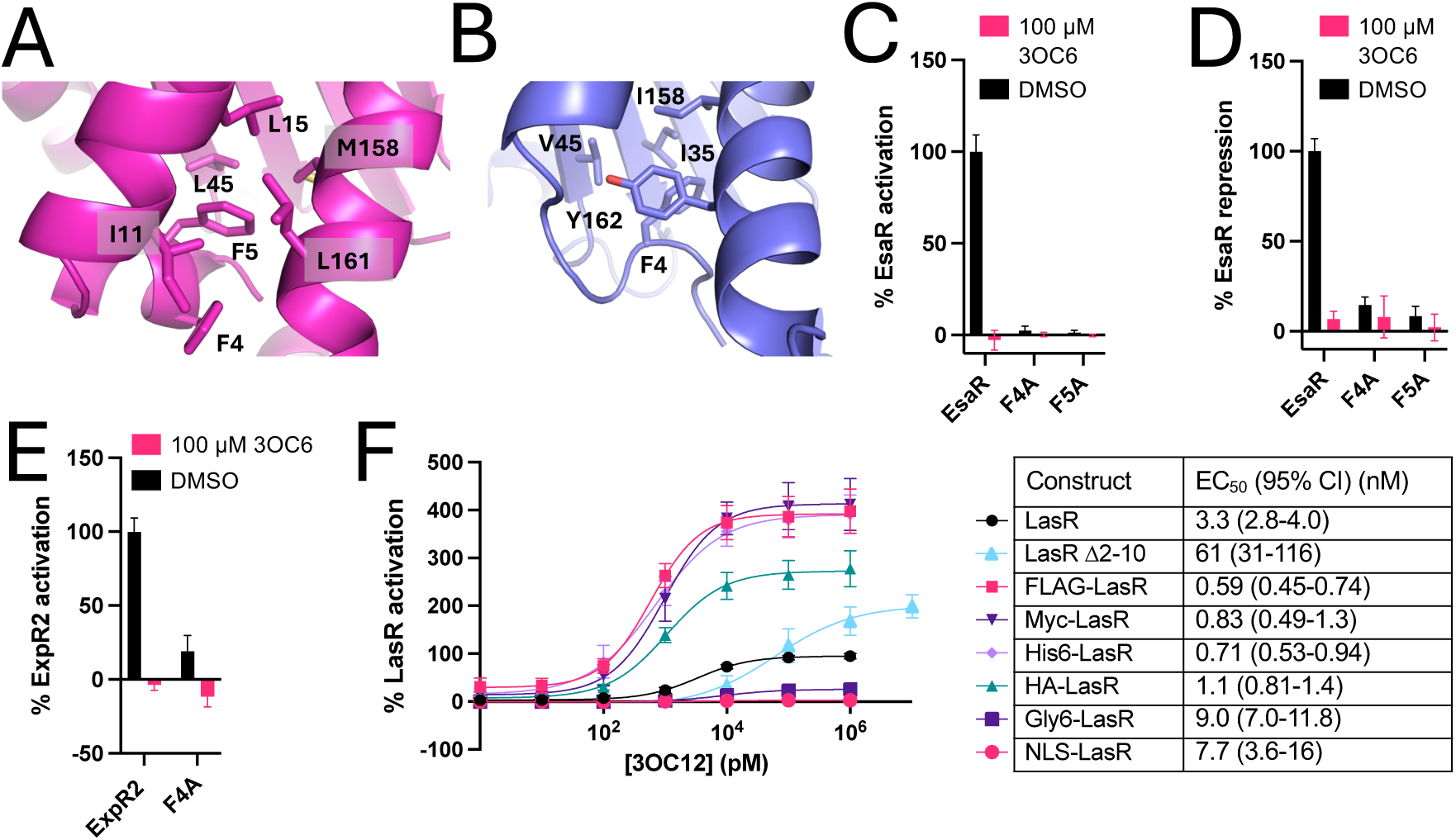
Effect of far N-terminal perturbations on EsaR, ExpR2, and LasR activity in *E. coli* reporter assays. AlphaFold 3-predicted structures of EsaR. **(A)** and ExpR2 **(B)** showing the N-terminus buried between the last helix of the LBD (front right) and the central β-sheet of the LBD (back). The side chains shown are within 4.5 Å of EsaR F4 or F5 or ExpR2 F4. **(C)** Transcriptional activation of the *esaS* promoter by EsaR mutants in *E. coli* BW27749, normalized to uninduced culture (0%) and maximum EsaR activation (100%). **(D)** Transcriptional repression of the *esaR* promoter by EsaR mutants in BW27749, normalized to uninduced culture (0%) and maximum EsaR repression (100%). **(E)** Transcriptional activation of the *rsmA* promoter by ExpR2 mutants in *E. coli* JLD271, normalized to uninduced culture (0%) and maximum ExpR2 activation (100%). **(F)** Transcriptional activation of the *lasI* promoter by LasR mutants, normalized to uninduced culture (0%) and LasR saturated with 3OC12 (100%). **(All graphs)** Data represent at least 3 biological replicates, each performed in triplicate. Error bars represent standard deviation.

To investigate the importance of the extended N-terminus in LasR, we deleted residues 2-10. Intriguingly, LasR Δ2-10 showed *higher* maximum transcriptional activation than LasR. High protein production can push the DNA-binding equilibrium towards the bound state and produce higher maximum activation and signal sensitivity in a cell-based reporter (45, 46), as we observed for LasR with various tags (**Figure 5F**). Surprisingly however, the LasR Δ2-10 mutant showed higher max activation and a ∼16-fold *higher* EC_50_ value than WT LasR (61 vs. 3.3 nM, respectively) in the reporter assay, suggesting that the observed change in ligand sensitivity was not due to differences in protein accumulation. These results highlight the different roles of the far N-terminus in associate vs. dissociative receptors and, to our knowledge, the unrecognized importance of this region for LuxR-type receptor activity and ligand response.

### LasR antagonists are active in chimeras and linker mutants

As a final set of experiments, we were interested in determining how the chimeras and linker mutants introduced above responded to non-native signals, especially competitive antagonists. Our lab (14, 20, 47, 48) and others (13, 14, 16–19) have developed a wide variety of small molecule antagonists of LasR, so we focused first on this receptor. We tested two of the most potent synthetic LasR antagonists developed by our lab, **B7** (49) and **K3** (47) (each a native AHL analog, see **Figure S9** for structures), in two LasR linker extensions (+21 and +42) and two LasR-ExpR2 chimeras (LX-1 and LX-2). We have previously shown that **B7** and **K3** may agonize but do not antagonize ExpR2 in an *E. coli* reporter (25, 26). Compounds were tested for antagonism (in competition against 3OC12) and agonism in reporter strains, and the resulting data are shown in **Figure S9** and summarized in **Table 2**.

**Table 2.**
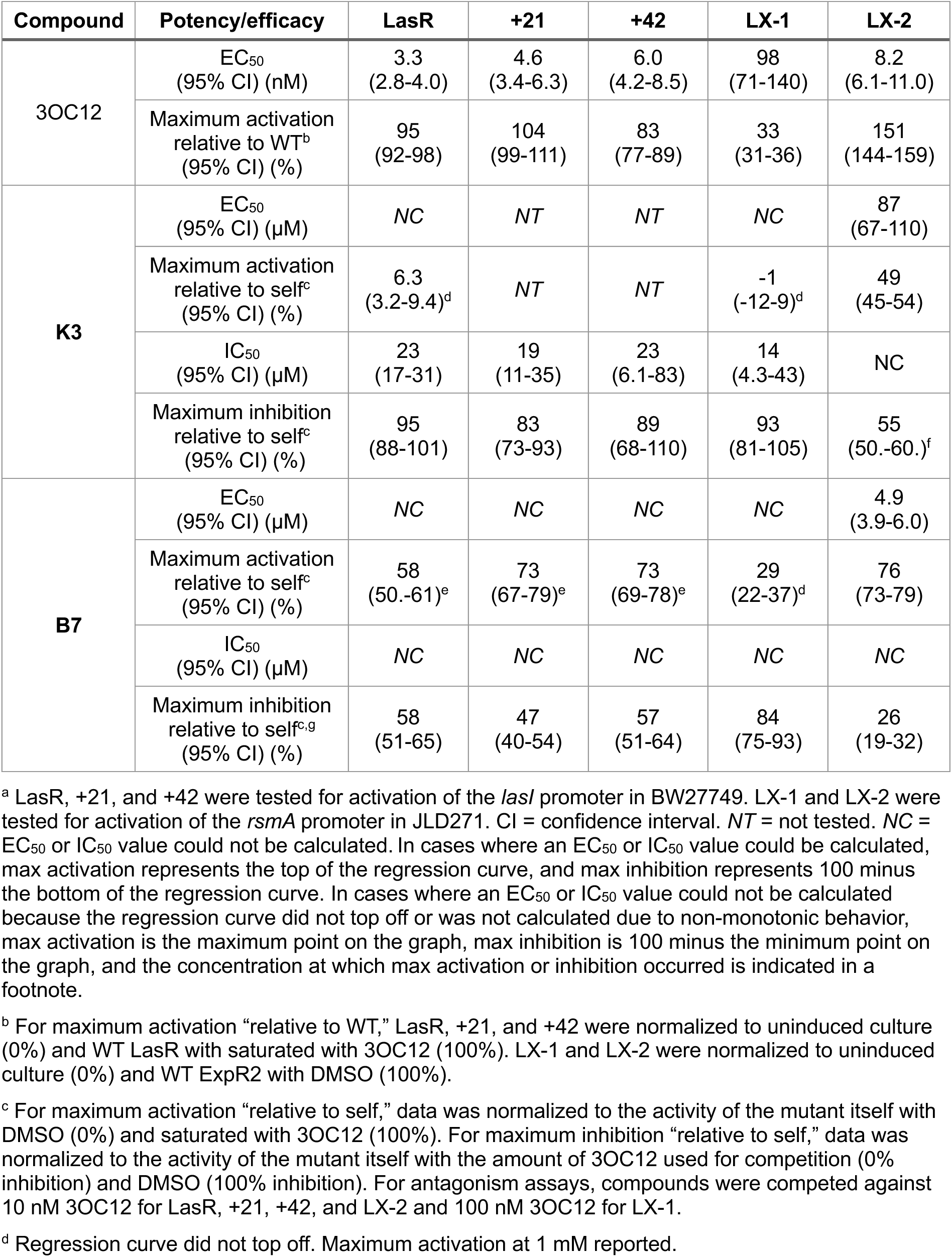
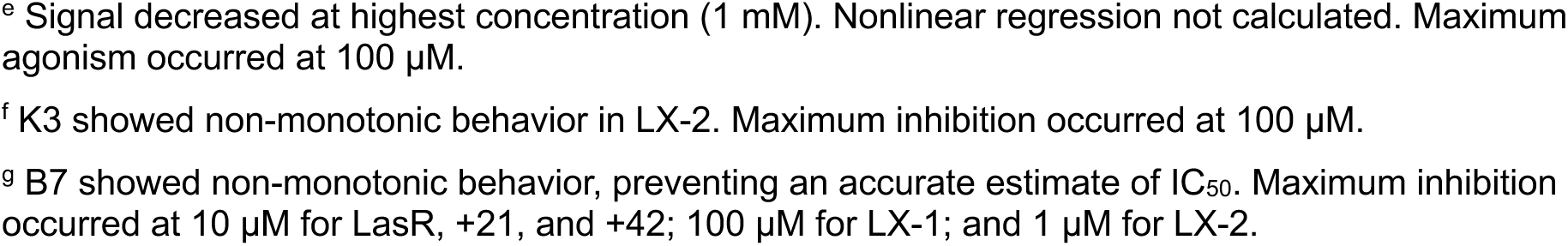
Agonism and antagonism data for selected LasR linker mutants and chimeras with a LasR LBD. Data is based on dose-response curves shown in **Figure S9**.^a^

| Compound | Potency/efficacy | LasR | +21 | +42 | LX-1 | LX-2 |
| --- | --- | --- | --- | --- | --- | --- |
| 3OC12 | EC<sub>50</sub><br>(95% CI) (nM) | 3.3<br>(2.8-4.0) | 4.6<br>(3.4-6.3) | 6.0<br>(4.2-8.5) | 98<br>(71-140) | 8.2<br>(6.1-11.0) |
|  | Maximum activation<br>relative to WT<sup>b</sup><br>(95% CI) (%) | 95<br>(92-98) | 104<br>(99-111) | 83<br>(77-89) | 33<br>(31-36) | 151<br>(144-159) |
| K3 | EC<sub>50</sub><br>(95% CI) (μM) | NC | NT | NT | NC | 87<br>(67-110) |
|  | Maximum activation<br>relative to self<sup>c</sup><br>(95% CI) (%) | 6.3<br>(3.2-9.4)<sup>d</sup> | NT | NT | -1<br>(-12-9)<sup>d</sup> | 49<br>(45-54) |
|  | IC<sub>50</sub><br>(95% CI) (μM) | 23<br>(17-31) | 19<br>(11-35) | 23<br>(6.1-83) | 14<br>(4.3-43) | NC |
|  | Maximum inhibition<br>relative to self<sup>c</sup><br>(95% CI) (%) | 95<br>(88-101) | 83<br>(73-93) | 89<br>(68-110) | 93<br>(81-105) | 55<br>(50-60.)<sup>f</sup> |
| B7 | EC<sub>50</sub><br>(95% CI) (μM) | NC | NC | NC | NC | 4.9<br>(3.9-6.0) |
|  | Maximum activation<br>relative to self<sup>c</sup><br>(95% CI) (%) | 58<br>(50-61)<sup>e</sup> | 73<br>(67-79)<sup>e</sup> | 73<br>(69-78)<sup>e</sup> | 29<br>(22-37)<sup>d</sup> | 76<br>(73-79) |
|  | IC<sub>50</sub><br>(95% CI) (μM) | NC | NC | NC | NC | NC |
|  | Maximum inhibition<br>relative to self<sup>c,g</sup><br>(95% CI) (%) | 58<br>(51-65) | 47<br>(40-54) | 57<br>(51-64) | 84<br>(75-93) | 26<br>(19-32) |

<sup>a</sup> LasR, +21, and +42 were tested for activation of the *lasI* promoter in BW27749. LX-1 and LX-2 were tested for activation of the *rsmA* promoter in JLD271. CI = confidence interval. NT = not tested. NC = EC<sub>50</sub> or IC<sub>50</sub> value could not be calculated. In cases where an EC<sub>50</sub> or IC<sub>50</sub> value could be calculated, max activation represents the top of the regression curve, and max inhibition represents 100 minus the bottom of the regression curve. In cases where an EC<sub>50</sub> or IC<sub>50</sub> value could not be calculated because the regression curve did not top off or was not calculated due to non-monotonic behavior, max activation is the maximum point on the graph, max inhibition is 100 minus the minimum point on the graph, and the concentration at which max activation or inhibition occurred is indicated in a footnote.
<sup>b</sup> For maximum activation “relative to WT,” LasR, +21, and +42 were normalized to uninduced culture (0%) and WT LasR with saturated with 3OC12 (100%). LX-1 and LX-2 were normalized to uninduced culture (0%) and WT ExpR2 with DMSO (100%).
<sup>c</sup> For maximum activation “relative to self,” data was normalized to the activity of the mutant itself with DMSO (0%) and saturated with 3OC12 (100%). For maximum inhibition “relative to self,” data was normalized to the activity of the mutant itself with the amount of 3OC12 used for competition (0% inhibition) and DMSO (100% inhibition). For antagonism assays, compounds were competed against 10 nM 3OC12 for LasR, +21, +42, and LX-2 and 100 nM 3OC12 for LX-1.
<sup>d</sup> Regression curve did not top off. Maximum activation at 1 mM reported.
<sup>e</sup> Signal decreased at highest concentration (1 mM). Nonlinear regression not calculated. Maximum agonism occurred at 100 μM.
<sup>f</sup> K3 showed non-monotonic behavior in LX-2. Maximum inhibition occurred at 100 μM.
<sup>g</sup> B7 showed non-monotonic behavior, preventing an accurate estimate of IC<sub>50</sub>. Maximum inhibition occurred at 10 μM for LasR, +21, and +42; 100 μM for LX-1; and 1 μM for LX-2.

LasR linker extensions responded like WT LasR in both agonism and antagonism assays. In a competition assay against 3OC12, **B7** and **K3** both produced almost identical IC_50_ and maximum inhibition values in the LasR linker extensions as in WT LasR. Like many LasR antagonists, **B7** antagonizes LasR at intermediate concentrations but acts as an agonist at high concentration (50), and this non-monotonic activity was observed in the linker extensions as well. These results are congruent with the native agonist 3OC12 having similar activities and potency in the linker mutants vs. WT LasR and suggest that these changes in the linker do not alter how LasR responds to either agonist or antagonists.

LasR-ExpR2 chimeras (LX-1 and LX-2) were also antagonized by **B7** and **K3** (**Table 2**, **Figure S9**), though their overall activity and sensitivity to both agonists and antagonists differed from WT LasR. Chimera LX-2 showed ∼4.5-fold higher activity than LX-1 and was more strongly agonized and less strongly antagonized than LasR and LX-1. High protein expression, which can push the DNA-binding equilibrium towards the bound state, could explain the high activity and weak antagonism of LX-2 (45, 46). The fact that **B7** and **K3** antagonize LX-1 and LX-2 is consistent with a model based on ongoing work in our lab, where certain LasR antagonists exert their influence mainly by modulating the stability of the LasR LBD.

In addition to LasR antagonists, we tested whether an MrtR antagonist could inhibit an MrtR-ExpR2 chimera. MrtR and MX-1 were both agonized by 7*Z*-3OHC14 and fully antagonized by 3OC8 (**Table 3**, **Figure S9**), suggesting that the activity of this antagonist depends primarily on the LBD, analogous to the LasR-ExpR2 chimeras tested above.

**Table 3.** Agonism and antagonism data for MrtR and an MrtR-ExpR2 chimera. Data based on dose-response curves shown in **Figure S9**.^a^

| Compound | Potency/efficacy | MrtR | MX-1 |
| --- | --- | --- | --- |
| 7Z-3OHC14 | EC<sub>50</sub> (95% CI) (nM) | 1.7 (1.3-2.1) | 9.6 (7.9-12) |
|  | Maximum activation relative to WT<sup>b</sup> (95% CI) (%) | 96 (93-99) | 144 (136-152) |
| 3OC8 | EC<sub>50</sub> (95% CI) (nM) | *NT* | *NT* |
|  | Maximum activation relative to self<sup>c</sup> (95% CI) (%) | -3 (1 to -7) | -2 (-1 to -3) |
|  | IC<sub>50</sub> (95% CI) (nM) | 670 (500-900) | 660 (440-970) |
|  | Maximum inhibition relative to self<sup>c</sup> (95% CI) (%) | 97 (92-102) | 108 (100-116) |

<sup>a</sup> MrtR was tested for activation of the *mrtI* promoter in BW27749. MX-1 was tested for activation of the *rsmA* promoter in JLD271. CI = confidence interval. *NT* = not tested. In cases where an EC<sub>50</sub> or IC<sub>50</sub> value was calculated, maximum activation represents the top of the regression curve, and maximum inhibition represents 100 minus the bottom of the regression curve. Maximum activation by 3OC8 represents activity at 100 μM 3OC8.
<sup>b</sup> For maximum activation “relative to WT,” MrtR was normalized to uninduced culture (0%) and WT MrtR saturated with 7Z-3OHC14 (100%). MX-1 was normalized to uninduced culture (0%) and ExpR2 with DMSO (100%).
<sup>c</sup> For maximum activation “relative to self,” data was normalized to uninduced culture (0%) and to the activity of the mutant itself with 1 μM 7Z-3OHC14 (100%). For maximum inhibition “relative to self,” data was normalized to the activity of the mutant itself with the amount of 7Z-3OHC14 used for competition (0% inhibition) and uninduced culture (100% inhibition). For antagonism assays, 3OC8 was competed against 8 nM 7Z-3OHC14 for MrtR and 40 nM 7Z-3OHC14 for MX-1.

## DISCUSSION

In the current study, we sought to investigate features that contribute to associative vs. dissociative mechanisms in LuxR homologs, as a step toward delineating the molecular mechanisms of ligand recognition and signal transduction in this family of bacterial transcription factors. We have shown that, among the four LuxR-type receptors examined (LasR, MrtR, EsaR, and ExpR2), the LBD determines whether a receptor displays associative vs. dissociative activity. Similarly, previous work with the LacI/GalR family of transcriptional repressors found that a chimera with the DBD of the dissociative LacI and the LBD of the associative PurR showed associative activity (51), and although several dissociative LacI homologs have been evolved to display associative activity, specifically altering the DNA recognition sequence did not reverse receptor mechanism (52).

Surprisingly, we found that both EsaR-LasR chimeras and WT EsaR lacking residues 159-162 showed associative activity. The relatively minor change of deleting these four residues, or even just Y159, essentially converted EsaR from a dissociative to an associative repressor. Such a mechanistic switch has not been previously reported for a LuxR-type receptor, but minor sequence changes have been shown to invert the ligand response of other ligand-responsive transcription factors, including TetR, LacI, LsrR, BetI, and BenM (52–57).

We also performed an extensive study of the linker domain in these four receptors, to learn how this understudied portion of LuxR homologs that unites the LBD and DBD affects receptor activity. The results of our analyses suggest key structural and mechanistic differences between associative and dissociative receptors and within each class. The receptors examined here tolerate a similar minimum linker length, suggesting that an extended linker is not required for dissociative receptor function, as has been previously suggested (28). Several related dissociative receptors, including EsaR and ExpR2, fall into what has been termed the “EsaR clade” (58), and the extended linker observed in this group may simply be a vestige of receptor evolution. The associative receptor CarR of *P. carotovorum* and the ligand-unresponsive CarR of *Serratia* sp. 39006 share 33-40% sequence identity with members of the EsaR clade and contain the extended linker often observed in dissociative receptors. It is possible that these CarR receptors represent intermediates in the divergent evolution of associative and dissociative LuxR-type receptors (37). In addition, it has been suggested that the dissociative mechanism may have evolved multiple times independently (59). For example, the dissociative receptors VjbR of *Brucella melitensis* and CepR2 of *Burkholderia cenocepacia* share low sequence identity with members of the EsaR clade and with each other (59). VjbR and CepR2 may utilize mechanisms of ligand response vastly different from EsaR and ExpR2, and it would be interesting to investigate their behavior in chimeras and their response to structural mutations.

Responses to a lengthened linker differed dramatically by receptor, suggesting that the linker plays diverse roles in receptor activation. LasR maintained activity with exceptionally long linker additions (>100 extra residues), which supports the hypothesis that the LasR linker functions mainly as a flexible tether between domains, rather than meditating specific interdomain interactions. EsaR, ExpR2, and MrtR, in contrast, lost activity even with 5-residue insertions, suggesting that these receptors may rely on specific interactions between the LBD, linker, and/or DBD that may be influenced by linker length and composition. The relative positions of the LBD and DBD have been shown to affect the activity of some LuxR-type receptors. As perhaps the most prominent example so far, the crystal structure of CviR of *Chromobacterium subtsugae* bound to the synthetic AHL-type antagonist CL reveals the DBDs interacting with the LBD of the opposite monomer, resulting in a splayed DBD conformation that is unfavorable for DNA binding (24). In addition, several synthetic antagonists of QscR of *P. aeruginosa* lead to an inactive dimer conformation (11). As in these examples, the inactivating linker mutations discovered in this study may place the LBD and DBD at unfavorable relative positions, possibly preventing specific interactions between the LBD, DBD, and/or linker. A better understanding of these interactions may shed light on how ligand binding affects receptor activity. For example, EsaR remains stable and dimerized in the presence and absence of ligand (23), suggesting that ligand-induced inactivation must occur via a mechanism other than a decrease in protein accumulation or dimerization. Our work here revealed that EsaR is highly sensitive to linker perturbations, and it is possible that changes in interactions between the LBD and DBD, rather than between monomers, are the mechanism by which ligand binding alters EsaR activity. In addition, although LasR and MrtR both function via an associative mechanism, whereby ligand binding induces stabilization and/or dimerization, they respond very differently to linker extensions. Ongoing structural and biochemical work aims to begin to define at a molecular level the interdomain interactions that govern LuxR-type receptor activity and ligand response and will be reported in due course.

So, we close how we started — by noting that many putative LuxR homologs have been identified but few characterized. This study provides several new and significant insights into their mechanisms of activation and ligand response. That said, the functional differences observed in the four receptors investigated herein likely represent a small fraction of the mechanistic diversity of LuxR-type receptors. Additional mechanistic and structural investigations are required to define the various molecular mechanisms of ligand response within this family and will create a more complete picture of the functional diversity among LuxR-type receptors.

## MATERIALS AND METHODS

See supplemental methods for standard reagent and instrumentation details.

**Compound handling.** Native AHLs used for screening were purchased from Sigma-Aldrich (3OC12) and Cayman Chemical (3OC6 and 7*Z*-3OHC14). Non-native AHL inhibitors (**B7** and **K3**) were acquired from in-house stocks synthesized according to published methods (47, 49). AHLs were dissolved in anhydrous DMSO to 10 mM and stored at –20 °C in Teflon-capped glass vials.

**Plasmid construction and reporter assays.** Full sequences of plasmids used for EsaR, ExpR2, and LasR reporters have been reported previously (25, 26, 60). Plasmid descriptions, gBlocks, and primers are provided in **Tables S2, S6, and S7**. Mutants were created using the Q5 Site-Directed Mutagenesis Kit (New England Biolabs cat. no. E0554S) or Gibson assembly (New England Biolabs cat. no. E2611). Reporters were constructed in *E. coli* JLD271 (61) or *E. coli* BW27749 (62); the latter enables control of protein production levels based on arabinose concentration. Reporter assays were performed as previously described (27), with minor optimization for each WT reporter strain. See supplemental methods for details.

**Protein structure predictions.** The AlphaFold 3 (63) web server was used to predict protein structures for EsaR, ExpR2, and MrtR. Results were visualized in PyMol (version 3.1.1).

## ACKNOWLEDGEMENTS

Financial support for this work was provided by the NIH (R35 GM131817). I.M.S. was supported in part by an NSF Graduate Research Fellowship (DGE-2137424) and is an affiliate of the UW−Madison NIH Chemistry−Biology Interface Training (CBIT) Program (T32 GM008505 and T32 GM152341). S.A.W. acknowledges support by the UW–Madison College of Letters & Science Honors Program and a Department of Chemistry Eugene and Patricia Kreger Herscher Scholarship. D. Widner and E. Santa are acknowledged for sharing technical expertise. We thank the laboratory of J. Martell for the use of their Azure gel imager. Figure 1A created with BioRender.com.

