## Supplemental methods and data for "Domain-swapped LuxR-type quorum sensing receptors reveal divergent ligand-response mechanisms among homologs"

**CONTENTS**


- Supplemental methods
- Discussion of accumulation of chimeric proteins in reporter strains
- Discussion of effect of FLAG tag on reporter activity
- Discussion of activity of DBDs alone
- **Figure S1.** Sequence alignment of selected LuxR-type receptors
- **Figure S2.** Sequence alignment comparison of CinR vs. MrtR and CinI vs. MrtI
- **Figure S3.** Response of chimeras to ligands corresponding to the LBD vs. DBD receptor
- **Figure S4.** Additional chimera and mutant activity data
- **Figure S5.** Effects of FLAG tag on activity
- **Figure S6.** Activity of DBDs alone
- **Figure S7.** Ligand potency in EsaR and ExpR2 linker deletion mutants
- **Figure S8.** Activity and accumulation of FLAG-tagged EsaR linker addition mutants
- **Figure S9.** Small molecule inhibition of select mutants and chimeras
- **Figure S10.** Optimization of arabinose concentration for *E. coli* BW27749 reporters
- **Figure S11.** Additional biological replicate of western blots
- **Table S1.** Protein sequences of WT LuxR-type receptors examined in this study
- **Table S2.** Bacterial strains and plasmids used in this study
- **Table S3.** Ordinary one-way ANOVA with Šídák's correction for multiple comparisons for activity of EsaR +5 linker mutants with DMSO, corresponding to data in **Figure 4F**
- **Table S4.** Ordinary one-way ANOVA with Šídák's correction for multiple comparisons for activity of ExpR2 +5 linker mutants with DMSO, corresponding to data in **Figure 4G**
- **Table S5.** Ordinary one-way ANOVA with Šídák's correction for multiple comparisons for activity of MrtR +4 linker mutants with 7Z-3OHC14, corresponding to data in **Figure 4B**
- **Table S6.** gBlocks used in this study
- **Table S7.** Primers used in this study
- Raw western blot images
- References

### Supplemental methods.

**General reagent and instrumentation information.** Arabinose was purchased from MilliporeSigma; isopropyl β-D-1-thiogalactopyranoside (IPTG), ampicillin, and gentamycin from Dot Scientific; and chloramphenicol from Sigma-Aldrich. Water (18 MΩ) was purified with a Sartorius Arium Pro system. Antibiotics stocks were prepared at 1000x working concentrations in water (gentamycin and ampicillin) or ethanol (chloramphenicol) and stored at -20 °C. Luminescence and fluorescence data were collected on a BioTek Synergy 2 plate reader using Gen5 software (version 3.12). AHLs were dissolved in anhydrous DMSO to 10 mM and stored at -20 °C in Teflon-capped glass vials. Arabinose stocks were prepared at 100x working concentration in water and stored at room temperature of up to 6 months.

**Bacteriology.** The bacteria and plasmids used in this study are listed in **Table S2**. Strains were stored in 25% glycerol at -80 °C. Cultures were grown in Lennox Broth (LB) (Research Products International) with appropriate antibiotics and incubated at 37 °C with shaking (200 rpm). Chemically competent cells were prepared in-house and used for cloning.

**Plasmid and reporter construction.** Plasmid descriptions, gBlocks, and primers are provided in **Tables S2**, **S6**, **and S7**. The sequences of all plasmids created for this study were confirmed by sequencing. To create pJN105-mrtR, the MrtR coding sequence was codon-optimized and purchased as a gBlock from Integrated DNA Technologies (IDT), then inserted by Gibson assembly into pJN105. To create pmrtI:GFP, a fragment of the MrtR-regulated *mrtI* promoter (1) consisting of bases -120 to -1 relative to the *mrtI* translational start site was ordered as a gBlock from IDT and inserted by Gibson assembly into pesaR-AC:GFP, replacing the *esaS* promoter.

Chimeras were constructed using Gibson assemblies with a backbone that matched the backbone of the WT protein production plasmid to which the chimera was compared in reporter assays. Mutants of individual WT receptors and FLAG-tagged constructs were created using the Q5 Site-Directed Mutagenesis Kit according to the manufacturer's protocol (New England Biolabs cat. no. E0554S) or Gibson assembly (New England Biolabs cat. no. E2611).

All chimera reporters were created by transforming the appropriate protein production plasmid and reporter plasmid (e.g., pJN105-lasR and pSC11-lasI for a WT LasR reporter) into *E. coli* JLD271. Reporters for mutants of the individual receptors LasR, EsaR, and MrtR were constructed in *E. coli* BW27749, which enables control of protein production levels based on arabinose concentration (2) and can allow for detection of more subtle differences between mutants. Because ExpR2 had a relatively low dynamic range in JLD271 with saturating arabinose, decreasing protein production was not practical, and ExpR2 mutants were tested in JLD271.

**Transcriptional reporter assays.** Overnight cultures of reporter strains were diluted 1:10 in fresh LB medium with 100 µg/mL ampicillin and 15 µg/mL gentamicin. Cultures were grown to an $OD_{600}$ of 0.22-0.27 as measured in a 96-well microtiter plate with 200 µL culture per well. Assays were performed in black 96-well microtiter plates with transparent bottoms (Corning 3904) for GFP reporters (EsaR, ExpR2, and MrtR) and clear plates (Corning 3370) for β-galactosidase reporters (LasR). Aliquots (2 µL) of compound stocks or DMSO (vehicle control) were added to each well. Cultures were induced by adding 1:100 of a 100x stock of arabinose, and 198 µL of culture was added to each well. Uninduced culture served as a control. Plates were incubated at 37 °C with shaking at 200 rpm for 4 h for β-galactosidase reporters and 3.5 h for GFP reporters. Plates were mixed by manual pipetting before reading GFP fluorescence and $OD_{600}$ for GFP reporters or $OD_{600}$ for β-galactosidase reporters using a plate reader. To detect β-galactosidase, cultures were diluted 1:10 in water, and 10-μL aliquots of diluted cells were mixed with 10 μL of 1:2 diluted Beta-Glo substrate (Promega, cat. No. E4720) in a white 384-well microtiter plate (Corning 3574). The Beta-Glo reaction was incubated for 30 min at 28 °C before reading luminescence signal on a plate reader. Microsoft Excel (version 16.92) was used to normalize data, and GraphPad Prism (version 10.2.0) was used to generate graphs and perform statistical analyses. Where outliers were suspected, a ROUT test with Q = 5% was used to test for outliers.

All JLD271 reporters were induced with a saturating concentration of arabinose (0.4 mg/mL for EsaR, ExpR2, and MrtR and 4 mg/mL for LasR reporters). Arabinose concentration was optimized for each WT receptor BW27749 reporter (**Figure S10**), and the same concentration was used for all the mutants in each reporter system. Unless noted in the figure legend, arabinose concentrations used in BW27749 were 1 μg/mL for the LasR reporter (PSC11-lasI), 0.5 μg/mL for the EsaR-activated reporter (pesaR-AC:GFP), 2 μg/mL for the EsaR-repressed reporter (pesaR:GFP), and 0.25 μg/mL for the MrtR reporter (pmrtI:GFP). Because FLAG-tagged EsaR showed decreased signal relative to WT EsaR, we used 0.4 mg/mL arabinose for reporters and western blots of FLAG-tagged EsaR and point mutants. In cases where mutant data were normalized to WT LasR or MrtR saturated with ligand, ≥1 μM 3OC12 or 7Z-3OHC14 was used (see **Figure S9** for dose-response curves). In cases where data were normalized to maximum EsaR activation, maximum EsaR repression, or maximum ExpR2 activation, maximum activity corresponds to the vehicle control (DMSO) condition.

**Western blots.** Cultures were grown as described for bacterial reporters, either in 96-well microtiter plates or in test tubes. The $OD_{600}$ of each sample was measured using a plate reader, and X mL of culture was pelleted, where X = 0.25/$OD_{600}$. Cell pellets were resuspended in 100 μL MilliQ water and stored at -20 °C until SDS-PAGE analysis (see SDS-PAGE protocol below). Gels were equilibrated in transfer buffer for 20 min and trimmed to a 15-70 kDa range before transfer. PVDF membrane (Immobilon-P) was soaked in methanol for 30 s to 1 min, then membrane, filter paper, and sponges were equilibrated in chilled transfer buffer (10 mM CAPS, 10% methanol, pH 10-11) for 20 min. Transfer was performed using a Bio-Rad Mini-PROTEAN Tetra Cell, with ice packs and on a stir plate, for 120 min at 40V. Blots were stained immediately or stored at 4 °C overnight between sheets of wet filter paper. Blots were stained with HRP-conjugated DYKDDDDK Tag Monoclonal antibody from Proteintech (Cat No. HRP-66008) to visualize FLAG-tagged proteins, or with HRP-conjugated GAPDH Monoclonal antibody from Proteintech (Cat No. HRP-66004) to visualize GAPDH as a loading control. In both cases, blots were blocked (EveryBlot Blocking Buffer, Bio-Rad) for 20 min, and then antibodies were added at a 1:15,000 dilution for 50 min to 1 hr. Blots were washed with TBST (20 mM Tris base, 150 mM NaCl, 0.1% Tween 20) once for 10 sec, then three times for 5-10 min. Blots were visualized with Radiance Plus (Azure Biosystems) on an Azure Biosystems C400 imager. After imaging, blots were incubated with 10% acetic acid for 30 min at 37 °C to inactivate HRP, then re-probed for the loading control. Alternatively, duplicate blots were performed for FLAG and GAPDH staining. Brightness and contrast of images was adjusted to improve visibility of bands, and no non-linear corrections were applied. Raw images are shown at the end of this SI document.

**SDS-PAGE for western blot and solubility assays.** Samples were prepared with 12 μL sample, 4.5 μL of 4x LDS buffer (G-Biosciences cat. #786323), and 1.5 μL of 20 mM DTT, then heated to 95 °C for 10 min before loading on a 10% Mini-PROTEAN TGX gel (Bio-Rad). Gels were run on ice for 100 min at 120V. Gels were used for western blots, as described above, or imaged using a Cytiva Amersham Typhoon imager for solubility assays.

**Solubility assay.** Proteins were expressed from *E. coli* BL21(DE3) harboring pLysS and a pET17b plasmid expressing the protein of interest. pET17b-FLAG-lasR was constructed by inserting a FLAG tag between LasR residues 1 and 2 in pET17b-lasR. Overnight cultures were diluted 1:100 in LB medium with 100 µg/mL ampicillin and 34 µg/mL chloramphenicol and grown at 37 °C with 200 rpm shaking to an $OD_{600}$ of 0.5-0.7, as measured using a cuvette on a Nanodrop 2000c spectrophotometer (Thermo Scientific). Cultures were chilled on ice for 10 min and supplemented with 0.5 µM IPTG to induce protein expression. Compound was added as a 1:1000 dilution of a 10 mM stock of 3OC12 in DMSO, and an equivalent volume of DMSO was added to control cultures. Cultures were incubated at 17 °C with shaking at 200 rpm for 16 hr. Cells were pelleted by centrifugation, resuspended in TEDG buffer (50 mM Tris HCl, 500 mM NaCl, 1 mM EDTA, 10% glycerol, 1 mM DTT, pH 7.4, supplemented with 10 µM 3OC12 or DMSO and a protease inhibitor cocktail) and lysed by sonication. Samples were taken of the whole cell lysate, then the lysate was centrifuged at 40,000g for 30 min. The supernatant was collected as the soluble fraction. Whole cell and soluble fractions were stored at -20 °C until analysis by SDS-PAGE.

**Sequence alignment generation.** Sequences from the National Center for Biotechnology Information (NCBI) database were aligned using Clustal Omega (3) and visualized with MView (3).

### Discussion of accumulation of chimeric proteins in reporter strains.

To investigate whether differences in chimera activity could be explained by differences in protein accumulation in the reporter strain, reporter strains expressing FLAG-tagged chimeras were analyzed via western blotting (see supplemental methods). Several EsaR-LasR chimeras with varying levels of activity were selected to test whether differences in activity correlated with differences in expression. All four LasR-EsaR chimeras were tested to see if their low activity was due to lack of protein accumulation. In some cases, the N-terminal FLAG tag affected activity, possibly due to changes in protein expression (**Figure S5A-D**; see discussion below). LasR is known to have high proteolytic turnover (4) and to be poorly soluble in the absence of a stabilizing ligand (5). EsaR has shown comparable accumulation in cells in the presence and absence of ligand but is more resistant to in vitro proteolysis in the presence of its cognate ligand, 3OC6 (6). Among dissociative EsaR-LasR chimeras, we observed that more active constructs showed higher accumulation than less active constructs and, like WT EsaR, EsaR-LasR chimeras showed comparable accumulation with and without ligand (**Figure S5E**). In contrast, LasR-EsaR chimeras showed robust accumulation only in the presence of ligand, as did WT LasR (**Figure S5E**). These results suggest that, like activity, accumulation in the absence of ligand depends on the characteristics of the LBD receptor. In addition, the low activity of LasR-EsaR chimeras is not due to a lack of accumulation in the reporter strain.

### Discussion of effect of FLAG tag on reporter activity.

To aid in visualization for western blotting, a FLAG tag was added to the N-terminus of each protein. In some cases, the addition of a FLAG or FLAG-Gly5 tag to a receptor affected its activity in a transcriptional reporter. EsaR was inactive with a FLAG tag but partially active when a Gly-5 linker was added after the FLAG tag (**Figure S5A** and **B**), so a FLAG-Gly5 tag was used for chimeras with the EsaR N-terminus. The addition of an N-terminal FLAG tag to WT LasR or a FLAG-Gly5 tag to chimeras with a LasR DBD dramatically increased activity in reporter assays (**Figure S5C** and **D**). Other tags also caused large changes in LasR activity in the reporter assay (**Figure 5F**). We considered that the FLAG tag may have a stabilizing effect on LasR in the absence of ligand, but a solubility assay showed that, like WT LasR, FLAG-LasR is soluble in the presence of 3OC12 but not in its absence (**Figure S5F**).

Alternatively, the presence of an N-terminal tag may alter protein expression by affecting mRNA folding around the translation initiation region (TIR) (7, 8). Higher protein expression would be expected to lead to higher activity in a transcriptional reporter assay. Notably, although WT LasR and chimeras with a LasR DBD differ in LBD, their respective protein production plasmids have the same sequence immediately upstream of the translational start site, and this sequence is slightly different from the plasmid expressing EsaR and chimeras with an EsaR DBD. Chimeras were constructed so that the backbone matched the backbone of the WT plasmid to which they were compared in reporter assays. While chimera EL-10 expressed from the same backbone as LasR showed a 13-fold increase in activity upon the addition of a FLAG tag, EL-10 expressed from that plasmid with 12 bp removed upstream of the start codon showed ~9-fold higher activity than EL-10 expressed from the original plasmid and showed a slight *decrease* in activity upon the addition of a FLAG tag (**Figure S5G**). These results support the hypothesis that tag-induced changes in LasR activity are due to changes in the TIR, rather than changes to protein activity.

In some cases, the FLAG tag also affected ligand responsiveness. While chimeras EL-3 and EL-6 showed decreased activity in the presence of 3OC6, FLAG-Gly5-EL-3 and FLAG-Gly5-EL-6 showed much higher overall signal than their untagged counterparts and no response to ligand. A very high level of protein expression could overwhelm the detection limit of the reporter system, causing receptors to appear constitutively active. To test the effects of protein expression level on ligand responsiveness, we expressed FLAG-Gly5-EL-3 and FLAG-Gly5-EL-6 in an *E. coli* BW27749 reporter strain in which protein expression could be controlled based on the concentration of arabinose used for induction. Decreasing arabinose concentrations led to a dramatic decrease in overall signal but did not restore ligand responsiveness in FLAG-Gly5-EL-3 and FLAG-Gly5-EL-6 (**Figure S5H-I**), suggesting that the lack of ligand response is not an artifact of protein expression levels.

### Discussion of activity of DBDs alone.

We note that the activity of chimeras with LasR, ExpR2, or EsaR DBDs was not due to constitutive activity of the DBDs alone. We found that the ExpR2 DBD (Δ2-175) was unable to activate transcription in the ExpR2 reporter (**Figure S6A**). Likewise, the LasR LBD (Δ4-161 and Δ4-168) was inactive in a LasR reporter, but these constructs recovered ~9 and ~3% activity, respectively, upon addition of an N-terminal FLAG tag, potentially due to an increase in protein expression (**Figure S6B**). The EsaR DBD (Δ2-160 and Δ2-178) has been reported to be unstable alone (9). These results indicate that the LBD, both in the native receptors and chimeras, is crucial for activity.

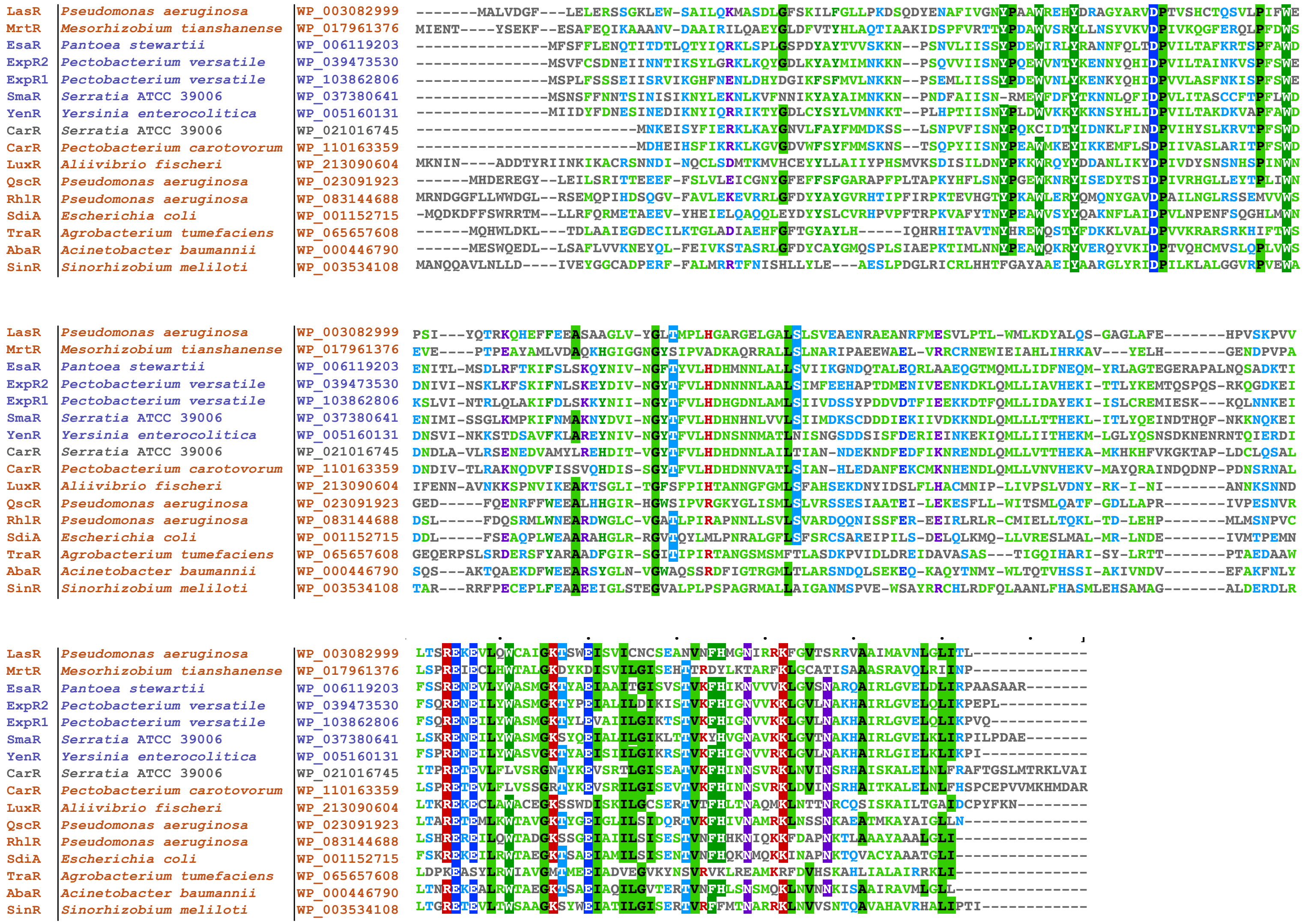


**Figure S1. Sequence alignment of selected LuxR-type receptors.** Sequences shown in purple are dissociative (9-12), those in orange are associative (11, 13-17), and CarR (gray) is unresponsive to ligand (11).

A

```
AAL59595.1_CinR_R_etli      MIENTYSDKFEPAFEQIKAAPNVDAAIRILQAEYGLDFVTYHLAQTIASKIDSPFVRTTY
MrtR                        MIENTYSEKFESAFEQIKAAANVDAAIRILQAEYGLDFVTYHLAQTIAAKIDSPFVRTTY
                            *******:*** ******** ***************************:***********

AAL59595.1_CinR_R_etli      PDAWVSRYLLNSYVKVDPIVKQGFERQLPFDWSEVEPTPEAYAMLVDAQKHGIGGNGYSI
MrtR                        PDAWVSRYLLNSYVKVDPIVKQGFERQLPFDWSEVEPTPEAYAMLVDAQKHGIGGNGYSI
                            ************************************************************

AAL59595.1_CinR_R_etli      PVADKAQRRALLSMNARIPADHWAELVRRCRNEWIEIAHLIHQKAVYELYGENDPVPALS
MrtR                        PVADKAQRRALLSLNARIPAEEWAELVRRCRNEWIEIAHLIHRKAVYELHGENDPVPALS
                            *************:******:.*********************:*******:********

AAL59595.1_CinR_R_etli      PREIECLHWTALGKDYKDISVILGISEHTTRDYLKTARFKLGCATISAAASRAVQLRIINP   241
MrtR                        PREIECLHWTALGKDYKDISVILGISEHTTRDYLKTARFKLGCATISAAASRAVQLRIINP   241
                            *************************************************************
```

B

```
AAF89990.1_CinI_R_leguminosarum      MFVIIQAHEYQKYAAVLDQMFRLRKKVFADTLCWDVPVIGPYERDSYDSLAPAYLVWCND
AAZ32755.1_MrtI_M_tianshanense       MFIIIQAHEYQKYAAVLDQMFRLRKKVFADTLGWDVPVIGPYERDSYDSLAPAYLVWCND
                                     **:***************************** ***************************

AAF89990.1_CinI_R_leguminosarum      SRTRLYGGMRLMPTTGPTLLYDVFRETFPDAADLIAPGIWEGTRMCIDEEAIAKDFPEID
AAZ32755.1_MrtI_M_tianshanense       SRTRLYGGMRLMPTTGPTLLYDVFRETFPDAANLIAPGIWEGTRMCIDEEAIAKDFPGID
                                     ********************************:************************ **

AAF89990.1_CinI_R_leguminosarum      AGRAFSMMLLALCECALDHGIHTMISNYEPYLKRVYKRAGAEVEELGRADGYGKYPVCCG
AAZ32755.1_MrtI_M_tianshanense       AGRAFSMMLLALCECALDHGIHTMISNYEPYLKRVYKRAGAEVEELGRADGYGKYPVCCG
                                     ************************************************************

AAF89990.1_CinI_R_leguminosarum      AFEVSDRVLRKMRAALGLTLPLYVRHVPARSVVTQFLEMAA  221
AAZ32755.1_MrtI_M_tianshanense       AFEVSDRVLRKMRAALGLTLPLYVRHVPARSVVTQFLEMAA  221
                                     *****************************************
```

**Figure S2. Sequence alignment comparison of CinR vs. MrtR and CinI vs. MrtI. (A)** Sequence alignment of CinR of *Rhizobium etli* and MrtR of *Mesorhizobium tianshanense* with matching translational start sites. CinR and MrtR are 96% identical. **(B)** Sequence alignment of AHL synthases CinI of *Rhizobium leguminosaurum* and MrtI of *M. tianshanense*. CinI and MrtI are 98% identical. NCBI accession numbers are indicated at the start of each row, except for MrtR, as the chosen translational start site differs from the MrtR sequence available in the NCBI database (AAZ32754.1) but matches non-redundant protein sequence WP_017961376. Alignments created with Clustal Omega (3).

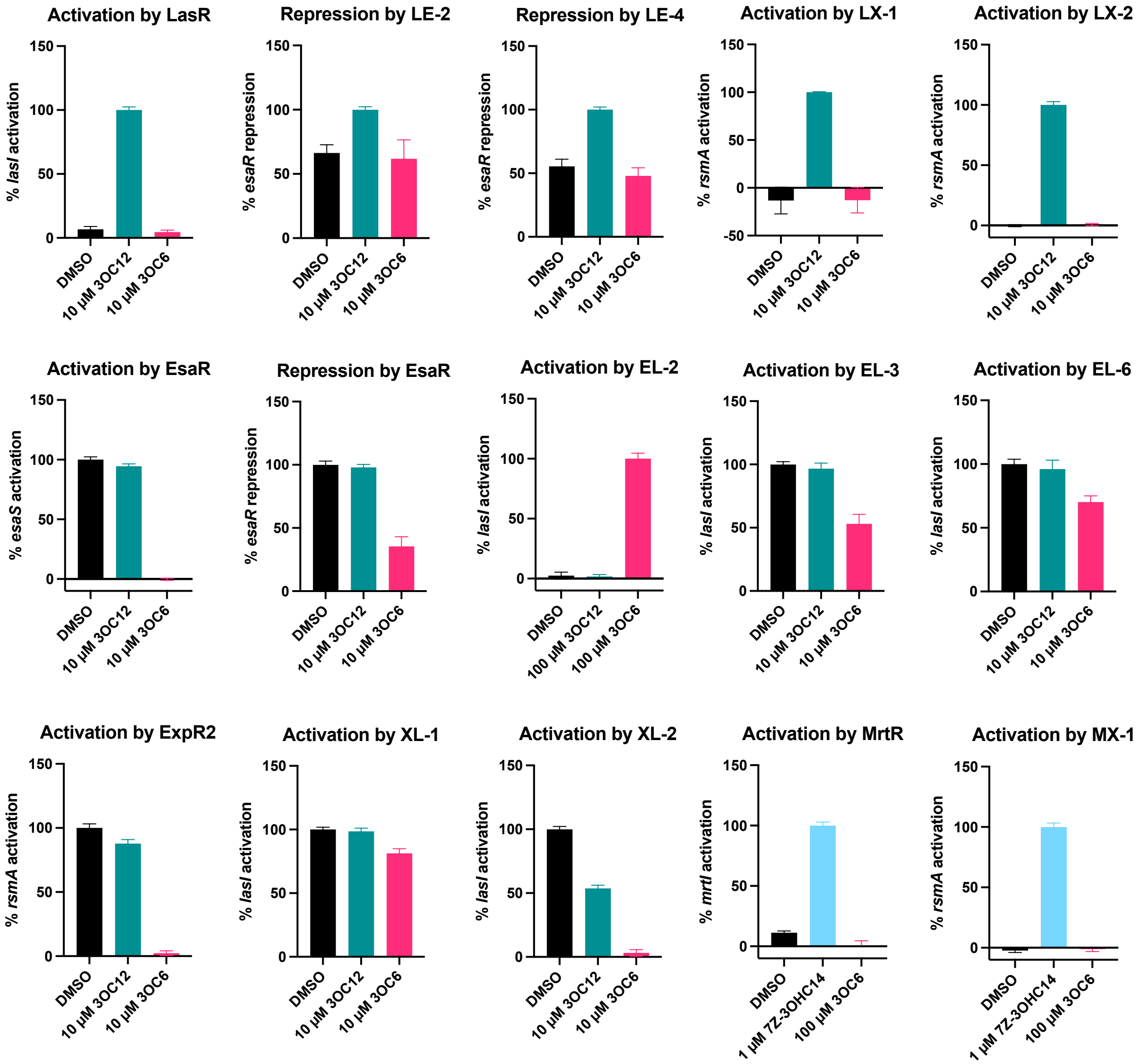


**Figure S3**. **Response of chimeras to ligands corresponding to the LBD vs. DBD receptor.** All data were collected in the same *E. coli* JLD271 reporter strains as the data shown in **Figure 3**. Data represent at least three biological replicates, each performed in technical triplicate, except for the "Activation by MrtR" data, which represents two biological replicates. Error bars indicate standard deviation. Data are normalized to uninduced culture (0%) and the maximum activity of each construct (100%).

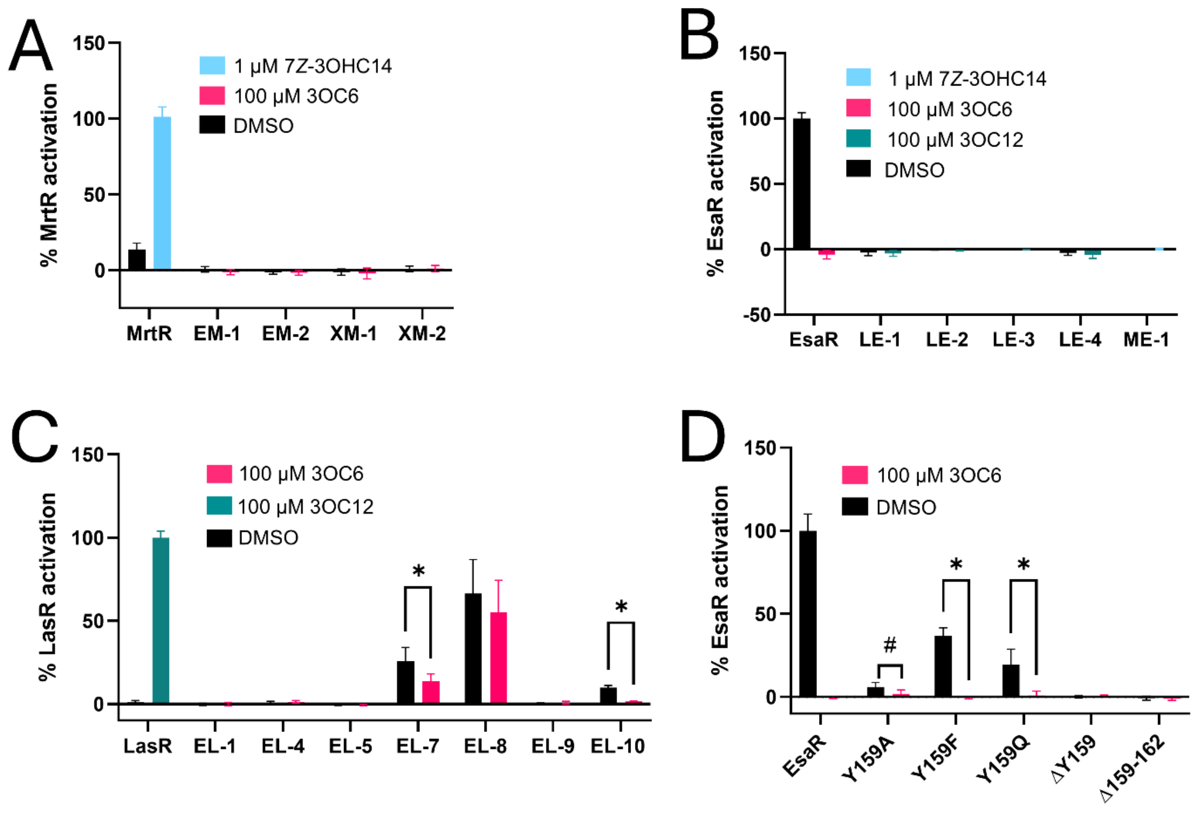


**Figure S4. Additional chimera and mutant activity data. (A)** Transcriptional activation of the *mrtI* promoter by chimeras with an MrtR DBD in an *E. coli* JLD271 reporter, normalized to uninduced culture (0%) and WT MrtR saturated with 7*Z*-3OHC14 (100%). Data represent at least four biological replicates, each performed in technical triplicate. WT MrtR activity is graphed for reference. **(B)** Transcriptional activation of the *esaS* promoter by chimeras with an EsaR DBD in an *E. coli* JLD271 reporter, normalized to uninduced culture (0%) and maximum EsaR activation (100%). Data represent at least two biological replicates, each performed in technical triplicate. WT EsaR activity is graphed for reference. **(C)** Transcriptional activation of the *lasI* promoter by chimeras with a LasR DBD (those not shown in **Figure 3B**) in a JLD271 reporter, normalized to uninduced culture (0%) and LasR saturated with 3OC12 (100%). Data represent at least three biological replicates, each performed in technical triplicate. WT LasR activity is graphed for reference. **(D)** Transcriptional activation of the *esaS* promoter by EsaR mutants in a BW27749 reporter, normalized to uninduced culture (0%) and maximum EsaR activation (100%). Data represent at least three biological replicates, each performed in technical triplicate. WT EsaR activity is graphed for reference. **(All graphs)** Error bars indicate standard deviation. Asterisks and hash marks indicate that mutant activity is different with and without ligand with $p \le 0.01$ or $p \le 0.05$, respectively, based on multiple unpaired t-tests with the Holm-Šídák correction for multiple comparisons.

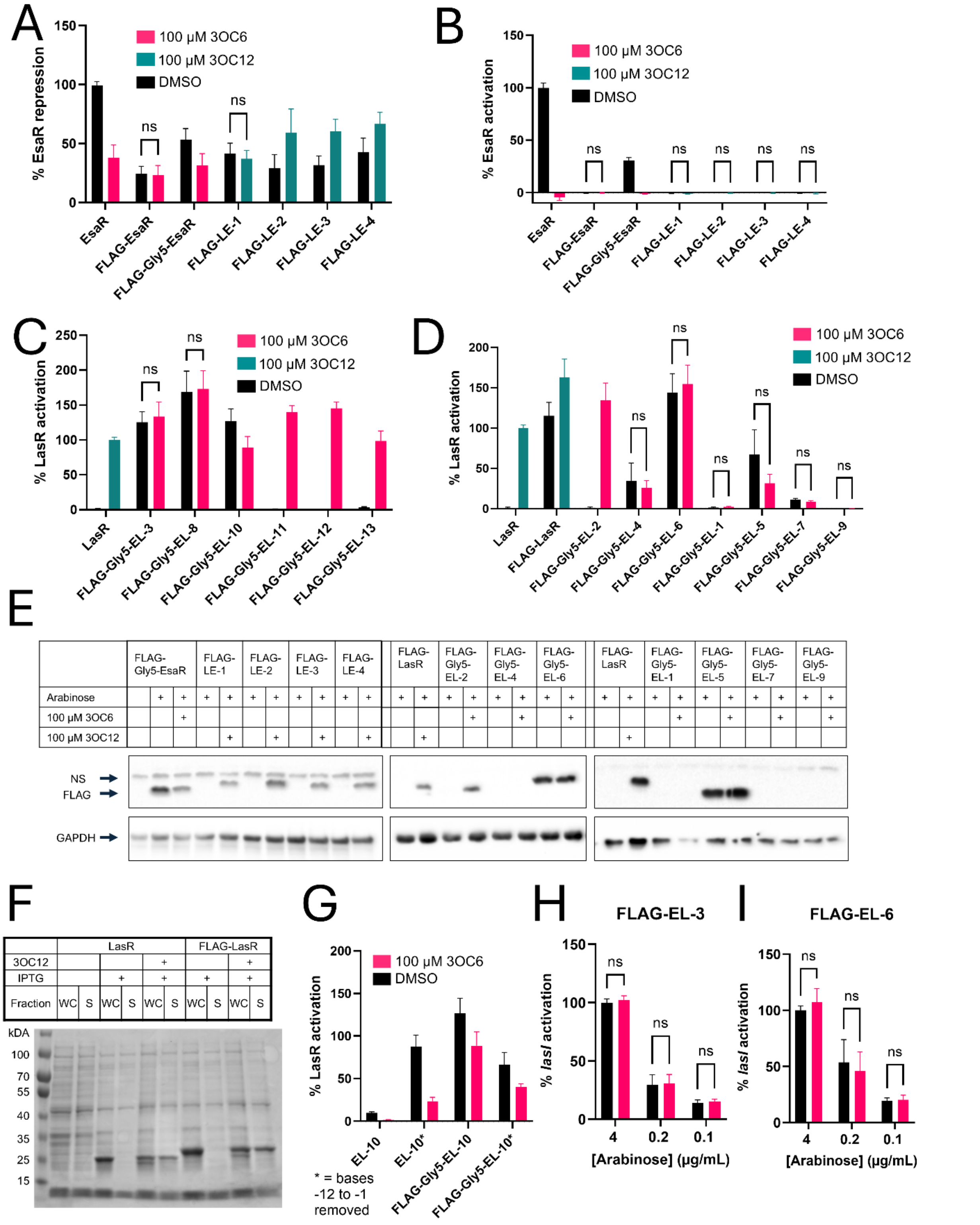


**Figure S5. Effects of FLAG tag on activity.** **(A)** Transcriptional repression of the *esaR* promoter by FLAG-tagged constructs with an EsaR DBD in an *E. coli* JLD271 reporter, normalized to uninduced culture (0%) and maximum EsaR repression (100%). Data represent at least two biological replicates, each performed in technical triplicate. WT EsaR activity is graphed for reference. **(B)** Transcriptional activation of the *esaS* promoter by FLAG-tagged constructs with an EsaR DBD in a JLD271 reporter, normalized to uninduced culture (0%) and maximum EsaR activation (100%). Data represent at least two biological replicates, each performed in technical triplicate. WT EsaR activity is graphed for reference. **(C)** and **(D)** Transcriptional activation of the *lasI* promoter by FLAG-tagged constructs with a LasR DBD in a JLD271 reporter, normalized to uninduced culture (0%) and LasR saturated with 3OC12 (100%). Data represent at least three biological replicates, each performed in technical duplicate or triplicate. WT LasR activity is graphed for reference. **(E)** Western blots of whole cell lysate from reporter strains used to collect data shown in panels **(B)** and **(D)**. Images are representative of two biological replicates. See **Figure S11A** for additional replicates and end of this SI document for full blot images. Note that nonspecific binding to GFP, the production of which is induced by FLAG-Gly5-EsaR in the absence of ligand, could affect the intensity of the FLAG-Gly5-EsaR bands. **(F)** Solubility test of LasR and FLAG-LasR overexpressed in *E. coli* BL21(DE3) in the presence and absence of 10 μM 3OC12. WC represents whole cell lysate and S represents soluble fraction. **(G)** Transcriptional activation of the *lasI* promoter in a JLD271 reporter by chimera EL-10 with and without a FLAG tag and/or 12 base pairs removed just upstream of the start codon, normalized to uninduced culture (0%) and LasR saturated with 3OC12 (100%). Data represent at least three biological replicates, each performed in technical triplicate. Data for EL-10 and FLAG-Gly5-EL10 are also shown in **Figure S4C** and **S5C**, respectively. Compound key applies to panels **G**, **H,** and **I**. **(H)** Transcriptional activation of the *lasI* promoter by FLAG-EL-3 in the presence of different concentrations of arabinose, normalized to FLAG-EL-3 activation with 4 μg/mL arabinose and no compound (DMSO) (100%). Data represent two biological replicates, each performed in technical triplicate. Activity was not significantly different with and without ligand even with a cutoff of $p \leq 0.1$. **(I)** Transcriptional activation of the *lasI* promoter by FLAG-EL-6 in the presence of different concentrations of arabinose, normalized to FLAG-EL-6 activation with 4 μg/mL arabinose and no compound (DMSO) (100%). Data represent two biological replicates, each performed in technical triplicate. Activity was not significantly different with and without ligand even with a cutoff of $p \leq 0.1$. **(All graphs)** Error bars indicate standard deviation. Unless indicated as ns (not significant), activity was significantly different with and without ligand based on multiple t-tests with the Holm-Šídák correction for multiple comparisons and $p \leq 0.01$.

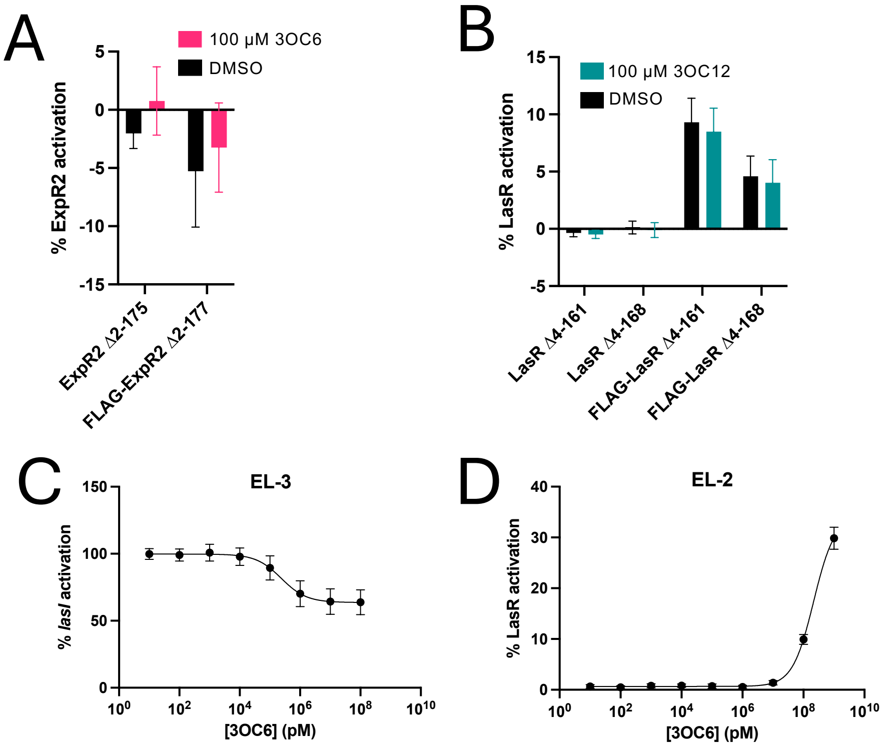


**Figure S6. Activity of DBDs alone. (A)** Transcriptional activation of the *rsmA* promoter by ExpR2 truncations in an *E. coli* JLD271 reporter, normalized to uninduced culture (0%) and maximum ExpR2 activation (100%). **(B)** Transcriptional activation of the *lasI* promoter by LasR truncations in a JLD271 reporter, normalized to uninduced culture (0%) and LasR saturated with 3OC12 (100%).
**(C)** Transcriptional activation of the *lasI* promoter by chimera EL-3, normalized to uninduced culture (0%) and mutant activity with no compound (DMSO) (100%). **(D)** Transcriptional activation of the *lasI* promoter by chimera EL-2, normalized to uninduced culture (0%) and LasR saturated with 3OC12 (100%). **(All graphs)** Data represent at least three biological replicates, each performed in technical triplicate. Error bars indicate standard deviation.

A

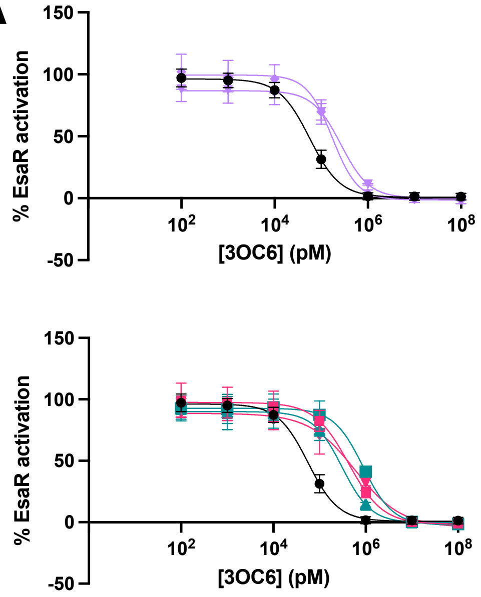


| Construct | $EC_{50}$ (95% CI) (nM) |
|---|---|
| EsaR | 56 (49-64) |
| Δ4: 168-171 (RAPA) | 260 (200-340) |
| Δ4: 172-175 (LNQS) | 170 (140-210) |
| Δ6: 164-169 (TEGERA) | 880 (710-1100) |
| Δ6: 171-176 (ALNQSA) | 310 (240-400) |
| Δ8: 164-171 (TEGERAPA) | 420 (330-540) |
| Δ8: 168-169, 171-176 (RA, ALNQSA) | 600 (430-820) |

B

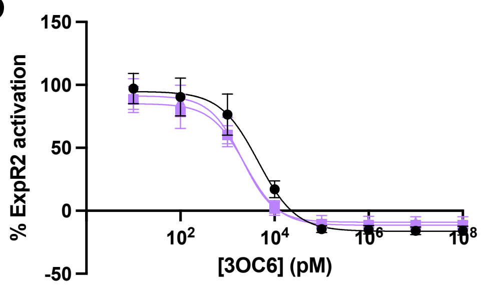


| Construct | $EC_{50}$ (95% CI) (nM) |
|---|---|
| ExpR2 | 4.5 (3.4-5.8) |
| Δ169-172 (PQSR) | 2.4 (2.0-2.8) |
| Δ173-176 (KQGD) | 2.0 (1.7-2.5) |

**Figure S7. Ligand potency in EsaR and ExpR2 linker deletion mutants. (A)** Dose-response curves and $EC_{50}$ values for EsaR linker mutants. Graphs show transcriptional activation of the *esaS* promoter in an *E. coli* BW27749 reporter, normalized to uninduced culture (0%) and maximum EsaR activation (100%). WT EsaR activity is plotted on both graphs for comparison. **(B)** Dose-response curves and $EC_{50}$ values for ExpR2 linker mutants showing transcriptional activation of the *rsmA* promoter in an *E. coli* JLD271 reporter, normalized to uninduced culture (0%) and maximum ExpR2 activation (100%). WT ExpR2 activity is plotted for comparison. **(All graphs)** Data represent at least three biological replicates, each performed in technical triplicate. Error bars indicate standard deviation. CI = confidence interval. Some data from dose-response curves are also included in on/off plots in **Figure 4**.

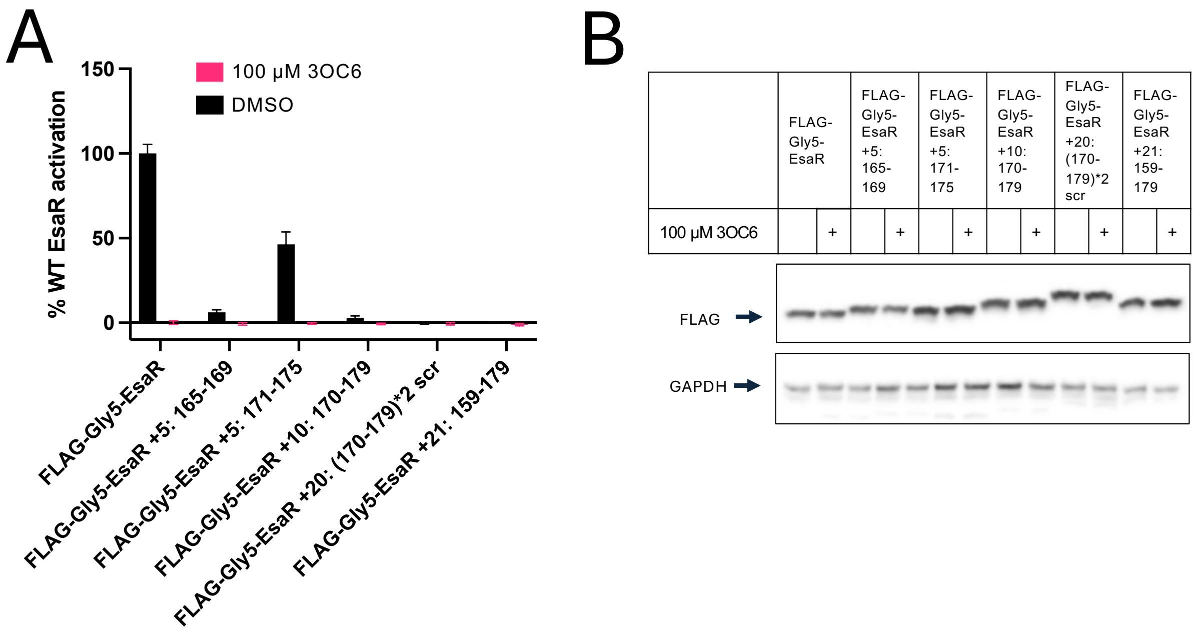


**Figure S8. Activity and accumulation of FLAG-tagged EsaR linker addition mutants.**
**(A)** Transcriptional activation of the *esaS* promoter in an *E. coli* BW27749 reporter induced with 0.4 mg/mL arabinose, normalized to uninduced culture (0%) and FLAG-Gly5-EsaR activation with DMSO (100%). Data represent at least three biological replicates, each performed in technical triplicate. Error bars indicate standard deviation. **(B)** Western blots of whole cell lysate from reporter strains used to collect data shown in panel **A**. Images are representative of two biological replicates. Additional replicate is shown in **Figure S11B**, and full blot images are shown at the end of this SI document.

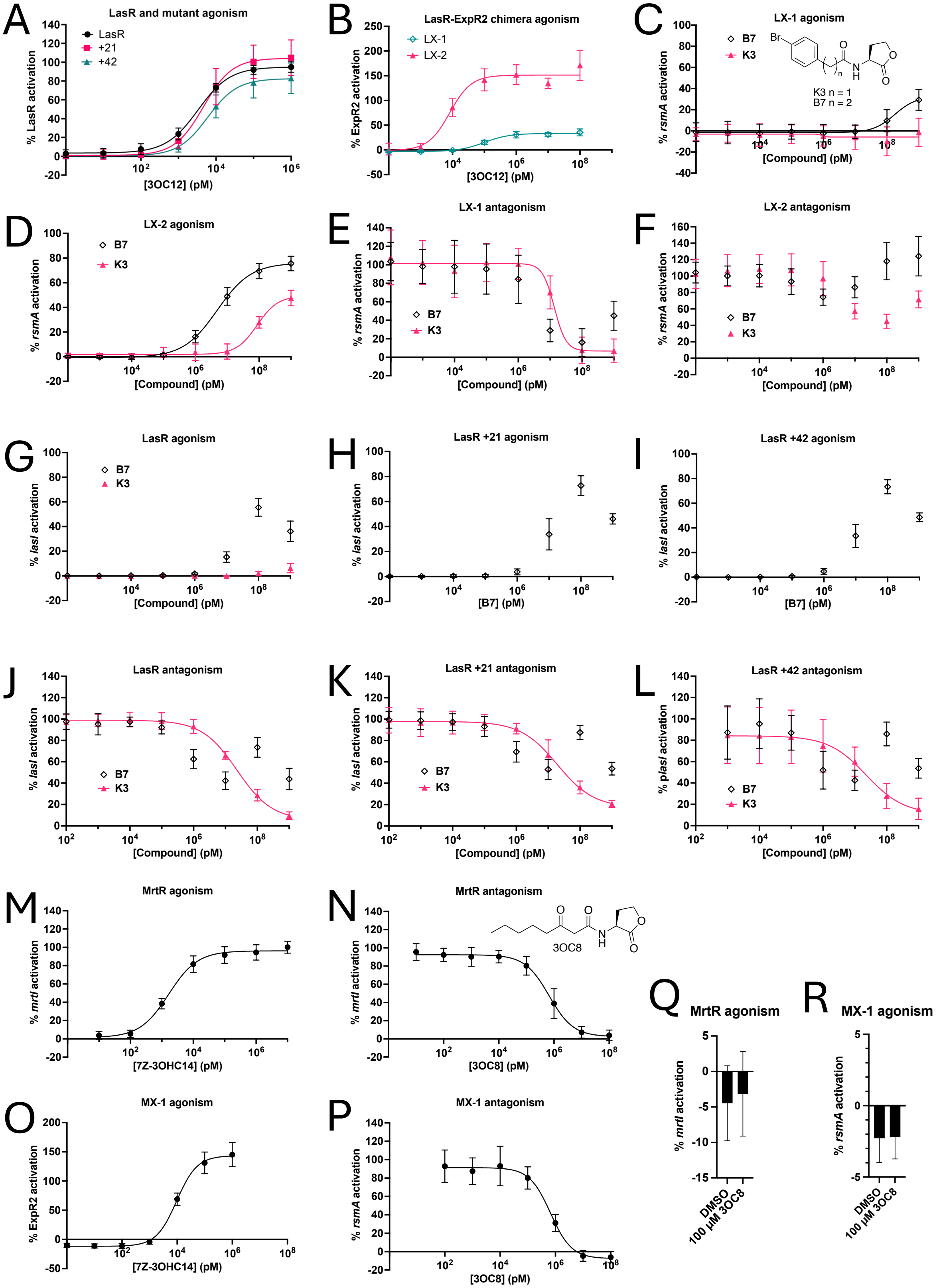
A
LasR and mutant agonism
LasR
+21
+42
% LasR activation
[3OC12] (pM)
B
LasR-ExpR2 chimera agonism
LX-1
LX-2
% ExpR2 activation
[3OC12] (pM)
C
LX-1 agonism
B7
K3
Br
K3 n = 1
B7 n = 2
% rsmA activation
[Compound] (pM)
D
LX-2 agonism
B7
K3
% rsmA activation
[Compound] (pM)
E
LX-1 antagonism
B7
K3
% rsmA activation
[Compound] (pM)
F
LX-2 antagonism
B7
K3
% rsmA activation
[Compound] (pM)
G
LasR agonism
B7
K3
% lasI activation
[Compound] (pM)
H
LasR +21 agonism
% lasI activation
[B7] (pM)
I
LasR +42 agonism
% lasI activation
[B7] (pM)
J
LasR antagonism
B7
K3
% lasI activation
[Compound] (pM)
K
LasR +21 antagonism
B7
K3
% lasI activation
[Compound] (pM)
L
LasR +42 antagonism
B7
K3
% plasI activation
[Compound] (pM)
M
MrtR agonism
% mrtI activation
[7Z-3OHC14] (pM)
N
MrtR antagonism
3OC8
% mrtI activation
[3OC8] (pM)
O
MX-1 agonism
% ExpR2 activation
[7Z-3OHC14] (pM)
P
MX-1 antagonism
% rsmA activation
[3OC8] (pM)
Q
MrtR agonism
% mrtI activation
DMSO
100 µM 3OC8
R
MX-1 agonism
% rsmA activation
DMSO
100 µM 3OC8

**Figure S9. Small molecule inhibition of select mutants and chimeras** (*figure on previous page)***.**
**(A)** Transcriptional activation of the *lasI* promoter by LasR linker extensions in an *E. coli* BW27749 reporter, normalized to uninduced culture (0%) and LasR saturated with 3OC12 (100%). See **Table S2** for sequence details. **(B)** Transcriptional activation of the *rsmA* promoter by chimeras with an ExpR2 DBD in an *E. coli* JLD271 reporter, normalized to uninduced culture (0%) and maximum ExpR2 activation (100%). Data represent at least two biological replicates, each performed in technical triplicate. **(C)** and **(D)** Transcriptional activation of the *rsmA* promoter by the LasR-ExpR2 chimeras LX-1 **(C)** and LX-2 **(D)** in a JLD271 reporter, normalized to DMSO (0%) and each mutant saturated with 3OC12 (100%). **(E)** Transcriptional activation of the *rsmA* promoter by chimera LX-1 in a JLD271 reporter, normalized to chimera LX-1 with DMSO (0%) and 100 nM 3OC12 (100%). Compounds were competed against 100 nM 3OC12. **(F)** Transcriptional activation of the *rsmA* promoter by LX-2 in a JLD271 reporter, normalized to LX-2 with DMSO (0%) and 10 nM 3OC12 (100%). Compounds were competed against 10 nM 3OC12. **(G-I)** Transcriptional activation of the *lasI* promoter by LasR **(G)**, LasR +21 **(H)**, and LasR +42 **(I)** in a BW27749 reporter, normalized to each construct with DMSO (0%) and saturated with 3OC12 (100%). **(J-L)** Transcriptional activation of the *lasI* promoter by LasR **(J)**, LasR +21 **(K)**, and LasR +42 **(L)** in a BW27749 reporter, normalized to each construct with DMSO (0%) and 10 nM 3OC12 (100%). Compounds were competed against 10 nM 3OC12. **B7** data collected in technical duplicate or triplicate. **(M)** and **(Q)** Transcriptional activation of the *mrtI* promoter by MrtR in a BW27749 reporter, normalized to uninduced culture (0%) and MrtR saturated with 7*Z*-3OHC14 (100%). **(N)** Transcriptional activation of the *mrtI* promoter by MrtR in a BW27749 reporter, normalized to uninduced culture (0%) and MrtR with 8 nM 7*Z*-3OHC14 (100%). Compound was competed against 8 nM 7*Z*-3OHC14. **(O)** Transcriptional activation of the *rsmA* promoter by MrtR-ExpR2 chimera MX-1 in a JLD271 reporter, normalized to uninduced culture (0%) and maximum ExpR2 activation (100%). **(P)** Transcriptional activation of the *rsmA* promoter by chimera MX-1 in a JLD271 reporter, normalized to uninduced culture (0%) and MX-1 with 40 nM 7*Z*-3OHC14 (100%). Compound was competed against 40 nM 7*Z*-3OHC14. **(R)** Transcriptional activation of the *rsmA* promoter by chimera MX-1 in a JLD271 reporter, normalized to uninduced culture (0%) and MX-1 with 1 μM 7*Z*-3OHC14 (100%). **(All graphs)** Unless indicated, data represent at least three biological replicates, each performed in technical triplicate. Error bars indicate standard deviation. Some data from panels **A** and **B** are also shown in **Figures 4** and **3**, respectively.

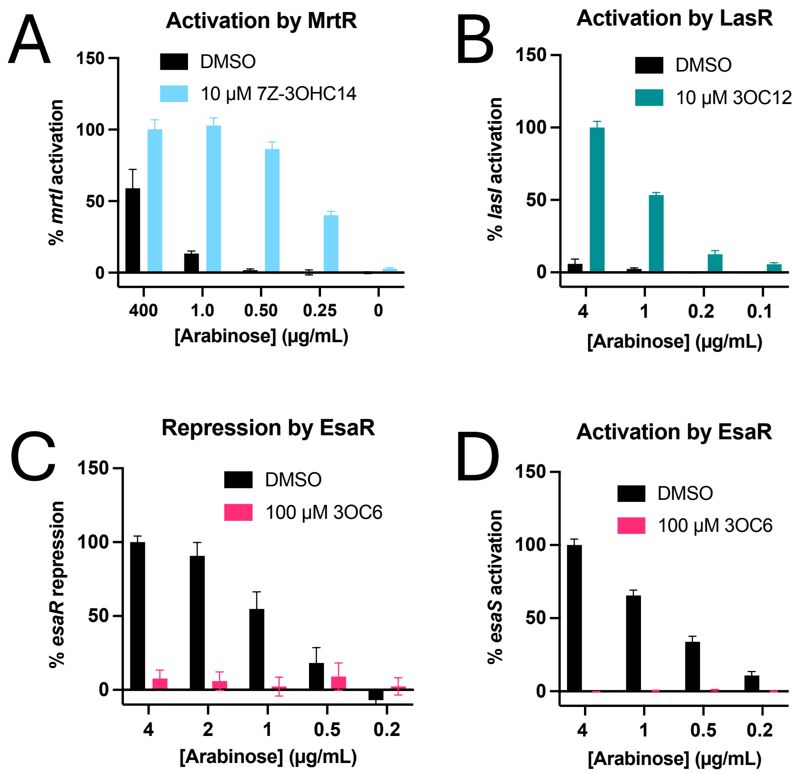


**Figure S10. Optimization of arabinose concentration for *E. coli* BW27749 reporters.**
**(A)** Transcriptional activation of the *mrtI* promoter by MrtR in a BW27749 reporter, normalized to uninduced culture (0%) and 400 µg/mL arabinose with 7Z-3OHC14 (100%). Data represent at least two biological replicates, each performed in technical triplicate. **(B)** Transcriptional activation of the *lasI* promoter by LasR in a BW27749 reporter, normalized to uninduced culture (0%) and 4 µg/mL arabinose with 3OC12 (100%). Data represent two biological replicates, each performed in technical triplicate. **(C)** Transcriptional repression of the *esaR* promoter by EsaR in a BW27749 reporter, normalized to uninduced culture (0%) and 4 µg/mL arabinose with DMSO (100%). Data represent three biological replicates, each performed in technical triplicate. **(D)** Transcriptional activation of the *esaS* promoter by EsaR in a BW27749 reporter, normalized to uninduced culture (0%) and 4 µg/mL arabinose with DMSO (100%). Data represent at least two biological replicates, each performed in technical triplicate. **(All graphs)** Error bars indicate standard deviation.

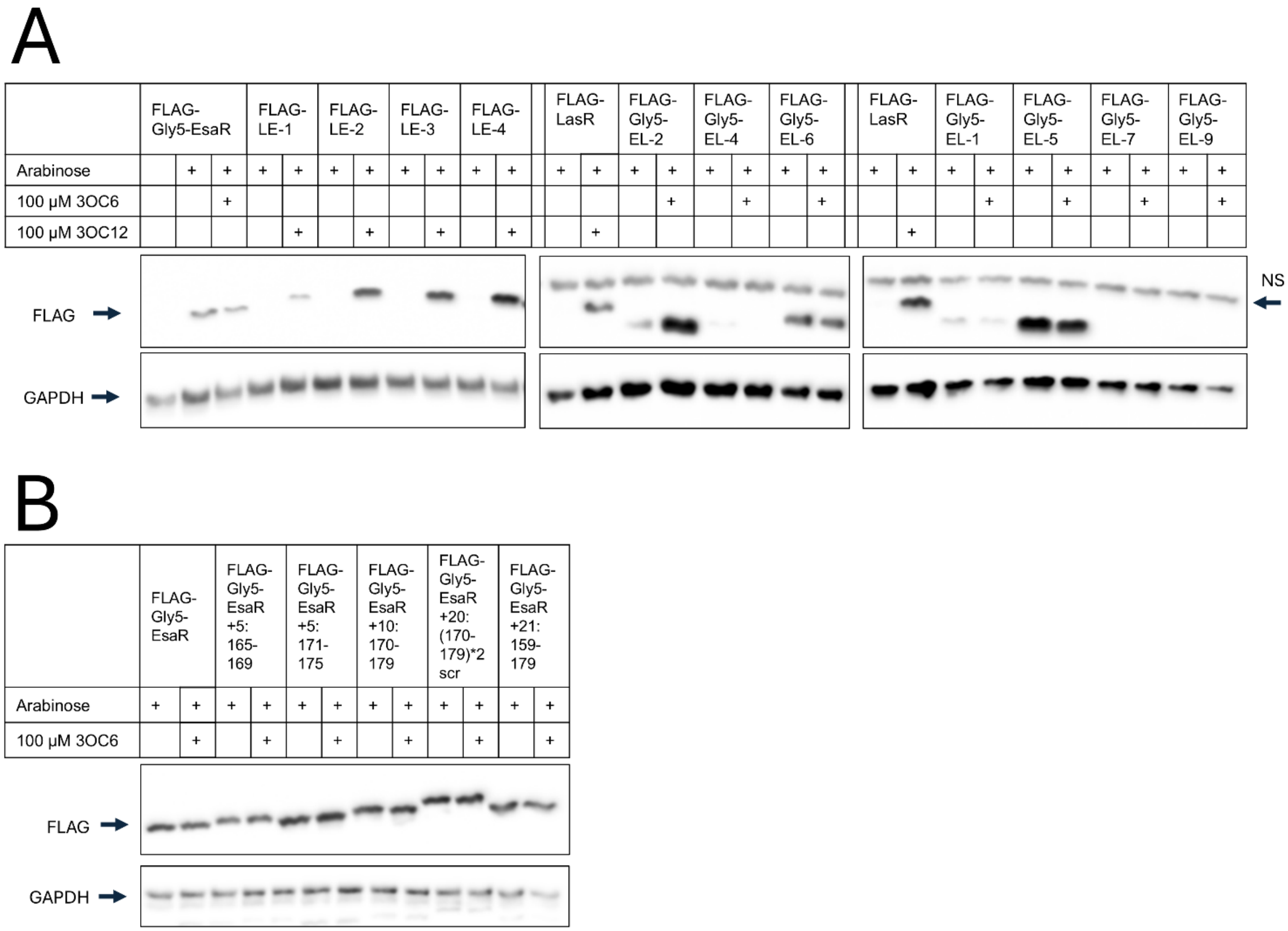


**Figure S11**. Additional biological replicate of western blots in **Figure S5E (A)** and **Figure S8B (B)**. “NS” indicates nonspecific band observed in FLAG blot.

**Table S1**. **Protein sequences of WT LuxR-type receptors examined in this study.**

| Protein | Sequence |
|---|---|
| **LasR** | MALVDGFLELERSSGKLEWSAILQKMASDLGFSKILFGLLPKDSQDYENAFIVGNYPAAWRE<br>HYDRAGYARVDPTVSHCTQSVLPIFWEPSIYQTRKQHEFFEEASAAGLVYGLTMPLHGARG<br>ELGALSLSVEAENRAEANRFMESVLPTLWMLKDYALQSGAGLAFEHPVSKPVVLTSREKEV<br>LQWCAIGKTSWEISVICNCSEANVNFHMGNIRRKFGVTSRRVAAIMAVNLGLITL |
| **EsaR** | MFSFFLENQTITDTLQTYIQRKLSPLGSPDYAYTVVSKKNPSNVLIISSYPDEWIRLYRANNFQ<br>LTDPVILTAFKRTSPFAWDENITLMSDLRFTKIFSLSKQYNIVNGFTYVLHDHMNNLALLSVIIK<br>GNDQTALEQRLAAEQGTMQMLLIDFNEQMYRLAGTEGERAPALNQSADKTIFSSRENEVLY<br>WASMGKTYAEIAAITGISVSTVKFHIKNVVVKLGVSNARQAIRLGVELDLIRPAASAAR |
| **ExpR2** | MSVFCSDNEIINNTIKSYLGRKLKQYGDLKYAYMIMNKKNPSQVVIISNYPQEWVNTYKENN<br>YQHIDPVILTAINKVSPFSWEDNIVINSKLKFSKIFNLSKEYDIVNGYTFVLHDNNNNLAALSIM<br>FEEHAPTDMENIVEENKDKLQMLLIAVHEKITTLYKEMTQSPQSRKQGDKEIFSQRENEILYW<br>ASMGKTYPEIALILDIKISTVKFHIGNVVKKLGVLNAKHAIRLGVELQLIKPEPL |
| **MrtR** | MIENTYSEKFESAFEQIKAAANVDAAIRILQAEYGLDFVTYHLAQTIAAKIDSPFVRTTYPDAW<br>VSRYLLNSYVKVDPIVKQGFERQLPFDWSEVEPTPEAYAMLVDAQKHGIGGNGYSIPVADK<br>AQRRALLSLNARIPAEEWAELVRRCRNEWIEIAHLIHRKAVYELHGENDPVPALSPREIECLH<br>WTALGKDYKDISVILGISEHTTRDYLKTARFKLGCATISAAASRAVQLRIINP |

**Table S2**. **Bacterial strains and plasmids used in this study.** $Ap^R$ = ampicillin resistance, $Gm^R$ = gentamycin resistance, $Cm^R$ = chloramphenicol resistance.

| **Strain** | **Description** | **Reference** |
|---|---|---|
| *E. coli* Top 10 | Used for cloning. | Lab collection |
| *E. coli* JLD271 | K12 *ΔsdiA*$_{ec}$, used for reporters. | (18) |
| *E. coli* BW27749 | Constitutive expression of low-affinity arabinose pump AraE and deletion of *araFGH*. Used for arabinose-titratable reporters. | (2) |
| *E. coli* BL21(DE3) pLysS | Used for protein overexpression. pLysS confers $Cm^R$. | Novagen |
| **Plasmid** | **Description** | **Reference** |
| pJN105L | Arabinose-inducible lasR expression vector; pBBRMCS backbone ($Gm^R$) | (14) |
| pSC11L | Broad host range p*lasI*-lacZ reporter ($Ap^R$) | (19) |
| pJN105-esaR | pJN105 with arabinose-inducible promoter controlling *esaR* ($Gm^R$) | (20) |
| pesaR-AC:GFP | p*esaS* promoter from *Pantoea stewartii* subsp. *stewartii* controlling GFP ($Ap^R$) | (9) |
| pesaR:GFP | p*esaR* promoter from *P. stewartii* controlling *GFP* ($Ap^R$) | (20) |
| pJN105-expR2 | pJN105 with arabinose-inducible promoter controlling *expR2* ($Gm^R$) | (20) |
| prsmA:GFP | *rsmA* promoter from *Pectobacterium versatile* Ecc71 controlling *GFP* ($Ap^R$) | (20) |
| pJN105-mrtR | pJN105 with arabinose-inducible promoter controlling *mrtR* ($Gm^R$) | This study |
| pmrtI:GFP | *mrtI* promoter (-120 to -1 relative the *mrtI* translational start site) from *Mesorhizobium tianshanense* controlling *GFP* ($Ap^R$) | This study |
| pET17b-lasR | Vector for LasR overexpression in BL21(DE3) ($Ap^R$) | (21) |
| pET17b-FLAG-lasR[a] | pET17b-lasR with a FLAG tag inserted between residues 1 and 2 ($Ap^R$) | This study |
| pJN105-EL-1 | pJN105L with LasR residues 1-151 replaced with EsaR 1-147 ($Gm^R$) | This study |
| pJN105-EL-2 | pJN105L with LasR residues 1-161 replaced with EsaR 1-158 ($Gm^R$) | This study |
| pJN105-EL-3 | pJN105L with LasR residues 1-168 replaced with EsaR 1-168 ($Gm^R$) | This study |
| pJN105-EL-4 | pJN105L with LasR residues 1-177 replaced with EsaR 1-181 ($Gm^R$) | This study |
| pJN105-EL-5 | pJN105L with LasR residues 1-168 replaced with EsaR 1-163 ($Gm^R$) | This study |

| | | |
|---|---|---|
| pJN105-EL-6 | pJN105L with LasR residues 1-168 replaced with EsaR 1-179 ($Gm^R$) | This study |
| pJN105-EL-7 | pJN105L with LasR residues 1-161 replaced with EsaR 1-162 ($Gm^R$) | This study |
| pJN105-EL-8 | pJN105L with LasR residues 1-161 replaced with EsaR 1-168 ($Gm^R$) | This study |
| pJN105-EL-9 | pJN105L with LasR residues 1-135 replaced with EsaR 1-131 ($Gm^R$) | This study |
| pJN105-EL-10 | pJN105L with LasR residues 1-172 replaced with EsaR 1-168 ($Gm^R$) | This study |
| pJN105-EL-10* | pJN105-EL-10 with base pairs -12 to -1 removed ($Gm^R$) | This study |
| pJN105-EL-11 | pJN105L with LasR residues 1-157 replaced with EsaR 1-158 ($Gm^R$) | This study |
| pJN105-EL-12 | pJN105L with LasR residues 1-160 replaced with EsaR 1-158 ($Gm^R$) | This study |
| pJN105-EL-13 | pJN105L with LasR residues 1-165 replaced with EsaR 1-158 ($Gm^R$) | This study |
| pJN105-XL-1 | pJN105L with LasR residues 1-161 replaced with ExpR2 1-158 ($Gm^R$) | This study |
| pJN105-XL-2 | pJN105L with LasR residues 1-172 replaced with ExpR2 1-168 ($Gm^R$) | This study |
| pJN105-LE-1 | pJN105-esaR with EsaR residues 1-147 replaced with LasR 1-151 ($Gm^R$) | This study |
| pJN105-LE-2 | pJN105-esaR with EsaR residues 1-158 replaced with LasR 1-161 ($Gm^R$) | This study |
| pJN105-LE-3 | pJN105-esaR with EsaR residues 1-168 replaced with LasR 1-168 ($Gm^R$) | This study |
| pJN105-LE-4 | pJN105-esaR with EsaR residues 1-181 replaced with LasR 1-177 ($Gm^R$) | This study |
| pJN105-ME-1 | pJN105-esaR with EsaR residues 1-172 replaced with MrtR 1-174 ($Gm^R$) | This study |
| pJN105-LX-1 | pJN105-expR2 with ExpR2 residues 1-158 replaced with LasR 1-161 ($Gm^R$) | This study |
| pJN105-LX-2 | pJN105-expR2 with ExpR2 residues 1-168 replaced with LasR 1-172 ($Gm^R$) | This study |
| pJN105-MX-1 | pJN105-expR2 with ExpR2 residues 1-169 replaced with MrtR 1-174 ($Gm^R$) | This study |
| pJN105-XM-1 | pJN105-mrtR with MrtR residues 1-165 replaced with ExpR2 1-158 ($Gm^R$) | This study |
| pJN105-XM-2 | pJN105-mrtR with MrtR residues 1-174 replaced with ExpR2 1-170 ($Gm^R$) | This study |

| | | |
|---|---|---|
| pJN105-EM-1 | pJN105-mrtR with MrtR residues 1-174 replaced with EsaR 1-168 ($Gm^R$) | This study |
| pJN105-EM-2 | pJN105-mrtR with MrtR residues 1-170 replaced with EsaR 1-168 ($Gm^R$) | This study |
| pJN105-FLAG-LasR[a] | pJN105L with a FLAG tag inserted between residues 1 and 2 ($Gm^R$) | This study |
| pJN105-FLAG-Gly5-EL1[b] | pJN105-EL-1 with a FLAG-Gly5 tag inserted between residues 1 and 2 ($Gm^R$) | This study |
| pJN105-FLAG-Gly5-EL2[b] | pJN105-EL-2 with a FLAG-Gly5 tag inserted between residues 1 and 2 ($Gm^R$) | This study |
| pJN105-FLAG-Gly5-EL3[b] | pJN105-EL-3 with a FLAG-Gly5 tag inserted between residues 1 and 2 ($Gm^R$) | This study |
| pJN105-FLAG-Gly5-EL4[b] | pJN105-EL-4 with a FLAG-Gly5 tag inserted between residues 1 and 2 ($Gm^R$) | This study |
| pJN105-FLAG-Gly5-EL5[b] | pJN105-EL-5 with a FLAG-Gly5 tag inserted between residues 1 and 2 ($Gm^R$) | This study |
| pJN105-FLAG-Gly5-EL6[b] | pJN105-EL-6 with a FLAG-Gly5 tag inserted between residues 1 and 2 ($Gm^R$) | This study |
| pJN105-FLAG-Gly5-EL7[b] | pJN105-EL-7 with a FLAG-Gly5 tag inserted between residues 1 and 2 ($Gm^R$) | This study |
| pJN105-FLAG-Gly5-EL8[b] | pJN105-EL-8 with a FLAG-Gly5 tag inserted between residues 1 and 2 ($Gm^R$) | This study |
| pJN105-FLAG-Gly5-EL9[b] | pJN105-EL-9 with a FLAG-Gly5 tag inserted between residues 1 and 2 ($Gm^R$) | This study |
| pJN105-FLAG-Gly5-EL10[b] | pJN105-EL-10 with a FLAG-Gly5 tag inserted between residues 1 and 2 ($Gm^R$) | This study |
| pJN105-FLAG-Gly5-EL10*[b] | pJN105-FLAG-Gly5-EL10 with base pairs -12 to -1 deleted ($Gm^R$) | This study |
| pJN105-FLAG-Gly5-EL11[b] | pJN105-EL-11 with a FLAG-Gly5 tag inserted between residues 1 and 2 ($Gm^R$) | This study |
| pJN105-FLAG-Gly5-EL12[b] | pJN105-EL-12 with a FLAG-Gly5 tag inserted between residues 1 and 2 ($Gm^R$) | This study |
| pJN105-FLAG-Gly5-EL13[b] | pJN105-EL-13 with a FLAG-Gly5 tag inserted between residues 1 and 2 ($Gm^R$) | This study |
| pJN105-lasR +21 | pJN105L with EsaR residues 159-179 inserted after LasR 168 ($Gm^R$) | This study |
| pJN105-lasR +42 | pJN105L with EsaR residues 159-179 doubled and inserted after LasR 168 ($Gm^R$) | This study |
| pJN105-lasR +105 | pJN105L with an insertion after LasR 168 of EsaR residues 159-168, (EsaR 159-179)*4 scrambled,[c] EsaR 169-179 ($Gm^R$) | This study |
| pJN105-lasR ΔP170 | pJN105L with a Δ170 mutation ($Gm^R$) | This study |
| pJN105-lasR ΔV171 | pJN105L with a Δ171 mutation ($Gm^R$) | This study |

| | | |
|---|---|---|
| pJN105-lasR ΔS172 | pJN105L with a Δ172 mutation (Gm$^R$) | This study |
| pJN105-lasR ΔK173 | pJN105L with a Δ173 mutation (Gm$^R$) | This study |
| pJN105-lasR Δ169, 171 | pJN105L with Δ169 and Δ171 mutations (Gm$^R$) | This study |
| pJN105-lasR Δ172-173 | pJN105L with a Δ172-173 mutation (Gm$^R$) | This study |
| pJN105-lasR Δ169, 171-172 | pJN105L with Δ169 and Δ171-172 mutations (Gm$^R$) | This study |
| pJN105-lasR Δ170-172 | pJN105L with a Δ170-172 mutation (Gm$^R$) | This study |
| pJN105-lasR Scr 169-174 | pJN105L with residues H169-P174 scrambled (HPVSKP to SPKPVH) (Gm$^R$) | This study |
| pJN105-lasR Δ4-161 | pJN105-lasR with residues 4 to 161 deleted (Gm$^R$) | This study |
| pJN105-lasR Δ4-168 | pJN105-lasR with residues 4 to 168 deleted (Gm$^R$) | This study |
| pJN105-FLAG-lasR Δ4-161[a] | pJN105-lasR Δ4-161 with a FLAG tag inserted between residues 3 and 4 of the mutant (Gm$^R$) | This study |
| pJN105-FLAG-lasR Δ4-168[a] | pJN105-lasR Δ4-168 with a FLAG tag inserted between residues 3 and 4 of the mutant (Gm$^R$) | This study |
| pJN105-Myc-lasR | pJN105-lasR with a Myc tag (EQKLISEEDL) inserted between residues 1 and 2 (Gm$^R$) | This study |
| pJN105-His6-lasR | pJN105-lasR with 6 His residues (HHHHHH) inserted between residues 1 and 2 (Gm$^R$) | This study |
| pJN105-HA-lasR | pJN105-lasR with an HA tag (YPYDVPDYA) inserted between residues 1 and 2 (Gm$^R$) | This study |
| pJN105-Gly6-lasR | pJN105-lasR with 6 Gly residues (GGGGGG) inserted between residues 1 and 2 (Gm$^R$) | This study |
| pJN105-NLS-lasR | pJN105-lasR with an NLS tag (PKKKRKV) inserted between residues 1 and 2 (Gm$^R$) | This study |
| pJN105-lasR Δ2-10 | pJN105L with residues 2-10 deleted (Gm$^R$) | This study |
| pJN105-FLAG-esaR[a] | pJN105-esaR with a FLAG tag inserted between residues 1 and 2 (Gm$^R$) | This study |
| pJN105-FLAG-Gly5-esaR[b] | pJN105-esaR with a FLAG-Gly5 tag inserted between residues 1 and 2 (Gm$^R$) | This study |
| pJN105-FLAG-LE-1[a] | pJN105-LE-1 with a FLAG tag inserted between residues 1 and 2 (Gm$^R$) | This study |
| pJN105-FLAG-LE-2[a] | pJN105-LE-2 with a FLAG tag inserted between residues 1 and 2 (Gm$^R$) | This study |
| pJN105-FLAG-LE-3[a] | pJN105-LE-3 with a FLAG tag inserted between residues 1 and 2 (Gm$^R$) | This study |
| pJN105-FLAG-LE-4[a] | pJN105-LE-4 with a FLAG tag inserted between residues 1 and 2 (Gm$^R$) | This study |
| pJN105-esaR +5: 165-169 | pJN105-esaR with residues 165-169 (EGERA) doubled (Gm$^R$) | This study |
| pJN105-esaR +5: 171-175 | pJN105-esaR with residues 171-175 (ALNQS) doubled (Gm$^R$) | This study |

| | | |
|---|---|---|
| pJN105-esaR +5G at 171 | pJN105-esaR with 5 glycine residues inserted after residue 171 ($Gm^R$) | This study |
| pJN105-esaR +5G at 176 | pJN105-esaR with 5 glycine residues inserted after residue 176 ($Gm^R$) | This study |
| pJN105-esaR +5A at 171 | pJN105-esaR with 5 alanine residues inserted after residue 171 ($Gm^R$) | This study |
| pJN105-esaR +5A at 176 | pJN105-esaR with 5 alanine residues inserted after residue 176 ($Gm^R$) | This study |
| pJN105-esaR +10: 170-179 | pJN105-esaR with residues 170-179 (PALNQSADKT) doubled ($Gm^R$) | This study |
| pJN105-esaR +20: (170-179)*2 scr | pJN105-esaR with ALSTQPSKDNLPNQAATKDA (Residues 170-179 (PALNQSADKT) doubled and scrambled) inserted after residue 169 ($Gm^R$) | This study |
| pJN105-esaR +21: 159-179 | pJN105-esaR with residues 159-179 (YRLAGTEGERAPALNQSADKT) doubled ($Gm^R$) | This study |
| pJN105-esaR Δ164-167 | pJN105-esaR with residues 164-167 (TEGE) deleted ($Gm^R$) | This study |
| pJN105-esaR Δ168-171 | pJN105-esaR with residues 168-171 (RAPA) deleted ($Gm^R$) | This study |
| pJN105-esaR Δ172-175 | pJN105-esaR with residues 172-175 (LNQS) deleted ($Gm^R$) | This study |
| pJN105-esaR Δ176-179 | pJN105-esaR with residues 176-179 (ADKT) deleted ($Gm^R$) | This study |
| pJN105-esaR Δ164-169 | pJN105-esaR with residues 164-169 (TEGERA) deleted ($Gm^R$) | This study |
| pJN105-esaR Δ171-176 | pJN105-esaR with residues 171-176 (ALNQSA) deleted ($Gm^R$) | This study |
| pJN105-esaR Δ168-169, 171-176 | pJN105-esaR with residues 168-169 (RA) and 171-176 (ALNQSA) deleted ($Gm^R$) | This study |
| pJN105-esaR Δ164-171 | pJN105-esaR with residues 164-171 (TEGERAPA) deleted ($Gm^R$) | This study |
| pJN105-esaR Δ164-175 | pJN105-esaR with residues 164-175 (TEGERAPALNQS) deleted ($Gm^R$) | This study |
| pJN105-esaR Scr 172-177 | pJN105-esaR with residues 172-177 (LNQSAD) scrambled to DQLASN ($Gm^R$) | This study |
| pJN105-esaR ΔY159 | pJN105-esaR with a deletion of Y159 ($Gm^R$) | This study |
| pJN105-esaR Δ159-162 | pJN105-esaR with a deletion of residues 159-162 ($Gm^R$) | This study |
| pJN105-esaR Y159A | pJN105-esaR with a Y159A mutation ($Gm^R$) | This study |
| pJN105-esaR Y159F | pJN105-esaR with a Y159F mutation ($Gm^R$) | This study |
| pJN105-esaR Y159Q | pJN105-esaR with a Y159F mutation ($Gm^R$) | This study |
| pJN105-esaR G163A | pJN105-esaR with a G163A mutation ($Gm^R$) | This study |

| | | |
|---|---|---|
| pJN105-esaR T164A | pJN105-esaR with a T164A mutation (Gm$^R$) | This study |
| pJN105-esaR E165A | pJN105-esaR with a E165A mutation (Gm$^R$) | This study |
| pJN105-esaR G166A | pJN105-esaR with a G166A mutation (Gm$^R$) | This study |
| pJN105-esaR E167A | pJN105-esaR with a E167A mutation (Gm$^R$) | This study |
| pJN105-esaR R168A | pJN105-esaR with a R168A mutation (Gm$^R$) | This study |
| pJN105-esaR P170A | pJN105-esaR with a P170A mutation (Gm$^R$) | This study |
| pJN105-esaR L172A | pJN105-esaR with a L172A mutation (Gm$^R$) | This study |
| pJN105-esaR N173A | pJN105-esaR with a N173A mutation (Gm$^R$) | This study |
| pJN105-esaR Q174A | pJN105-esaR with a Q174A mutation (Gm$^R$) | This study |
| pJN105-esaR S175A | pJN105-esaR with a S175A mutation (Gm$^R$) | This study |
| pJN105-esaR D177A | pJN105-esaR with a D177A mutation (Gm$^R$) | This study |
| pJN105-esaR K178A | pJN105-esaR with a K178A mutation (Gm$^R$) | This study |
| pJN105-esaR T179A | pJN105-esaR with a T179A mutation (Gm$^R$) | This study |
| pJN105-esaR I180A | pJN105-esaR with a I180AA mutation (Gm$^R$) | This study |
| pJN105-FLAG-Gly5-esaR +5: 165-169[b] | pJN105-esaR +5: 165-169 with a FLAG-Gly5 tag inserted between residues 1 and 2 (Gm$^R$) | This study |
| pJN105-FLAG-Gly5-esaR +5: 171-175[b] | pJN105-esaR +5: 171-175 with a FLAG-Gly5 tag inserted between residues 1 and 2 (Gm$^R$) | This study |
| pJN105-FLAG-Gly5-esaR +10: 170-179[b] | pJN105-esaR +10: 170-179 with a FLAG-Gly5 tag inserted between residues 1 and 2 (Gm$^R$) | This study |
| pJN105-FLAG-Gly5-esaR +20: (170-179)*2 scr[b] | pJN105-esaR +20: (170-179)*2 scr with a FLAG-Gly5 tag inserted between residues 1 and 2 (Gm$^R$) | This study |
| pJN105-FLAG-Gly5-esaR +21: 159-179[b] | pJN105-esaR +21: 159-179 with a FLAG-Gly5 tag inserted between residues 1 and 2 (Gm$^R$) | This study |
| pJN105-esaR F4A | pJN105-esaR with an F4A mutation (Gm$^R$) | This study |
| pJN105-esaR F5A | pJN105-esaR with an F5A mutation (Gm$^R$) | This study |
| pJN105-mrtR +4G at 171 | pJN105-mrtR with 4 glycine residues inserted after residue 171 (Gm$^R$) | This study |
| pJN105-mrtR +4G at 175 | pJN105-mrtR with 4 glycine residues inserted after residue 175 (Gm$^R$) | This study |
| pJN105-mrtR +4A at 171 | pJN105-mrtR with 4 alanine residues inserted after residue 171 (Gm$^R$) | This study |
| pJN105-mrtR +4A at 175 | pJN105-mrtR with 4 alanine residues inserted after residue 175 (Gm$^R$) | This study |
| pJN105-mrtR +5A at 171 | pJN105-mrtR with 5 alanine residues inserted after residue 171 (Gm$^R$) | This study |
| pJN105-mrtR +5A at 175 | pJN105-mrtR with 5 alanine residues inserted after residue 175 (Gm$^R$) | This study |
| pJN105-mrtR Δ171-173 | pJN105-mrtR with residues 171-173 (GEN) deleted (Gm$^R$) | This study |

| | | |
|---|---|---|
| pJN105-expR2 +5G at 171 | pJN105-expR2 with 5 glycine residues inserted after residue 171 ($Gm^R$) | This study |
| pJN105-expR2 +5G at 176 | pJN105-expR2 with 5 glycine residues inserted after residue 176 ($Gm^R$) | This study |
| pJN105-expR2 +5A at 171 | pJN105-expR2 with 5 alanine residues inserted after residue 171 ($Gm^R$) | This study |
| pJN105-expR2 +5A at 176 | pJN105-expR2 with 5 alanine residues inserted after residue 176 ($Gm^R$) | This study |
| pJN105-expR2 +5: 172-176 | pJN105-expR2 with residues 172-176 (RKQGD) doubled ($Gm^R$) | This study |
| pJN105-expR2 +10: 168-177 | pJN105-expR2 with residues 168-177 (SPQSRKQGDK) doubled ($Gm^R$) | This study |
| pJN105-expR2 +21:158-178 | pJN105-expR2 with residues 158-178 (ITTLYKEMTQSPQSRKQGDKE) doubled ($Gm^R$) | This study |
| pJN105-expR2 Δ169-172 | pJN105-expR2 with residues 169-172 (PQSR) deleted ($Gm^R$) | This study |
| pJN105-expR2 Δ173-176 | pJN105-expR2 with residues 173-176 (KQGD) deleted ($Gm^R$) | This study |
| pJN105-expR2 Δ170-175 | pJN105-expR2 with residues 170-175 (QSRKQG) deleted ($Gm^R$) | This study |
| pJN105-expR2 Δ169-176 | pJN105-expR2 with residues 169-176 (PQSRKQGD) deleted ($Gm^R$) | This study |
| pJN105-expR2 K177A | pJN105-expR2 with a K177A mutation ($Gm^R$) | This study |
| pJN105-expR2 E178A | pJN105-expR2 with an E178A mutation ($Gm^R$) | This study |
| pJN105-expR2 F4A | pJN105-expR2 with an F4A mutation ($Gm^R$) | This study |
| pJN105-expR2 Δ2-175 | pJN105-expR2 with residues 2-175 deleted ($Gm^R$) | This study |
| pJN105-FLAG-expR2 Δ2-177[a] | pJN105-expR2 with residues 2-177 deleted and a FLAG tag added between residues 1 and 2 of the mutant[d] ($Gm^R$) | This study |

[a] Sequence of FLAG tag: GATTATAAAGATGATGATGATAAA.

[b] Sequence of FLAG-Gly5 tag: GATTATAAAGATGATGATGATAAAGGTGGTGGTGGTGGC.

[c] Amino acid sequence of (EsaR 159-179)*4 scrambled that was inserted to make LasR +105 (see **Table S6** for gBlock): PYTTEARDRNLADLTGETNLKQASYATRGESGEYQRAKDAKGAGAARAAPGQPLALA RNPLEATELKYESLARTATRDNSQGEG.

[d] Analogous to ExpR2 Δ2-175 because the last 2 residues of the FLAG tag replace native D176 and K177.

**Table S3.** Ordinary one-way ANOVA with Šídák's correction for multiple comparisons for activation by EsaR +5 linker mutants with DMSO, corresponding to data in **Figure 4F**.

| Comparison | Mean diff. | 99.00% CI of diff. | Adjusted p value |
|---|---|---|---|
| **+5A at 171 vs. +5G at 171** | -21 | -36 to -5 | <0.001 |
| **+5A at 171 vs. +5A at 176** | -23 | -39 to -8 | <0.001 |
| **+5G at 171 vs. +5G at 176** | -13 | -28 to 2 | 0.038 |
| **+5A at 176 vs. +5G at 176** | -10 | -26 to 5 | 0.134 |

**Table S4.** Ordinary one-way ANOVA with Šídák's correction for multiple comparisons for activation by ExpR2 +5 linker mutants with DMSO, corresponding to data in **Figure 4G**.

| Comparison | Mean diff. | 99.00% CI of diff. | Adjusted p value |
|---|---|---|---|
| **+5A at 171 vs. +5G at 171** | -28 | -41 to -16 | <0.001 |
| **+5A at 171 vs. +5A at 176** | 30 | 17 to 42 | <0.001 |
| **+5G at 171 vs. +5G at 176** | 44 | 31 to 58 | <0.001 |
| **+5A at 176 vs. +5G at 176** | -14 | -27 to -0.2 | 0.008 |

**Table S5.** Ordinary one-way ANOVA with Šídák's correction for multiple comparisons for activation by MrtR +4 linker mutants with 7Z-3OHC14, corresponding to data in **Figure 4B**.

| Comparison | Mean diff. | 99.00% CI of diff. | Adjusted p value |
|---|---|---|---|
| **+4G at 171 vs. +4A at 171** | -5 | -15 to 6 | 0.474 |
| **+4G at 171 vs. +4G at 175** | 23 | 12 to 33 | <0.001 |
| **+4A at 171 vs. +4A at 175** | 27 | 17 to 37 | <0.001 |
| **+4G at 175 vs. +4A at 175** | -0.4 | -11 to 10 | 0.999 |

**Table S6. gBlocks used in this study.**

| gBlock | Sequence |
|---|---|
| *mrtI* promoter fragment used for pmrtI:GFP | CTGCACACTTTTGCGCGATATGCGCCCCCCTCATCTGAGGGGGCCCATCTGAGG<br>GAATTTCCGAACCGGCCCGCTTGAACCATTCTGCTTTCCACGAACTTGAAAACGC<br>TGGAGGGCAAA |
| Codon-optimized *mrtR* sequence | ATGATCGAAAACACATATTCGGAGAAGTTCGAGAGTGCGTTCGAGCAAATCAAAG<br>CTGCTGCGAATGTCGATGCCGCAATTCGTATCCTTCAGGCAGAATATGGGTTGGA<br>TTTCGTCACGTATCACTTGGCGCAAACCATCGCCGCCAAAATCGACTCGCCATTC<br>GTTCGCACTACATATCCTGATGCGTGGGTCAGCCGCTACCTGCTGAACAGCTATG<br>TCAAGGTGGACCCAATCGTGAAGCAAGGATTTGAACGCCAATTGCCTTTTGATTG<br>GAGTGAAGTGGAGCCCACACCTGAGGCGTACGCCATGCTGGTCGATGCACAGAA<br>ACACGGAATCGGAGGTAACGGATACTCTATTCCAGTTGCCGACAAGGCCCAACG<br>TCGCGCATTACTGTCGTTAAACGCTCGTATCCCTGCCGAAGAGTGGGCGGAGTT<br>GGTACGTCGTTGTCGCAACGAATGGATTGAGATTGCACATTTGATTCATCGCAAA<br>GCGGTTTACGAGTTGCACGGGGAGAACGATCCGGTCCCCGCCCTTTCGCCTCGT<br>GAAATCGAATGCCTGCATTGGACTGCCCTTGGTAAGGATTACAAGGATATCTCAG<br>TGATTTTAGGTATCAGTGAGCACACCACGCGTGATTACCTGAAAACTGCCCGTTTT<br>AAATTAGGTTGCGCGACCATCTCAGCAGCGGCCAGCCGTGCCGTTCAGCTTCGC<br>ATCATCAATCCGTAA |
| gBlock used to create pJN105-lasR +105 | GTGCCGGACTGGCCTTCGAATACCGCCTGGCAGGCACCGAAGGTGAACGACCTT<br>ACACTACGGAAGCTCGTGACCGTAACTTAGCAGACCTGACAGGTGAAACAAACCT<br>TAAGCAGGCTTCCTATGCTACCCGTGGAGAATCTGGGGAGTATCAGCGCGCAAA<br>AGACGCCAAGGGGGCAGGCGCGGCGCGTGCTGCGCCGGGCCAACCTCTTGCC<br>CTGGCTCGTAACCCTTTGGAAGCGACTGAATTAAAGTATGAGTCCCTTGCTCGTA<br>CAGCGACACGTGACAATAGCCAGGGTGAAGGTGCCCCGGCGTTAAATCAGAGCG<br>CGGACAAAACGCATCCGGTCAGCAAACCGGTG |

**Table S7. Primers used in this study.**

| Plasmid | Primer name | Sequence |
|---|---|---|
| pJN105-EL-1 | Vector (from pJN105L) FWD | CTGCCGAACAGGGCACGATGTGGATGCTCAAGGACTACGC |
| | Vector REV | TCAAGGAAGAAAGAGAACATAGCGCTACGTTCTTCTTAAGA |
| | Insert (from pJN105-esaR) FWD | CTTAAGAAGAACGTAGCGCTATGTTCTCTTTCTTCCTTGAAAA |
| | Insert REV | GCGTAGTCCTTGAGCATCCACATCGTGCCCTGTTCGGCAG |
| pJN105-LE-1 | Vector (from pJN105-esaR) FWD | AGTCGGTCCTGCCGACCCTGCAGATGCTGCTGATTGATTT |
| | Vector REV | AAACCGTCAACCAAGGCCATGAATTCGCTAGCCCAAAAAAACG |
| | Insert (from pET17b-lasR) FWD | TTTTTTGGGCTAGCGAATTCATGGCCTTGGTTGACGGTTT |
| | Insert REV | AAATCAATCAGCAGCATCTGCAGGGTCGGCAGGACCGACT |
| pJN105-EL-2 | Vector (from pJN105L) FWD | TTGATTTTAACGAGCAGATGGGTGCCGGACTGGCCTTCGA |
| | Vector REV | TCAAGGAAGAAAGAGAACATAGCGCTACGTTCTTCTTAAGA |
| | Insert (from pJN105-esaR) FWD | CTTAAGAAGAACGTAGCGCTATGTTCTCTTTCTTCCTTGAAAA |
| | Insert REV | TCGAAGGCCAGTCCGGCACCCATCTGCTCGTTAAAATCAATC |
| pJN105-LE-2 | Vector (from pJN105-esaR) FWD | AGGACTACGCACTGCAGAGCTACCGCCTGGCAGGCACCGA |
| | Vector REV | AAACCGTCAACCAAGGCCATGAATTCGCTAGCCCAAAAAAACG |
| | Insert (from pET17b-lasR) FWD | TTTTTTGGGCTAGCGAATTCATGGCCTTGGTTGACGGTTT |
| | Insert REV | TCGGTGCCTGCCAGGCGGTAGCTCTGCAGTGCGTAGTCCT |
| pJN105-EL-3 | Vector (from pJN105L) FWD | CAGGCACCGAAGGTGAACGACATCCGGTCAGCAAACCGGT |
| | Vector REV | TCAAGGAAGAAAGAGAACATAGCGCTACGTTCTTCTTAAG |
| | Insert (from pJN105-esaR) FWD | CTTAAGAAGAACGTAGCGCTATGTTCTCTTTCTTCCTTGAAAA |
| | Insert REV | ACCGGTTTGCTGACCGGATGTCGTTCACCTTCGGTGCCTG |

| | | |
|---|---|---|
| pJN105-LE-3 | Vector (from pJN105-esaR) FWD | GTGCCGGACTGGCCTTCGAAGCCCCGGCGTTAAATCAGAG |
| | Vector REV | AAACCGTCAACCAAGGCCATGAATTCGCTAGCCCAAAAAA |
| | Insert (from pET17b-lasR) FWD | TTTTTTGGGCTAGCGAATTCATGGCCTTGGTTGACGGTTT |
| | Insert REV | CTCTGATTTAACGCCGGGGCTTCGAAGGCCAGTCCGGCAC |
| pJN105-EL-4 | Vector (from pJN105L) FWD | GCGCGGACAAAACGATATTTACCAGCCGGGAGAAGGAAGT |
| | Vector REV | TCAAGGAAGAAAGAGAACATAGCGCTACGTTCTTCTTAAGA |
| | Insert (from pJN105-esaR) FWD | CTTAAGAAGAACGTAGCGCTATGTTCTCTTTCTTCCTTGAAAA |
| | Insert REV | ACTTCCTTCTCCCGGCTGGTAAATATCGTTTTGTCCGC |
| pJN105-LE-4 | Vector (from pJN105-esaR) FWD | TCAGCAAACCGGTGGTTCTGTCCTCGCGTGAAAATGAGGT |
| | Vector REV | AAACCGTCAACCAAGGCCATGAATTCGCTAGCCCAAAAAAACG |
| | Insert (from pET17b-lasR) FWD | TTTTTTGGGCTAGCGAATTCATGGCCTTGGTTGACGGTTT |
| | Insert REV | ACCTCATTTTCACGCGAGGACAGAACCACCGGTTTGCTGA |
| pJN105-EL-5 | Vector (from pJN105L) FWD | AGATGTACCGCCTGGCAGGCCATCCGGTCAGCAAACCG |
| | Vector REV | TCAAGGAAGAAAGAGAACATAGCGCTACGTTCTTCTTAAGA |
| | Insert (from pJN105-esaR) FWD | CTTAAGAAGAACGTAGCGCTATGTTCTCTTTCTTCCTTGAAAA |
| | Insert REV | ACCGGTTTGCTGACCGGATGGCCTGCCAGGCGGTACATC |
| pJN105-EL-6 | Vector (from pJN105L) FWD | ATCAGAGCGCGGACAAAACGCATCCGGTCAGCAAACCG |
| | Vector REV | TCAAGGAAGAAAGAGAACATAGCGCTACGTTCTTCTTAAGA |
| | Insert (from pJN105-esaR) FWD | CTTAAGAAGAACGTAGCGCTATGTTCTCTTTCTTCCTTGAAAA |
| | Insert REV | CGGTTTGCTGACCGGATGCGTTTTGTCCGCGCTCTGAT |
| pJN105-EL-7 | Vector (from pJN105L) FWD | AGCAGATGTACCGCCTGGCAGGTGCCGGACTGGCCTTCGA |

| | | |
|---|---|---|
| | Vector REV | TCAAGGAAGAAAGAGAACATAGCGCTACGTTCTTC TTAAGA |
| | Insert (from pJN105-esaR) FWD | CTTAAGAAGAACGTAGCGCTATGTTCTCTTTCTTC CTTGAAAA |
| | Insert REV | TCGAAGGCCAGTCCGGCACCTGCCAGGCGGTACA TCTGCT |
| pJN105-EL-8 | Vector (from pJN105L) FWD | CAGGCACCGAAGGTGAACGAGGTGCCGGACTGG CCTTCGA |
| | Vector REV | TCAAGGAAGAAAGAGAACATAGCGCTACGTTCTTC TTAAGA |
| | Insert (from pJN105-esaR) FWD | CTTAAGAAGAACGTAGCGCTATGTTCTCTTTCTTC CTTGAAAA |
| | Insert REV | TCGAAGGCCAGTCCGGCACCTCGTTCACCTTCGG TGCCTG |
| pJN105-EL-9 | Vector (from pJN105L) FWD | CCGTGATCATTAAAGGCAACAACCGGGCCGAGGC CAAC |
| | Vector REV | TCAAGGAAGAAAGAGAACATAGCGCTACGTTCTTC TTAAGA |
| | Insert (from pJN105-esaR) FWD | CTTAAGAAGAACGTAGCGCTATGTTCTCTTTCTTC CTTGAAAA |
| | Insert REV | CGGTTGGCCTCGGCCCGGTTGTTGCCTTTAATGA TCACGG |
| pJN105-EL-10* | Vector (from pJN105L) FWD | AGGCACCGAAGGTGAACGAAAACCGGTGGTTCTG ACCAG |
| | Vector REV | TCAAGGAAGAAAGAGAACATTTCTTAAGAATTCGC TAGCCC |
| | Insert (from pJN105-esaR) FWD | GGCTAGCGAATTCTTAAGAAATGTTCTCTTTCTTCC TTGAAAACCA |
| | Insert REV | CTGGTCAGAACCACCGGTTTTCGTTCACCTTCGGT GCCTG |
| pJN105-EL-10 | from pJN105-EL10* FWD | AGCGCTATGTTCTCTTTCTTCCTTG |
| | from pJN105-EL10* REV | ACGTTCTTCTTAAGAATTCGCTAGC |
| pJN105-EL-11 | Vector (from pJN105L) FWD | GATTTTAACGAGCAGATGGCACTGCAGAGCGGTG |
| | Vector REV | TCAAGGAAGAAAGAGAACATAGCGCTACGTTCTTC TTAAGA |
| | Insert (from pJN105-esaR) FWD | CTTAAGAAGAACGTAGCGCTATGTTCTCTTTCTTC CTTGAAAA |
| | Insert REV | GGCACCGCTCTGCAGTGCCATCTGCTCGTTAAAAT CAATC |

| | | |
|---|---|---|
| pJN105-EL-12 | Vector (from pJN105L) FWD | GATTTTAACGAGCAGATGAGCGGTGCCGGACTGG |
| | Vector REV | TCAAGGAAGAAAGAGAACATAGCGCTACGTTCTTC TTAAGA |
| | Insert (from pJN105-esaR) FWD | CTTAAGAAGAACGTAGCGCTATGTTCTCTTTCTTC CTTGAAAA |
| | Insert REV | GGCCAGTCCGGCACCGCTCATCTGCTCGTTAAAA TCAATC |
| pJN105-EL-13 | Vector (from pJN105L) FWD | GATTTTAACGAGCAGATGGCCTTCGAACATCCGGT C |
| | Vector REV | TCAAGGAAGAAAGAGAACATAGCGCTACGTTCTTC TTAAGA |
| | Insert (from pJN105-esaR) FWD | CTTAAGAAGAACGTAGCGCTATGTTCTCTTTCTTC CTTGAAAA |
| | Insert REV | GACCGGATGTTCGAAGGCCATCTGCTCGTTAAAAT CAATC |
| pJN105-XL-1 | Vector (from pJN105L) FWD | TTGCCGTTCATGAGAAAATCGGTGCCGGACTGGC CTTCGA |
| | Vector REV | TCAGAGCAAAATACAGACATAGCGCTACGTTCTTC TTAAGAATTCGC |
| | Insert (from pJN105-expR2) FWD | CTTAAGAAGAACGTAGCGCTATGTCTGTATTTTGC TCTGAC |
| | Insert REV | TCGAAGGCCAGTCCGGCACCGATTTTCTCATGAA CGGCAA |
| pJN105-XL-2 | Vector (from pJN105L) FWD | ACAAAGAGATGACGCAAAGCAAACCGGTGGTTCT GACCAG |
| | Vector REV | TCAGAGCAAAATACAGACATAGCGCTACGTTCTTC TTAAG |
| | Insert (from pJN105-expR2) FWD | CTTAAGAAGAACGTAGCGCTATGTCTGTATTTTGC TCTGAC |
| | Insert REV | CTGGTCAGAACCACCGGTTTGCTTTGCGTCATCTC TTTGT |
| pJN105-LX-1 | Vector (from pJN105-expR2) FWD | AGGACTACGCACTGCAGAGCACCACGCTTTACAA AGAGAT |
| | Vector REV | AAACCGTCAACCAAGGCCATGAATTCGCTAGCCC AAAAAAACG |
| | Insert (from pJN105L) FWD | TTTTTTGGGCTAGCGAATTCATGGCCTTGGTTGAC GGTTT |
| | Insert REV | ATCTCTTTGTAAAGCGTGGTGCTCTGCAGTGCGTA GTCCT |
| pJN105-LX-2 | Vector (from pJN105-expR2) FWD | CCTTCGAACATCCGGTCAGCCCGCAGAGCAGAAA ACAAGG |

| | | |
|---|---|---|
| | Vector REV | AAACCGTCAACCAAGGCCATGAATTCGCTAGCCCAAAAAAACG |
| | Insert (from pJN105L) FWD | TTTTTTGGGCTAGCGAATTCATGGCCTTGGTTGACGGTTT |
| | Insert REV | CCTTGTTTTCTGCTCTGCGGGCTGACCGGATGTTCGAAGG |
| pJN105-mrtR | Vector (from pJN105-esaR) FWD | CGCATCATCAATCCGTAATCTAGAGCGGCCGCCACC |
| | Vector REV | CGAATATGTGTTTTCGATCATGAATTCGCTAGCCCAAAAA |
| | Insert (from gBlock) FWD | TGGGCTAGCGAATTCATGATCGAAAACACATATTCGGAG |
| | Insert REV | GGTGGCGGCCGCTCTAGATTACGGATTGATGATGCGAAGC |
| pmrtI:GFP | Vector (from pesaR-AC:GFP) FWD | AAAACGCTGGAGGGCAAACCAGCACACTGGCGGCC |
| | Vector REV | TCGCGCAAAAGTGTGCAGAATTCTTGAAGACGAAAGGGCCT |
| | Insert (from gBlock) FWD | CTTTCGTCTTCAAGAATTCTGCACACTTTTGCGCGA |
| | Insert REV | CGGCCGCCAGTGTGCTGGTTTGCCCTCCAGCGTTTT |
| pJN105-ME-1 | Vector (from pJN105-esaR) FWD | TTGCACGGGGAGAACGATAATCAGAGCGCGGACAAA |
| | Vector REV | CGAATATGTGTTTTCGATCATGAATTCGCTAGCCCAAAAA |
| | Insert (from pJN105-mrtR) FWD | TGGGCTAGCGAATTCATGATCGAAAACACATATTCGGAG |
| | Insert REV | TTTGTCCGCGCTCTGATTATCGTTCTCCCCGTGCAA |
| pJN105-EM-1 | Vector (from pJN105-esaR) FWD | CGCATCATCAATCCGTAATCTAGAGCGGCCGCCACC |
| | Vector REV | CGAAAGGGCGGGGACCGGTCGTTCACCTTCGGTGCC |
| | Insert (from pJN105-mrtR) FWD | GGCACCGAAGGTGAACGACCGGTCCCCGCCCTTTCG |
| | Insert REV | GGTGGCGGCCGCTCTAGATTACGGATTGATGATGCGAAGC |
| pJN105-EM-2 | Vector (from pJN105-esaR) FWD | CGCATCATCAATCCGTAATCTAGAGCGGCCGCCACC |
| | Vector REV | GACCGGATCGTTCTCCCCTCGTTCACCTTCGGTGCC |

| | | |
|---|---|---|
| | Insert (from pJN105-mrtR) FWD | GGCACCGAAGGTGAACGAGGGGAGAACGATCCGGTC |
| | Insert REV | GGTGGCGGCCGCTCTAGATTACGGATTGATGATGCGAAGC |
| pJN105-MX-1 | Vector (from pJN105-expR) FWD | TTGCACGGGGAGAACGATCAGAGCAGAAAACAAGGTG |
| | Vector REV | CGAATATGTGTTTTCGATCATGAATTCGCTAGCCCAAAAA |
| | Insert (from pJN105-mrtR) FWD | TGGGCTAGCGAATTCATGATCGAAAACACATATTCGGAG |
| | Insert REV | ACCTTGTTTTCTGCTCTGATCGTTCTCCCCGTGCAA |
| pJN105-XM-1 | Vector (from pJN105-expR) FWD | CGCATCATCAATCCGTAATCTAGAGCGGCCGCCACC |
| | Vector REV | CCCGTGCAACTCGTAAACGATTTTCTCATGAACGGCAATGA |
| | Insert (from pJN105-mrtR) FWD | GCCGTTCATGAGAAAATCGTTTACGAGTTGCACGGG |
| | Insert REV | GGTGGCGGCCGCTCTAGATTACGGATTGATGATGCGAAGC |
| pJN105-XM-2 | Vector (from pJN105-expR) FWD | CGCATCATCAATCCGTAATCTAGAGCGGCCGCCACC |
| | Vector REV | CGAAAGGGCGGGGACCGGCTGCGGGCTTTGCGTCATCT |
| | Insert (from pJN105-mrtR) FWD | ATGACGCAAAGCCCGCAGCCGGTCCCCGCCCTTTCG |
| | Insert REV | GGTGGCGGCCGCTCTAGATTACGGATTGATGATGCGAAGC |
| pJN105-FLAG-lasR | From pJN105-lasR FWD | GATGATGATAAAGCCTTGGTTGACGG |
| | From pJN105-lasR REV | ATCTTTATAATCCATAGCGCTACGTTC |
| pJN105-FLAG-esaR | From pJN105-esaR FWD | GATGATGATAAATTCTCTTTCTTCCTTGAAAAC |
| | From pJN105-esaR REV | ATCTTTATAATCCATGAATTCGCTAGCC |
| pJN105-FLAG-Gly5-esaR: adding tag to WT and mutants | From pJN105-esaR or mutant FWD | ATAAAGGTGGTGGTGGTGGCTTCTCTTTCTTCCTTGAAAAC |
| | From pJN105-esaR or mutant REV | CATCATCATCTTTATAATCCATGAATTCGCTAGCC |
| | From pJN105-EL FWD | ATAAAGGTGGTGGTGGTGGCTTCTCTTTCTTCCTTGAAAAC |

| | | |
|---|---|---|
| Adding FLAG-Gly5 tag to all pJN105-EL chimeras | From pJN105-EL REV | CATCATCATCTTTATAATCCATAGCGCTACGTTC |
| Adding FLAG tag to pJN105-LE chimeras | From pJN105-LE FWD | GATGATGATAAAGCCTTGGTTGACGG |
| | From pJN105-LE REV | ATCTTTATAATCCATGAATTCGCTAGCC |
| pJN105-FLAG-Gly5-EL-10* | From pJN105-EL-10* FWD | ATAAAGGTGGTGGTGGTGGCTTCTCTTTCTTCCTTGAAAAC |
| | From pJN105-EL-10* REV | CATCATCATCTTTATAATCCATTTCTTAAGAATTCGCTAG |
| pJN105-lasR +21 | Vector (from pJN105-EL-6) FWD | GTGCCGGACTGGCCTTCGAATACCGCCTGGCAGGCACCGA |
| | Vector REV | AAACCGTCAACCAAGGCCATAGCGCTACGTTCTTCTTAAGAATTCGCT |
| | Insert (from pET17b-lasR) FWD | CTTAAGAAGAACGTAGCGCTATGGCCTTGGTTGACGGTTT |
| | Insert REV | TCGGTGCCTGCCAGGCGGTATTCGAAGGCCAGTCCGGCAC |
| pJN105-lasR +42 | Vector (from pJN105-lasR +21) FWD | TGGGTCTTATTACTCTCTGATCTTGCCTCTAGAGCGGCCG |
| | Vector REV | TCGGTGCCTGCCAGGCGGTACGTTTTGTCCGCGCTCTGAT |
| | Insert (from pJN105-lasR +21) FWD | ATCAGAGCGCGGACAAAACGTACCGCCTGGCAGGCACCGA |
| | Insert REV | CGGCCGCTCTAGAGGCAAGATCAGAGAGTAATAAGACCCAAATTAACGGCC |
| pJN105-lasR +105 | Vector (from pJN105L) FWD | CATCCGGTCAGCAAACCGGTG |
| | Vector REV | TTCGAAGGCCAGTCCGGCAC |
| | Insert (from gBlock) FWD | GTGCCGGACTGGCCTTCGAA |
| | Insert REV | CATCCGGTCAGCAAACCGGTG |
| pJN105-lasR ∆P170 | From pJN105L FWD | GTCAGCAAACCGGTGG |
| | From pJN105L REV | ATGTTCGAAGGCCAG |
| pJN105-lasR ∆V171 | From pJN105L FWD | AGCAAACCGGTGGTTCTGAC |
| | From pJN105L REV | CGGATGTTCGAAGGCCAGTC |
| pJN105-lasR ∆S172 | From pJN105L FWD | AAACCGGTGGTTCTGACCAG |
| | From pJN105L REV | GACCGGATGTTCGAAGGCCAG |
| pJN105-lasR ∆K173 | From pJN105L FWD | CCGGTGGTTCTGACCAGCCG |

| | | |
|---|---|---|
| | From pJN105L REV | GCTGACCGGATGTTCGAAGGC |
| pJN105-lasR Δ169, 171 | From pJN105L FWD | CCGAGCAAACCGGTGGTTCTGACC |
| | From pJN105L REV | TTCGAAGGCCAGTCCGGCAC |
| pJN105-lasR Δ172-173 | From pJN105L FWD | CCGGTGGTTCTGACCAGCCG |
| | From pJN105L REV | GACCGGATGTTCGAAGGCCAG |
| pJN105-lasR Δ169, 171-172 | From pJN105L FWD | CCGAAACCGGTGGTTCTG |
| | From pJN105L REV | TTCGAAGGCCAGTCC |
| pJN105-lasR Δ170-172 | From pJN105L FWD | AAACCGGTGGTTCTG |
| | From pJN105L REV | ATGTTCGAAGGCCAG |
| pJN105-lasR Scr 169-174 | From pJN105L FWD | CCCGTCCACGTGGTTCTGACCAGC |
| | From pJN105L REV | TTTAGGCGATTCGAAGGCCAGTCC |
| pJN105-lasR Δ4-161 | From pJN105L FWD | TCGAAGGCCAGTCCGGCACCCAAGGCCATAGCGCTACGTT |
| | From pJN105L REV | AACGTAGCGCTATGGCCTTGGGTGCCGGACTGGCCTTCGA |
| pJN105-lasR Δ4-168 | From pJN105L FWD | ACCGGTTTGCTGACCGGATGCAAGGCCATAGCGCTACGTT |
| | From pJN105L REV | AACGTAGCGCTATGGCCTTGCATCCGGTCAGCAAACCGGT |
| pJN105-FLAG-lasR Δ4-161 | FLAG-LasR Δ4-161 FWD | GATGATGATAAAGGTGCCGGACTGGC |
| | FLAG-LasR Δ4-161 REV | ATCTTTATAATCCAAGGCCATAGCGCTAC |
| pJN105-FLAG-lasR Δ4-168 | FLAG-LasR Δ4-168 FWD | GATGATGATAAACATCCGGTCAGCAAAC |
| | FLAG-LasR Δ4-168 REV | ATCTTTATAATCCAAGGCCATAGCGC |
| pJN105-Myc-LasR | From pJN105L FWD | AGCGAAGAAGATCTGGCCTTGGTTGACGGTTTTC |
| | From pJN105L REV | AATCAGTTTCTGTTCCATAGCGCTACGTTCTTC |
| pJN105-HA-lasR | From pJN105L FWD | TGCCGGATTATGCGGCCTTGGTTGACGGTTTTC |
| | From pJN105L REV | CATCATACGGATACATAGCGCTACGTTCTTC |
| pJN105-NLS-lasR | From pJN105L FWD | AACGCAAAGTGGCCTTGGTTGACGGTTTTC |
| | From pJN105L REV | TTTTTTTCGGCATAGCGCTACGTTCTTC |
| pJN105-His6-lasR | From pJN105L FWD | CACCACCACGCCTTGGTTGACGGTTTTC |
| | From pJN105L REV | ATGATGATGCATAGCGCTACGTTCTTC |
| pJN105-Gly6-lasR | From pJN105L FWD | GGTGGTGGTGGTGGCGCCTTGGTTGACGGTTTTC |

| | | |
|---|---|---|
| | From pJN105L REV | ACCCATAGCGCTACGTTCTTC |
| pJN105-lasR ∆2-10 | From pJN105L FWD | GAACGCTCAAGTGGAAAATTG |
| | From pJN105L REV | CATAGCGCTACGTTCTTC |
| pJN105-esaR +5: 165-169 | From pJN105-esaR FWD | AACGAGCCGAAGGTGAACGAGCCCCGG |
| | From pJN105-esaR REV | CACCTTCGGTGCCTGCCAGGCGGTA |
| pJN105-esaR +5: 171-175 | From pJN105-esaR FWD | ATCAGAGCGCGTTAAATCAGAGCGCG |
| | From pJN105-esaR REV | TTAACGCCGGGGCTCGTTCACCTTC |
| pJN105-esaR +5G at 171 | From pJN105-esaR FWD | GTGGTGGCTTAAATCAGAGCGCGGACAAAAC |
| | From pJN105-esaR REV | CGCCACCCGCCGGGGCTCGTTC |
| pJN105-esaR +5G at 176 | From pJN105-esaR FWD | GTGGTGGCGACAAAACGATATTTTCCTC |
| | From pJN105-esaR REV | CGCCACCCGCGCTCTGATTTAAC |
| pJN105-esaR +5A at 171 | From pJN105-esaR FWD | CAGCCGCAGCGTTAAATCAGAGCGCG |
| | From pJN105-esaR REV | CCGCTGCCGGGGCTCGTTCACC |
| pJN105-esaR +5A at 176 | From pJN105-esaR FWD | CAGCCGCAGCGGACAAAACGATATTTTC |
| | From pJN105-esaR REV | CCGCTGCGCTCTGATTTAACGCC |
| pJN105-esaR +10: 170-179 | From pJN105-esaR FWD | CCGATAAAACGCCGGCGTTAAATCAG |
| | From pJN105-esaR REV | CTGATTGATTAAGGGCCGGGGCTCGTTCACCTTC |
| pJN105-esaR +20: (170-179)*2 Scr | From pJN105-esaR FWD | CAAGCCGCCACCAAGGACGCCCCGGCGTTAAATCAG |
| | From pJN105-esaR REV | ATTGGGAAGGTTGTCCTTAGATGGCTGCGTAGACAGCGCGGCTCGTTCACCTTC |
| pJN105-esaR +21: 159-179 | Vector from pJN105-esaR FWD | CCACCGCGGTGGAGCTCCAATTCGCCCTATAGTGAGTCGT |
| | Vector REV | TCGGTGCCTGCCAGGCGGTACGTTTTGTCCGCGCTCTGAT |
| | Insert (from pJN105-esaR) FWD | ATCAGAGCGCGGACAAAACGTACCGCCTGGCAGGCACCGA |

| | | |
|---|---|---|
| | Insert REV | ACGACTCACTATAGGGCGAATTGGAGCTCCACCG CGGTGG |
| pJN105-esaR Δ164-167 | From pJN105-esaR FWD | CGAGCCCCGGCGTTAAATC |
| | From pJN105-esaR REV | GCCTGCCAGGCGGTAC |
| pJN105-esaR Δ168-171 | From pJN105-esaR FWD | TTAAATCAGAGCGCGG |
| | From pJN105-esaR REV | TTCACCTTCGGTGCC |
| pJN105-esaR Δ172-175 | From pJN105-esaR FWD | GCGGACAAAACGATATTTTCCTCG |
| | From pJN105-esaR REV | CGCCGGGGCTCGTTC |
| pJN105-esaR Δ176-179 | From pJN105-esaR FWD | ATATTTTCCTCGCGTG |
| | From pJN105-esaR REV | GCTCTGATTTAACGCC |
| pJN105-esaR Δ170-173 | From pJN105-esaR FWD | CAGAGCGCGGACAAAACG |
| | From pJN105-esaR REV | GGCTCGTTCACCTTCGGTG |
| pJN105-esaR Δ164-169 | From pJN105-esaR FWD | CCGGCGTTAAATCAGAGCG |
| | From pJN105-esaR REV | GCCTGCCAGGCGGTAC |
| pJN105-esaR Δ171-176 | From pJN105-esaR FWD | GACAAAACGATATTTTCCTCGC |
| | From pJN105-esaR REV | CGGGGCTCGTTCACCTT |
| pJN105-esaR Δ168-169, 171-176 | From pJN105-esaR FWD | CCGGACAAAACGATATTTTCCTCGCG |
| | From pJN105-esaR REV | TTCACCTTCGGTGCCTGC |
| pJN105-esaR Δ164-171 | From pJN105-esaR FWD | TTAAATCAGAGCGCGG |
| | From pJN105-esaR REV | GCCTGCCAGGCGGTAC |
| pJN105-esaR Δ164-175 | From pJN105-esaR FWD | GCGGACAAAACGATATTTTCCTCG |
| | From pJN105-esaR REV | GCCTGCCAGGCGGTAC |

| | | |
|---|---|---|
| pJN105-esaR scr 172-177 | From pJN105-esaR FWD | GCGAGCAATAAAACGATATTTTCCTCGCGTGAAAATGAGG |
| | From pJN105-esaR REV | TAACTGGTCCGCCGGGGCTCGTTCAC |
| pJN105-esaR ΔY159 | From pJN105-esaR FWD | CGCCTGGCAGGCACCGAA |
| | From pJN105-esaR REV | CATCTGCTCGTTAAAATCAATCAGCAGCATCTG |
| pJN105-esaR Δ159-162 | From pJN105-esaR FWD | GGCACCGAAGGTGAACGAG |
| | From pJN105-esaR REV | CATCTGCTCGTTAAAATCAATCAGC |
| pJN105-esaR Y159A | From pJN105-esaR FWD | CAGGCACCGAAGGTGAACGAG |
| | From pJN105-esaR REV | CCAGGCGCGCCATCTGCTC |
| pJN105-esaR Y159F | From pJN105-esaR FWD | CGAGCAGATGTTTCGCCTGGCAG |
| | From pJN105-esaR REV | TTAAAATCAATCAGCAGCATCTG |
| pJN105-esaR Y159Q | From pJN105-esaR FWD | CAGGCACCGAAGGTGAACGAG |
| | From pJN105-esaR REV | CCAGGCGCTGCATCTGCTC |
| pJN105-esaR G163A | From pJN105-esaR FWD | CACCTTCGGTGGCTGCCAGGCGG |
| | From pJN105-esaR REV | CCGCCTGGCAGCCACCGAAGGTG |
| pJN105-esaR T164A | From pJN105-esaR FWD | GTTCACCTTCGGCGCCTGCCAGGCG |
| | From pJN105-esaR REV | CGCCTGGCAGGCGCCGAAGGTGAAC |
| pJN105-esaR E165A | From pJN105-esaR FWD | GGCTCGTTCACCTGCGGTGCCTGCCAG |
| | From pJN105-esaR REV | CTGGCAGGCACCGCAGGTGAACGAGCC |
| pJN105-esaR G166A | From pJN105-esaR FWD | GGGGCTCGTTCAGCTTCGGTGCCTG |
| | From pJN105-esaR REV | CAGGCACCGAAGCTGAACGAGCCCC |
| pJN105-esaR E167A | From pJN105-esaR FWD | GCCGGGGCTCGTGCACCTTCGGTGC |

| | | |
|---|---|---|
| | From pJN105-esaR REV | GCACCGAAGGTGCACGAGCCCCGGC |
| pJN105-esaR R168A | From pJN105-esaR FWD | GCCGGGGCTGCTTCACCTTCGGTGCCTG |
| | From pJN105-esaR REV | CAGGCACCGAAGGTGAAGCAGCCCCGGC |
| pJN105-esaR P170A | From pJN105-esaR FWD | TCTGATTTAACGCCGCGGCTCGTTCACCTTC |
| | From pJN105-esaR REV | GAAGGTGAACGAGCCGCGGCGTTAAATCAGA |
| pJN105-esaR L172A | From pJN105-esaR FWD | CCGCGCTCTGATTTGCCGCCGGGGCTCGTT |
| | From pJN105-esaR REV | AACGAGCCCCGGCGGCAAATCAGAGCGCGG |
| pJN105-esaR N173A | From pJN105-esaR FWD | CCGCGCTCTGAGCTAACGCCGGGGCTCG |
| | From pJN105-esaR REV | CGAGCCCCGGCGTTAGCTCAGAGCGCGG |
| pJN105-esaR Q174A | From pJN105-esaR FWD | GTCCGCGCTCGCATTTAACGCCGGGGCTCG |
| | From pJN105-esaR REV | CGAGCCCCGGCGTTAAATGCGAGCGCGGAC |
| pJN105-esaR S175A | From pJN105-esaR FWD | TATCGTTTTGTCCGCGGCCTGATTTAACGCCGGG G |
| | From pJN105-esaR REV | CCCCGGCGTTAAATCAGGCCGCGGACAAAACGAT A |
| pJN105-esaR D177A | From pJN105-esaR FWD | GAGGAAAATATCGTTTTGGCCGCGCTCTGATTTAA CG |
| | From pJN105-esaR REV | CGTTAAATCAGAGCGCGGCCAAAACGATATTTTCC TC |
| pJN105-esaR K178A | From pJN105-esaR FWD | ACGCGAGGAAAATATCGTTGCGTCCGCGCTCTGA TTTAAC |
| | From pJN105-esaR REV | GTTAAATCAGAGCGCGGACGCAACGATATTTTCCT CGCGT |
| pJN105-esaR T179A | From pJN105-esaR FWD | CGAGGAAAATATCGCTTTGTCCGCGCTCTGATTTA AC |
| | From pJN105-esaR REV | GTTAAATCAGAGCGCGGACAAAGCGATATTTTCCT CG |
| pJN105-esaR I180A | From pJN105-esaR FWD | ATTTTCACGCGAGGAAAATGCCGTTTTGTCCGCGC TCTGA |
| | From pJN105-esaR REV | TCAGAGCGCGGACAAAACGGCATTTTCCTCGCGT GAAAAT |

| | | |
|---|---|---|
| pJN105-esaR F4A | From pJN105-esaR FWD | AACCAAACAATAACGGATACG |
| | From pJN105-esaR REV | TTCAAGGAACGCAGAGAACATG |
| pJN105-esaR F5A | From pJN105-esaR FWD | AACCAAACAATAACGGATACG |
| | From pJN105-esaR REV | TTCAAGCGCGAAAGAGAACATG |
| pJN105-mrtR +4G at 171 | From pJN105-mrtR FWD | GGTGGTGAGAACGATCCGGTCC |
| | From pJN105-mrtR REV | GCCACCCCCGTGCAACTCGTAAAC |
| pJN105-mrtR +4G at 175 | From pJN105-mrtR FWD | GGTGGTGTCCCCGCCCTTTCG |
| | From pJN105-mrtR REV | GCCACCCGGATCGTTCTCCCC |
| pJN105-mrtR +4A at 171 | From pJN105-mrtR FWD | Resulted from cloning of pJN105-MrtR +5A at 171 |
| | From pJN105-mrtR REV | |
| pJN105-mrtR +4A at 175 | From pJN105-mrtR FWD | GCGGCAGTCCCCGCCCTTTCG |
| | From pJN105-mrtR REV | CGCTGCCGGATCGTTCTCCCC |
| pJN105-mrtR +5A at 171 | From pJN105-mrtR FWD | CAGCGGCAGAGAACGATCCGGTCC |
| | From pJN105-mrtR REV | CCGCTGCCCCGTGCAACTCGTAAAC |
| pJN105-mrtR +5A at 175 | From pJN105-mrtR FWD | CAGCGGCAGTCCCCGCCCTTTCG |
| | From pJN105-mrtR REV | CCGCTGCCGGATCGTTCTCCCC |
| pJN105-mrtR Δ171-173 | From pJN105-mrtR FWD | GATCCGGTCCCCGCCCTT |
| | From pJN105-mrtR REV | GTGCAACTCGTAAACCGCTTTGC |
| pJN105-expR2 +5A at 171 | From pJN105-expR2 FWD | CAGCGGCAAGAAAACAAGGTGACAAAGAG |
| | From pJN105-expR2 REV | CCGCTGCGCTCTGCGGGCTTTG |
| pJN105-expR2 +5A at 176 | From pJN105-expR2 FWD | CAGCGGCAAAAGAGATTTTCTCCCAG |

| | | |
|---|---|---|
| | From pJN105-expR2 REV | CCGCTGCGTCACCTTGTTTTCTGC |
| pJN105-expR2 +5G at 171 | From pJN105-expR2 FWD | GTGGTGGCAGAAAACAAGGTGACAAAGAG |
| | From pJN105-expR2 REV | CGCCACCGCTCTGCGGGCTTTG |
| pJN105-expR2 +5G at 176 | From pJN105-expR2 FWD | GTGGTGGCAAAGAGATTTTCTCCCAG |
| | From pJN105-expR2 REV | CGCCACCGTCACCTTGTTTTCTGC |
| pJN105-expR2 +5: 172-176 | From pJN105-expR2 FWD | GGTGACAAAGAGATTTTCTCCCAGCG |
| | From pJN105-expR2 REV | TTGTTTTCTGTCACCTTGTTTTCTGCTCTG |
| pJN105-expR2 +10: 168-177 | From pJN105-expR2 FWD | AAGGAGACAAAAGCCCGCAGAGCAGAAAAC |
| | From pJN105-expR2 REV | GTTTACGACTTTGAGGTGATTGCGTCATCTCTTTG TAAAGCG |
| pJN105-expR2 +21:158-178 | From pJN105-expR2 FWD | ACGTAAGCAAGGAGACAAAGAGATCACCACGCTT TACAAAG |
| | From pJN105-expR2 REV | GACTGCGGTGATTGTGTCATTTCCTTATACAAAGT AGTAATTTTCTCATGAACGGCAATG |
| pJN105-expR2 Δ169-172 | From pJN105-expR2 FWD | AAACAAGGTGACAAAGAG |
| | From pJN105-expR2 REV | GCTTTGCGTCATCTCT |
| pJN105-expR2 Δ173-176 | From pJN105-expR2 FWD | AAAGAGATTTTCTCCCAGCGC |
| | From pJN105-expR2 REV | TCTGCTCTGCGGGCTTTG |
| pJN105-expR2 Δ170-175 | From pJN105-expR2 FWD | GACAAAGAGATTTTCTCCCAGCG |
| | From pJN105-expR2 REV | CGGGCTTTGCGTCATCTCTTTG |
| pJN105-expR2 Δ169-176 | From pJN105-expR2 FWD | AAAGAGATTTTCTCCCAGCG |
| | From pJN105-expR2 REV | GCTTTGCGTCATCTCTTTGTAAAG |
| pJN105-expR2 K177A | From pJN105-expR2 FWD | ACAAGGTGACGCGGAGATTTTCTCCCAG |
| | From pJN105-expR2 REV | TTTCTGCTCTGCGGG |

| | | |
|---|---|---|
| pJN105-expR2 E178A | From pJN105-expR2 FWD | AGGTGACAAAGCGATTTTCTCCC |
| | From pJN105-expR2 REV | TGTTTTCTGCTCTGCG |
| pJN105-expR2 F4A | From pJN105-expR2 FWD | CAATGAAATTATCAATAACACGAT |
| | From pJN105-expR2 REV | TCAGAGCACGCTACAGACAT |
| pJN105-expR2 ∆2-175 | From pJN105-expR2 FWD | GACAAAGAGATTTTCTCCC |
| | From pJN105-expR2 REV | CATGAATTCGCTAGCC |
| pJN105-FLAG-expR2 ∆2-177 | From pJN105-expR2 FWD | GATGATGATAAAGAGATTTTCTCCCAGC |
| | From pJN105-expR2 REV | ATCTTTATAATCCATGAATTCGCTAGCC |

**<u>Raw western blot images.</u>**

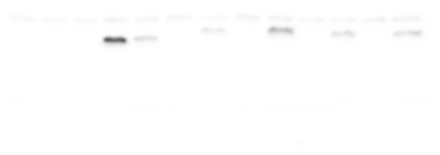

Lanes 3-13 correspond to FLAG blot shown on left in **Figure S5E**.

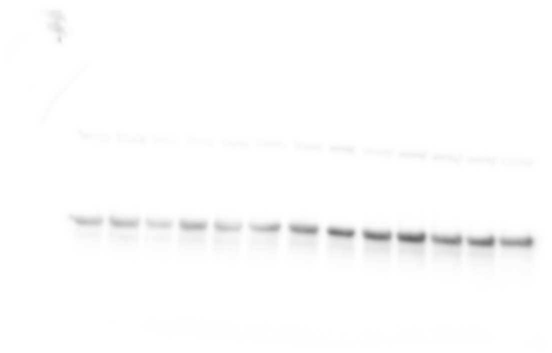

Lanes 3-13 correspond to GAPDH blot shown on left in **Figure S5E**.

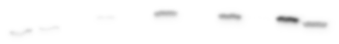

Lanes 1-11 correspond to FLAG blot shown on left of **Figure S11A**.

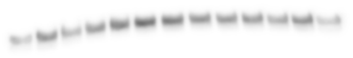

Lanes 1-11 correspond to GAPDH blot shown on left of **Figure S11A**.

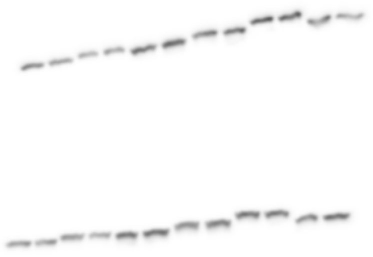

Two gels with separate biological replicates were transferred to same membrane and imaged together. Top corresponds to FLAG for **Figure S11B**. Bottom corresponds to FLAG for **Figure S8B**.

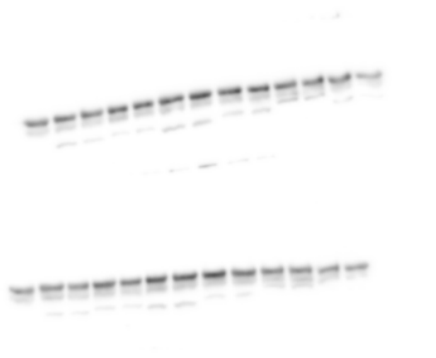

Two gels with separate biological replicates transferred to same membrane and imaged together. Top corresponds to GAPDH for **Figure S11B**. Bottom corresponds to GAPDH for **Figure S8B**. The corresponding FLAG blot shown above was treated with 10% acetic acid for 30 min at 37 °C to inactivate HRP, then washed in water and TBST and re-stained for GAPDH.

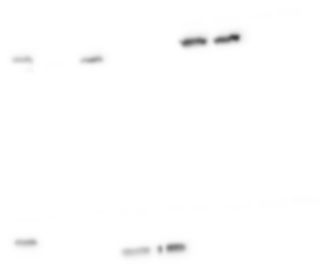

Top row, lanes 4-11 correspond to FLAG blot shown in middle of **Figure S5E**. Bottom row, lanes 4-13 correspond to FLAG blot shown on right in **Figure S5E**.

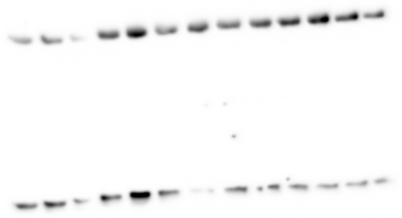

Top row, lanes 4-11 correspond to GAPDH blot shown in middle of **Figure S5E**. Bottom row, lanes 4-13 correspond to GAPDH blot shown on right in **Figure S5E**. The corresponding FLAG blot shown above was treated with 10% acetic acid for 30 min at 37 °C to inactivate HRP, then washed in water and TBST and stained for GAPDH.

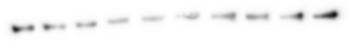

GAPDH blot using the same samples as the GAPDH blot shown on right of **Figure S5E** and directly above, which showed uneven staining.

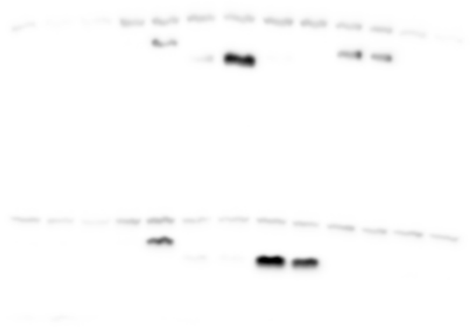

Top row, lanes 4-11 correspond to FLAG blot shown in middle of Figure **S11A**. Bottom row, lanes 4-13 correspond to FLAG blot shown on right of Figure **S11A**.

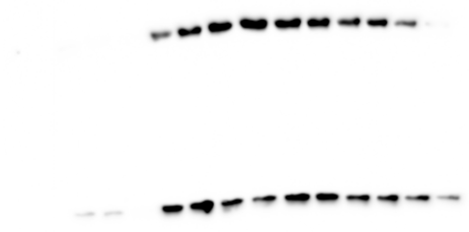

Top row corresponds to FLAG blot shown in middle of **Figure S11A**. Bottom row, lanes 4-13 correspond to FLAG blot shown on right of **Figure S11A**. Lanes 1-3 are very faint in both blots. The corresponding FLAG blot shown directly above was treated with 10% acetic acid for 30 min at 37 °C to inactivate HRP, then washed in water and TBST and stained for GAPDH.

**References.**
